# Homeostasis of a representational map in the neocortex

**DOI:** 10.1101/2023.06.13.544358

**Authors:** Takahiro Noda, Eike Kienle, Jens-Bastian Eppler, Dominik F. Aschauer, Matthias Kaschube, Yonatan Loewenstein, Simon Rumpel

**Affiliations:** Institute of Physiology, Focus Program Translational Neurosciences, University Medical Center, Johannes Gutenberg University-Mainz, Mainz, Germany; Frankfurt Institute for Advanced Studies and Institute for Computer Science, Goethe University Frankfurt, Frankfurt, Germany; The Edmond & Lily Safra Center for Brain Sciences, Department of Cognitive and Brain Sciences, The Alexander Silberman Institute of Life Sciences and The Federmann Center for the Study of Rationality, The Hebrew University, Jerusalem, Israel

## Abstract

Cortical function in general and the processing of sensory stimuli in particular are remarkably robust against the continuous loss of neurons during aging, and even the accelerated loss during prodromal stages of neurodegeneration^1,2^. Population activity of neurons in sensory cortices represents the environment in form of a map, which is structured in an informative way for guiding behavior. Here, we used the mouse auditory cortex as a model and tested in how far the structure of the representational map is protected by homeostatic network mechanisms against the removal of neurons. We combined longitudinal two-photon calcium imaging of population responses evoked by a diverse set of sound stimuli with a targeted microablation of functionally characterized neurons. Unilateral microablation of 30 - 40 selected highly sound-responsive neurons in layer 2/3 led to a temporary disturbance of the representational map in the spared population that, however, recovered in subsequent days. At the level of individual neurons, we observed that the recovery of the spared network was predominantly driven by neurons unresponsive to the sounds before microablation which strengthened the correlation structure of the local network after gaining responsiveness. In contrast, selective microablation of inhibitory neurons induced a prolonged disturbance of the representational map that was primarily characterized by a destabilization of sound responses across trials. Together, our findings provide a link between the tuning and plasticity of individual neurons and the structure of a representational map at the population level which reveals homeostatic network mechanisms safeguarding sensory processing in neocortical circuits.

## Introduction

Sensory information is represented in the cortex in form of a representational map in which various properties of sensory entities are encoded as distances between sensory-driven neuronal activity patterns^3–5^. The representational map offers a general framework to describe neuronal population activity along a sensory-motor transformation. In sensory areas of the cortex, the structure of the map is dominated by perceptual aspects of the stimuli. Experimentally, the structure can be estimated by a similarity measure between population responses to different sensory stimuli^6^ and has been utilized as a readout of neuronal function in various sensory domains in order to characterize perception-related behavior^7–10^, including the auditory modality^11,12^.

A representational map is formed during development and its stability is continuously challenged during adulthood. At the level of sensory evoked activity, a continuous reformatting of tuning properties of individual neurons has been described in recent year as representational drift^9,13^. Moreover, a continuous drop in and drop out of individual neurons from the population response has been demonstrated in various brain regions^9,12–16^. A possible mechanism for these functional changes in tuning may lie in the substantial plastic changes in the synaptic structure of neocortical circuits that can be observed under environmentally and behaviorally stable conditions^17–19^.

Apart from changes in the synaptic connections, representing the edges in a network, also the loss of neurons, i.e., the nodes in the network, challenges the stability of representational maps. Specifically, a continuous loss of neurons has been observed during aging^20–22^. In the context of neurodegenerative diseases this loss is considerably increased. Surprisingly, however, during healthy aging as well as even advanced prodromal neurodegenerative stages, a remarkable robustness in cognitive functions has been described until a tipping point is reached^1,2,23^.

What processes could endow cortical networks with robustness against these challenges? On the one hand, specific activity patterns could rely on a highly redundant connectivity structure, such that significant changes in the network do not alter activity patterns^24–26^. On the other hand, specific manipulations on a small number of neurons can nevertheless have a big effect on activity patterns and behavior^27–30^. This hints that neuronal activity patterns may not be intrinsically robust, but be stabilized by dynamic feedback mechanisms.

In the body various important physiological parameters such as blood pressure or body temperature are challenged by various short- and long-term disturbances, but kept within a functional range due to homeostatic regulation^31^. In neural systems, several examples of homeostasis have been demonstrated^32–34^, including firing rate homeostasis at the level of individual neurons^35–37^ and network homeostasis of pacemaker patterns in the gastropyloric ganglion in crayfish^38^. However, in how far homeostatic mechanisms shape and safeguard population dynamics a cortical circuit is not well understood. Here, we probed the robustness of a representational map against the experimental removal of functionally selected neurons. We found that the representational map is transiently impaired, but recovers over the period of few days. Analyzing the underlying changes in single-neuron tuning properties, we find evidence that cortical circuits actively employ various mechanisms to mediate the homeostatic compensation of this loss.

## Results

### Long-term stability of a representational map despite volatility in single-neuron tuning

To monitor the long-term dynamics of sound representations in the mouse auditory cortex, we co-injected two recombinant adeno-associated virus vectors to express the genetically encoded calcium indicator, GCaMP6m as a neuronal activity marker^39^ and the fusion protein H2B::mCherry to highlight the nuclei of transduced neurons for high-fidelity image registration and analysis (Fig. 1a,b, see Methods)^12,40^. We performed intrinsic signal imaging in response to a set of pure-tone stimuli with different frequencies to functionally identify the tonotopic structure of the auditory cortex to guide subsequent two-photon imaging (Extended Data Fig. 1a). For calcium imaging experiments in awake, head-fixed, passively listening mice, we used a stimulus set of brief (50 – 70 ms) sounds consisting of 19 sinusoidal pure tones (PT) and 15 complex sounds (CS) characterized by temporally modulated power in multiple frequency bands (34 stimuli in total, 10 repetitions each, presented in a pseudorandom order, Fig. 1c) delivered in free field using a calibrated speaker at 70 dB sound pressure level^12^. Mice were extensively habituated to head fixation and pre-exposed to the set of sound stimuli for at least one week to perform imaging in familiar behavioral and experimental conditions^12,41^. In a given mouse, we consecutively imaged neuronal activities in 6-8 Fields Of Views (FOVs) vertically aligned within layer 2/3 (Extended Data Fig. 1b). When assessing the trial-averaged calcium responses to the set of sound stimuli in a FOV (∼140 - 180 neurons), we observed diverse tuning (Fig. 1c) in a smaller fraction of neurons (∼10 - 14%), whereas the majority of neurons were unresponsive in a given imaging session. The neuronal populations were tracked and re-imaged for 6 time points at a 2-day interval and a last imaging time point after a 4-day interval (9,451 neurons, 59 FOVs, 9 mice; Extended Data Fig. 1c).

**Fig. 1.**
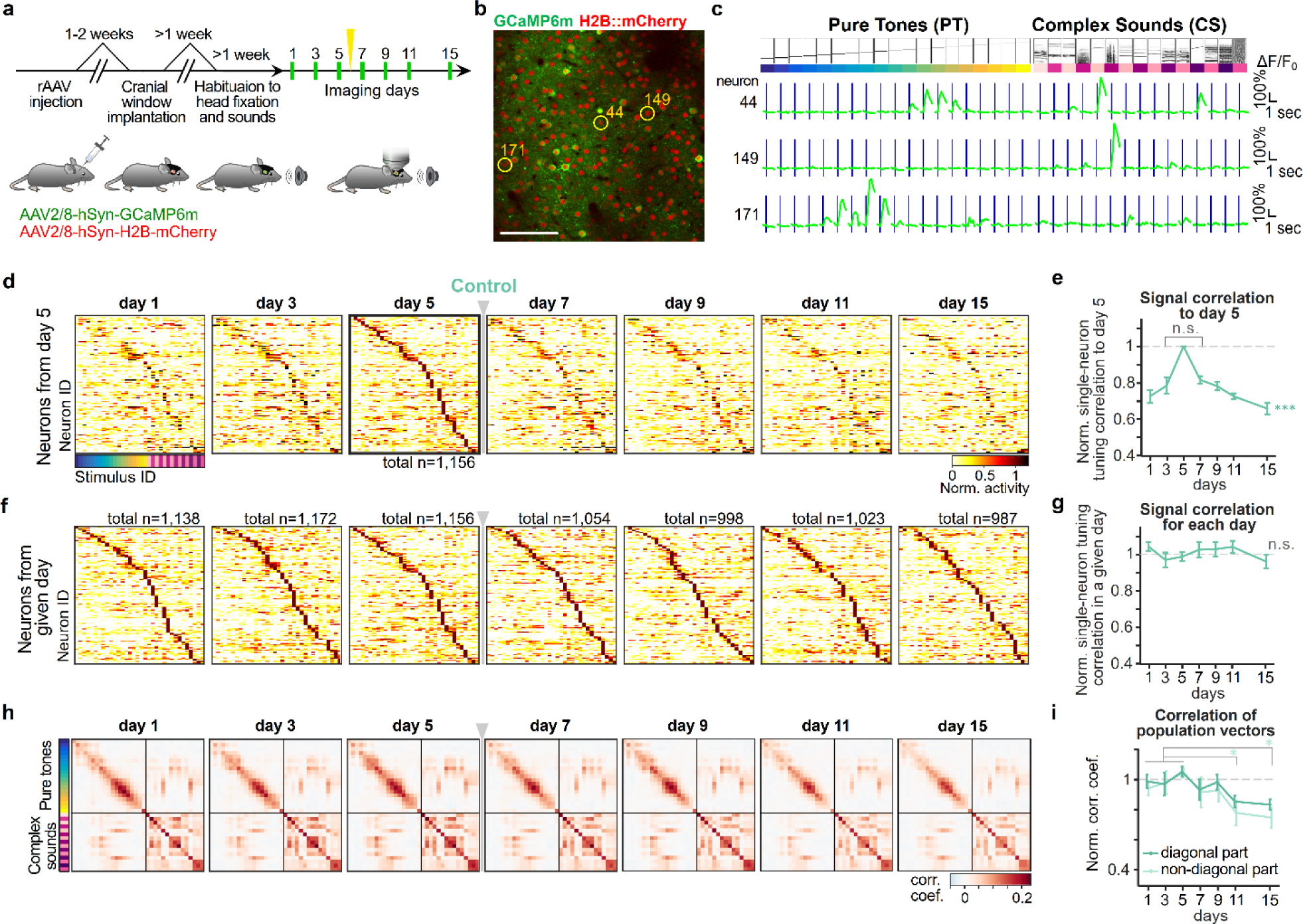
Neurons with unstable tuning properties form a stable representational map. **a.** Schematic of the experiment for longitudinal Ca^2+^ imaging in the mouse auditory cortex. On day 6 a sham ablation was applied in the control cohort as a reference for other experimental cohorts described later (yellow arrow, see Methods). **b**. Example imaging plane showing broad expression of GCaMP6m and H2B::mCherry in layer 2/3 neurons. **c**. Exemplary responses to auditory stimuli (19 pure tones ranging between 2 - 45 kHz and 15 complex sounds) of the neurons indicated in **b**. **d**. Normalized response profiles for each imaging day for the subset of neurons categorized as sound responsive and sorted by stimulus with highest response amplitude on day 5. The gray arrow and bar indicate the day when the sham procedure was applied in this control cohort. **e**. Normalized correlation of single-neuron stimulus tuning between day 5 and the other day, for the neurons shown in **d**. **f**. Same as **d**, however, considering the neurons that were categorized as sound responsive and sorted by stimulus with highest response amplitude on each given day. **g**. Same as **e**, but baseline-normalized single-neuron correlation for each day. The baseline correlation was defined as the average of the three baseline days from day 1 to day 5. **h**. Similarity matrix of population response vectors for all stimuli averaged across all FOVs for each day. **i**. Normalized correlations averaged across diagonal elements (dark green) and all off-diagonal elements (light green) in the similarity matrices shown in **h** normalized to the average of baseline days.

Consistent with previous reports^12,42^, we observed a substantial change in responsiveness and tuning in individual neurons during the time course of the experiment in our dataset. Whereas the tuning curves of neurons categorized as responsive to any of the 34 stimuli on day 5 tiled the set of stimuli, the responses in the same set of neurons on earlier or later imaging time points showed a progressively increasing degree of redistribution (Fig. 1d). To quantify the changes in tuning in an individual neuron, we calculated the Pearson correlation of pairs of tuning curves constructed by randomly splitting the trials into two halves and averaging the responses. For the Pearson correlation on day 5 we used all trials recorded on that day, for the other intervals, one half of the trials stemmed from day 5 and the other half from the other imaging days. The correlation of tuning curves showed a monotonous and symmetric decay from imaging day 5 to increasing intervals (Fig. 1e; one-way ANOVA across days, F(6,56) = 7.77, p = 4.21×10^-6^; correlation between day 3 and day 7, between day 1 and day 9, paired t-test p = 0.69, p = 0.12, respectively).

The volatility in tuning of individual neurons, however, was balanced at the population level, as observed in previous reports^9,12–14^. When identifying the neurons that were significantly responsive on a given day, their number was comparable throughout the experiment and their sorted response profiles were highly similar across days (Fig. 1f and Extended Data Fig. 1d). Also, the Pearson correlation of average tuning curves from the halves of trials obtained on a given day was largely constant, indicating a stable level of response reliability across trials (Fig. 1g; one-way ANOVA, F(6, 56) = 0.55, p = 0.77). Similarly, the ability to decode the identity of a pair of sound stimuli from neural population activity gradually decreased when single-trial neural activity patterns were tested at increasing intervals from the day when activity patterns were recorded to train the classifier. However, when the decoder was trained and tested with activity patterns recorded at the same day, the performance was largely stable across imaging days (Extended Data Fig. 1e).

We next assessed how sounds were represented at the population level in the form of a representational map using an approach analogous to a Representational Similarity Analysis (RSA)^6^. In a given FOV, we created single-trial population response vectors by averaging the amplitude of the calcium signals for each neuron within a 400ms time window after each sound presentation. We constructed a similarity matrix by calculating the average Pearson correlation of the response vectors for all pairwise combinations of trials evoked by the same stimulus for the elements along the diagonal and for all combinations of response vectors from a given pair of stimuli for the off-diagonal elements^12^. To obtain a global measure of pairwise similarity of the population responses, we averaged the similarity matrices of all FOVs for each day (Fig. 1h). The resulting grand-averaged similarity matrix of the neuronal activity patterns evoked by the set of sound stimuli provides a pairwise measure of perceptual similarity in mice allowing to predict the level of generalization in various behavioral paradigms (Extended Data Fig. 2)^11,12^. Therefore, the similarity matrix provides a useful estimate of the internal representational map reflecting behaviorally relevant relational information of sounds.

Over the time course of the experiment, the structure of the similarity matrices showed a remarkably high level of similarity (Fig. 1h). Correspondingly, the average correlation in the matrices along the diagonal and off-diagonal remained at a level comparable to the first five days of the experiment (Fig. 1i), although we detected a slight and gradual decay (Fig. 1i; two-sample t-test between baseline days vs. days after ablation with False Discovery Rate (FDR) correction. diagonal part: p = 0.830, p > 0.9, p = 0.0585, p = 0. 0585; non-diagonal part: p = 0.835, p = 0.835, p = 0.0293, p = 0.0214, for day 7, 9, 11, 15, respectively). This indicated that the pairwise representational similarity or dissimilarity of the various pairs of sounds is largely preserved despite the substantial drop-in and drop-out of individual neurons from the population response (Fig. 1d).

Together, these observations showed that the global structure of the representational map is stably maintained despite a substantial drift in single neuron tuning and a progressive decorrelation of population response vectors for a given stimulus over days, suggesting that plasticity mechanisms constantly preserve higher-order statistics of the population responses.

### The loss of sound responsive neurons leads to a transient disturbance of the representational map

In order to reveal network mechanisms that support the dynamic maintenance of the representational map, we next studied the effects induced by a permanent removal of individual neurons from the local microcircuit. We considered three major scenarios of how the map could be affected: i) A permanent removal of neurons could lead to a permanent impairment of the map. ii) The map could recover from the effects induced by the removal of neurons by a gradual wash-out mediated by the ongoing volatility of tuning observed during baseline. iii) The map could recover from the effects induced by the removal of neurons by the engagement of specific homeostatic processes.

Towards this end, we used targeted laser microablation^29,43,44^ (Fig. 2a, Extended Data Fig. 3a-c) yielding a highly specific and permanent removal of individual, functionally characterized neurons (Fig. 2b, Extended Data Fig. 3c-i, see Methods).

**Fig 2.**
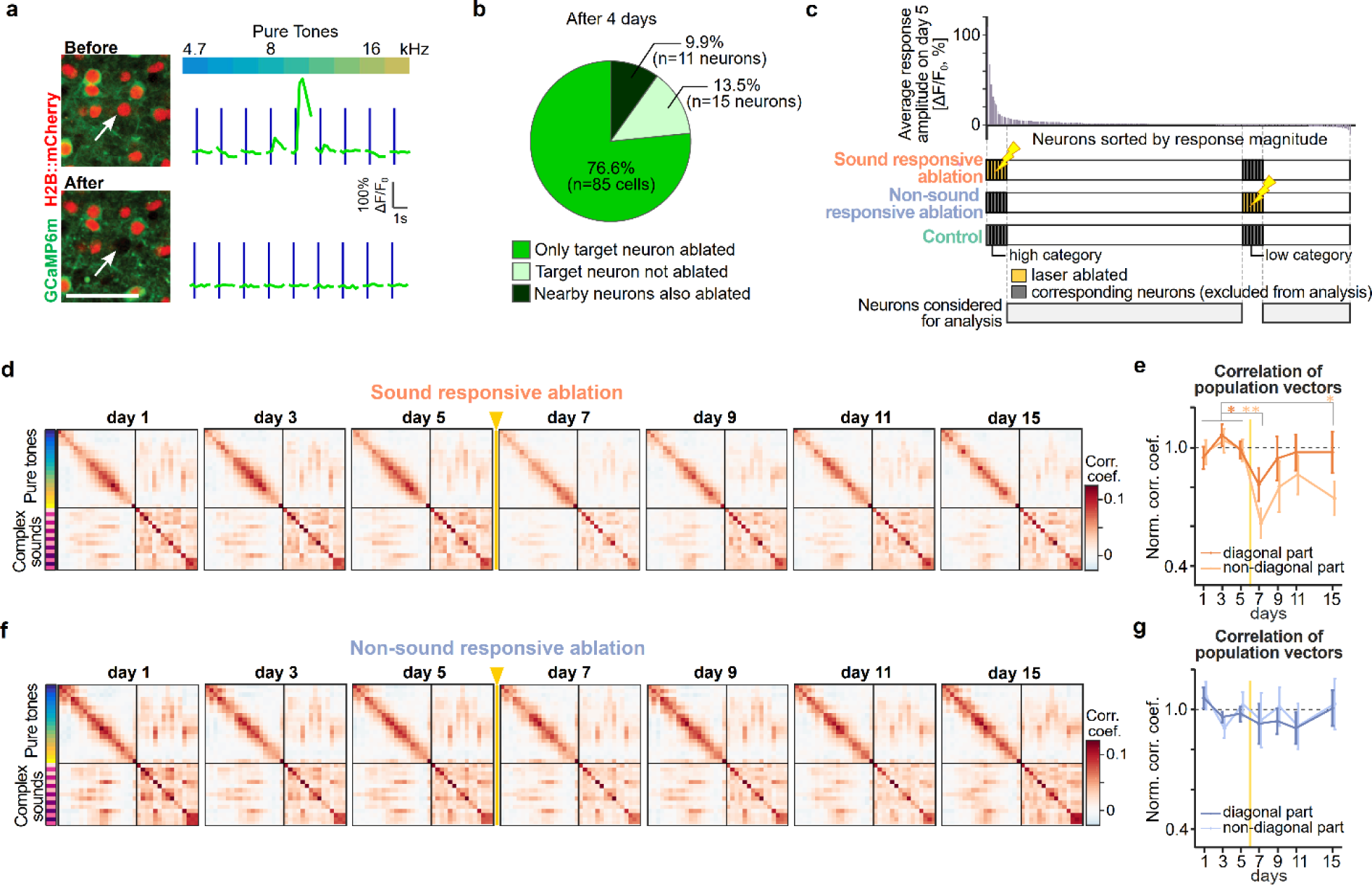
Microablation-induced disturbance and recovery of the representational map. **a.** Representative example of two-photon microablation of a functionally characterized neuron with pure tone tuning. Upper inset: Before ablation, Lower inset: 2 days after ablation. Microablated neurons showed permanent loss of nucleus signal and sound-evoked signal. Scale: 50 μm. **b**. Specificity of single-neuron ablation protocol obtained in a pilot experiment in two mice. **c**. Target neurons for microablation. Top: Sorted sound-evoked amplitudes before microablation in a FOV. For the group of sound responsive microablation, 4 - 8 neurons of highly responsive neurons were targeted per FOV, in 6 - 8 FOVs vertically aligned in layers 2/3 in a given mouse (see Methods). For the non-sound responsive microablation group, a similar number of non-responsive neurons were targeted. The control group had no microablation on day 6. Bottom: Experimentally targeted or corresponding responsive and non-responsive neurons were excluded, and further data analysis was conducted on remaining spared neurons (gray bars). **d**. Similarity matrix of population vectors for all stimuli averaged across all FOVs for each day in sound responsive microablation cohort, analogous to Fig. 1 **h**. **e**. Baseline-normalized correlations averaged across diagonal elements (dark orange) and off-diagonal elements (light orange) in the similarity matrices in **d**. **f**. Same as **d**, but similarity matrix in non-sound responsive ablation cohort. **g**. Same as **e**, but baseline-normalized correlations averaged across diagonal elements (dark blue) and off-diagonal elements (light blue) in the similarity matrices in **f**.

To probe the robustness of the representational map, we started by obtaining the distribution of sound-evoked response amplitudes across neurons in each FOV during baseline considering day 3 and day 5 before the day of microablation on day 6 (Fig. 2c, see methods) and devised two cohorts of mice, one in which particularly sound-responsive neurons were targeted for microablation (10 mice), and a second cohort in which non-sound responsive neurons were targeted (10 mice) (Extended Data Fig. 4a,b). In each cohort 3 - 7 neurons were microablated per FOV resulting in a largely uniform distribution along cortical depth within layer 2/3 (Extended Data Fig. 4c). In total, we microablated 34.1 ± 2.8 neurons on average per mouse in the sound-responsive cohort and 35.3 ± 2.4 neurons per mouse in the non-sound responsive cohort, which corresponded to ∼3% of imaged and analyzed neurons.

This number of individually microablated neurons reflects the maximum achievable within a single session given our procedure. To assess the effects of microablation on the spared network, we excluded the actually microablated neurons in the sound responsive and non-sound responsive cohort from analysis for all time points before and after microablation. In addition, for each sound responsive and non-sound responsive cohort we excluded a comparable number of neurons with similarly high responses and low responses of the distribution of sound-evoked response amplitudes (“high category neurons” and “low category neurons”, respectively), that corresponded to the ablated neurons in the other cohorts (Fig. 2c). In the control group, both “high category neurons” and “low category neurons” were similarly excluded from the analysis. Overall, we considered 10,078 neurons in 69 FOVs (10 mice) in the sound-responsive cohort, 10,604 neurons in 73 FOVs (10 mice) in the non-sound responsive cohort and 8,878 neurons in 59 FOVs (9 mice) in the control group allowing us to focus our analysis on the effects of microablation in the part of the neuronal network that was not directly targeted and to provide an unbiased comparison of sound-evoked responses between the different experimental cohorts (Extended Data Fig. 4d-j).

To assess the sensitivity of the representational map against the loss of a few tens of neurons, we again constructed averaged similarity matrices for each time point throughout the experiment. During baseline, we observed a largely stable structure of the representational map in the sound-responsive cohort as well as in the non-sound responsive cohort, that was similar to the control group shown in Fig. 1h,j (Fig. 2d-g and Extended Data Fig. 5a). Remarkably, in the cohort where particularly sound responsive neurons were microablated, we observed that similarity matrix showed a substantially lower contrast, i.e. lower absolute values of correlations, on the day after microablation, but showed a recovery during subsequent days regaining a structure that was again comparable to the one observed during baseline (Fig. 2d). This was also reflected when quantifying the baseline-normalized average correlation along the diagonal, which manifests the reliability of the population response to repeated presentations of the same stimulus (Fig. 2e; two-sample t-test of normalized average correlation between baseline days vs. days after ablation with FDR correction, in diagonal elements: p = 0.0436, p = 0.957, p = 0.957, p = 0.957 for day 7, 9, 11, 15, respectively; one-way ANOVA of normalized average correlation across days in diagonal elements: F(6,63) = 0.91, p = 0.49). This was generally also the case for the off-diagonal correlations, which manifests the pairwise similarities of the population responses elicited by two different stimuli, except for day 15 where the correlation did not fully recovered but reached to the level comparable with that of the control cohort (Fig. 2e; two-sample t-test for off-diagonal elements: p = 2.06×10^-5^, p = 0.239, p = 0.239, p = 0.0144 for day 7, 9, 11, 15, respectively; one-way ANOVA for off-diagonal elements: F(6,63) = 2.25, p = 0.049; Extended Data Fig. 5b-d).

In contrast, in the cohort of mice in which non-sound responsive neurons were mircoablated, the structure of the representational map was largely stable throughout the experiment (Fig. 2f,g; two-sample t-test of normalized average correlation between baseline days vs. days after ablation in diagonal elements: p > 0.72 for all post-ablation days; off-diagonal elements: p > 0.83 for all post-ablation days, One-way ANOVA of normalized average correlation across days in diagonal elements: F(6,63) = 0.68, p = 0.67; off-diagonal elements: F(6,63) = 0.58, p = 0.75; Extended Data Fig. 5 b-d).

In summary, our analyses showed that the selective removal of 30-40 highly sound-responsive neurons was sufficient to introduce a transient disturbance of the representational map formed by the spared neurons in the local network in layers 2/3, that, however, showed a recovery in its structure comparable to baseline within three days after microablation.

### Changes in single-neuron response properties underlying the plasticity of the representational map

The representational map is shaped by the collective statistics of the tuning properties of the individual neurons in the network^45^. To shed light on connection between neuronal activity and the representational geometry, we next focused our analyses on the changes of responses in individual neurons. The transient decrease in the diagonal entries of the similarity matrices constructed from single-trial population response vectors, indicated that the reliability of the sound-evoked population response may be destabilized in the early days following the microablation of sound responsive neurons. Using the correlation of tuning curves constructed from subsampled trials from an individual neuron as a measure of response reliability across trials (see Methods), we observed a transient decrease after microablation in the sound responsive ablation cohort (Fig. 3a; two-sample t-test of normalized reliability between baseline days vs. days after ablation with FDR correction, sound responsive cohort: p = 0.034 on day 7, p > 0.6 for the other post-ablation days, i.e., day 9, 11, 15; permutation test across groups on day 7: sound responsive cohort: p = 0.045; non-sound responsive cohort and control: p > 0.62, see Methods). The response reliability largely remained stable for the other experimental cohorts (Fig. 3a; two-sample t-test between baseline days vs. the other days after ablation with FDR correction, non-sound responsive cohort: p = 0.504, p = 0.276, p = 0.276, p = 0.526 for day 7, 9, 11, 15, respectively; control: p > 0.6 for all post-ablation days).

**Fig. 3.**
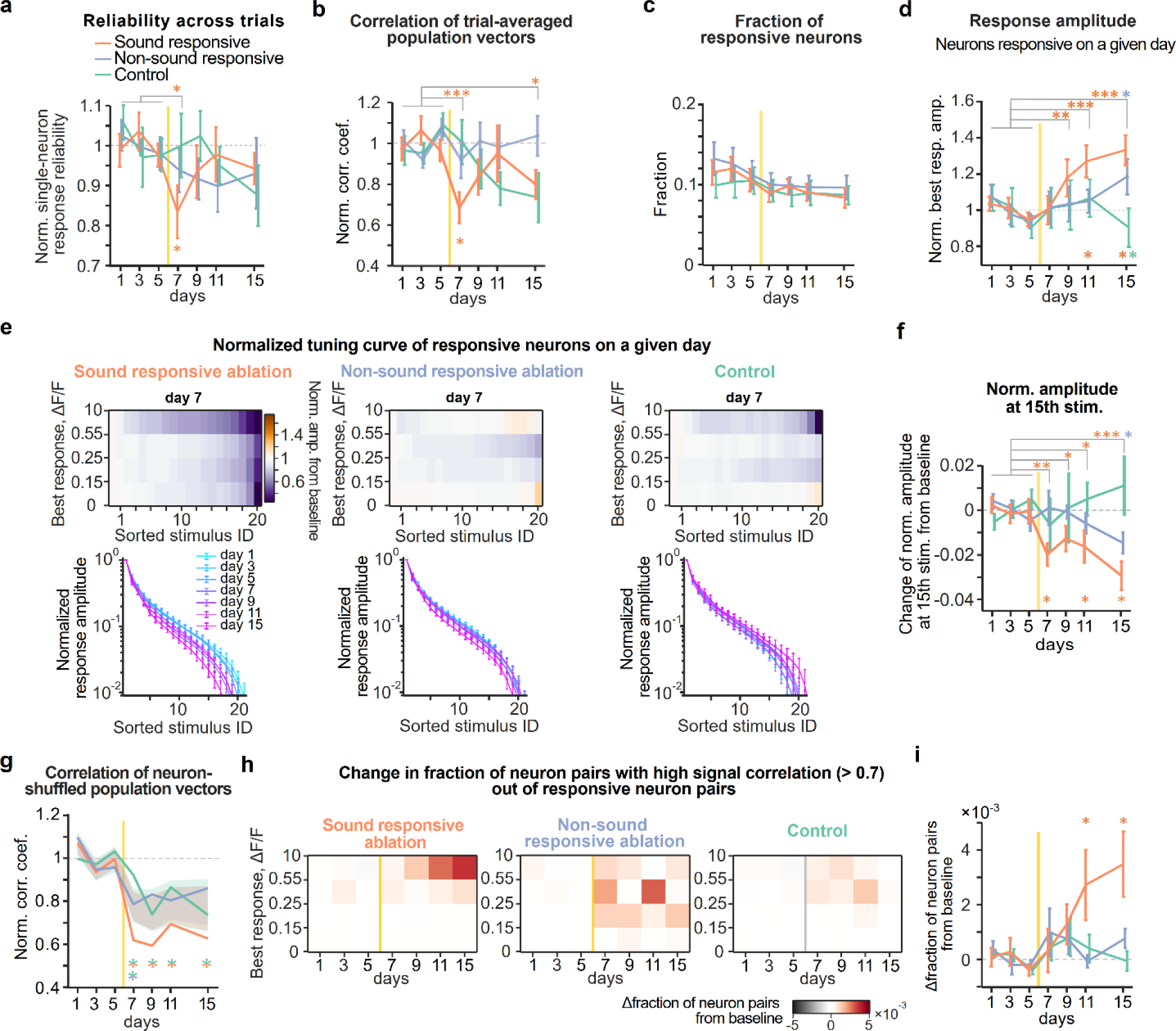
Microablation induces changes in single-neuron response properties at multiple timescales. **a.** Baseline-normalized response reliability across trials of single neurons averaged across all neurons on each given day (mean ± s.e.m. across mice. Orange, blue, and green lines indicate sound responsive ablation, non-sound responsive ablation and control group, respectively). **b**. Baseline-normalized correlations averaged across off-diagonal elements in the similarity matrix constructed from trial-averaged population response vectors. **c**. Fraction of sound-responsive neurons over the imaging days. **d**. Baseline-normalized best response amplitude of neurons responsive for each day. The best response amplitude is the response amplitude of the stimulus evoking the largest significant response. **e**. Normalized tuning curve of responsive neurons. For each neuron stimulus responses were sorted by magnitude, normalized to the maximum and then averaged across neurons within one of four categories according to their best response amplitude. Top: For each of the four best response categories, the change in the amplitude for the sorted stimuli on day 7 compared to baseline is color coded. Note sharpening of tuning in the sound responsive cohort for neurons with large response amplitudes. Bottom: normalized tuning curves for neurons of the large best response category across days. Left, middle and right columns are from sound responsive ablation, non-sound responsive ablation and control cohort, respectively. **f**. Change of normalized response amplitudes from baseline at 15^th^ largest stimulus index in the tuning curve in **e** over days. **g**. Average baseline-normalized off-diagonal correlations from similarity matrices constructed from population response vectors shuffled across FOVs. Shaded areas indicate 95% confidence interval after shuffling neurons across experimental cohorts (see Methods). Note that apparent recovery of off-diagonal correlations is no longer observed, suggesting changes in the local correlation structure occurring during later days after microablation. **h**. Color map displaying the change in the fraction of pairs among responsive neurons showing a high signal correlation (> 0.7). Neuron pairs were categorized according to their best response amplitudes (averaged over the two neurons). **i**. Change in fraction of neuron pairs with high signal correlation from baseline at the largest best response category in **h**. Note that the maximal change in fraction corresponds to a threefold increase of neuron pairs with high signal correlations compared to baseline.

In an attempt to delineate the effects induced by the lower reliability in sound-evoked responses in individual neurons from additional factors that could contribute to the temporary disturbance in the similarity matrices, we averaged the population vectors over the trials elicited by the same stimulus and constructed similarity matrices from the Pearson correlations between these trial-averaged population vectors (Extended Data Fig. 5e-g). By construction, the diagonal elements of these matrices are equal to 1 and cannot be affected by microablation. Despite averaging across trials, we observed a significant reduction in off-diagonal correlations on day 7 and subsequent recovery during later days in the sound responsive ablation cohort, although on day 15 the correlation did not fully recover to the baseline but reached a level comparable to the control cohort (Fig. 3b; two-sample t-test of normalized correlation coefficient between baseline days vs. days after ablation, sound responsive cohort: p = 7.90×10^-4^, p = 0.081, p > 0.60, p = 0.040 for day 7, 9, 11, 15, respectively; non-sound responsive cohort: p > 0.60 for all post-ablation days; control: p > 0.60, p = 0.48, p = 0.051, p = 0.051 for day 7, 9, 11, 15, respectively; permutation test across groups, sound responsive cohort: p < 0.005, p = 0.15, p = 0.71, p = 0.305 for day 7, 9, 11, 15; non-sound responsive cohort: p > 0.78 for all post-ablation days; control: p = 0.95, p = 0.37, p = 0.10, p = 0.065 for day 7, 9, 11, 15, respectively; Extended Data Fig. 5e-g). This analysis suggested that, apart from the changes in response reliability across trials, additional changes in the tuning of neurons contribute to the remodeling of the representational maps.

To identify what changes in tuning of individual neurons may be involved, we next investigated several major single-neuron response properties. When categorizing neurons in sound-responsive or non-sound responsive, based on a significant response to at least one stimulus (see Methods), we did not observe a significant change in the overall fraction of sound responsive neurons (Fig. 3c; one-way ANOVA across days, sound responsive cohort: F(6,63) = 1.47, p = 0.20; non-sound responsive cohort: F(6,63) = 0.80, p = 0.57; control: F(6,56) = 0.26, p = 0.95).

When analyzing the average response amplitude across trials elicited by the stimulus giving rise to the maximal response (best response amplitude), we observed an increase in the cohort of mice in which sound responsive neurons were microablated (Fig. 3d). Importantly, this effect on best response amplitudes increase became overt only several days after the microablation and therefore could not explain the disturbance of the representational map observed on the day after the microablation and rather may contribute to the recovery of the representational map (Fig. 3d; two-sample t-test between baseline days and days after ablation with adjusted p-values by FDR method: p = 0.977, p = 0.00452, p = 1.77×10^-5^, p = 3.71×10^-7^ for day 7, 9, 11, 15; non-sound responsive cohort: p = 0.835, p = 0.595, p = 0.595, p = 0.0191 for day 7, 9, 11, 15, respectively; control: p > 0.824 for all post-ablation days; permutation test across groups, sound responsive cohort: p = 0.565, p = 0.11, p = 0.005, p = 0.005 for day 7, 9, 11, 15; non-sound responsive cohort: p = 0.49, p = 0.70, p = 0.925, p = 0.315 for day 7, 9, 11, 15; control: p = 0.475, p = 0.80, p = 0.82, p = 0.995 for day 7, 9, 11, 15, respectively).

As tuning width is an important parameter determining the structure of representations at the population level, we examined the tuning width in neurons categorized as responsive, as broader tuning in individual neurons can increase the correlation between two population responses to different stimuli. In the sound responsive ablation cohort, but not for the other cohorts, we observed that the tuning width was reduced the day after microablation (Fig. 3e). This effect was particularly pronounced in neurons with large response amplitudes (Fig. 3e,f; two-sample t-test between baseline days and days after ablation with FDR correction at 15^th^ stimulus index, sound responsive cohort on day 7: p = 0.00189). We then assessed if the reduction in tuning width is sufficient to explain the transient disturbance of the similarity matrices constructed from trial-averaged population response vectors. When artificially narrowing the tuning curves of neurons recorded during baseline to a similar degree as seen after microablation, we reproduced the reduction of off-diagonal correlations in the similarity matrices (Extended Data Fig. 5h). Thus, the reduction of tuning width, together with a decreased reliability across trials in sound responses, could mediate the transient disturbance of the representational map. However, given that the narrowing of tuning persisted throughout the further days of the experiment (day 9: p = 0.0492; day 11: p = 0.0255; day 15: p = 5.66×10^-5^; non-sound responsive cohort: p > 0.89 for day 7, 9, 11 respectively and p = 0.0326 for day 15; control cohort: p > 0.9 for all post-ablation days; permutation test across groups, sound responsive cohort: p = 0.020, p = 0.11, p = 0.025, p = 0.005 for day 7, 9, 11, 15, respectively; non-sound responsive cohort: p > 0.39 for all post ablation days; control: p > 0.63 for all post-ablation days), we wondered if other changes in single-neuron properties than tuning width may mediate the later recovery of the representational map.

To test if changes in second-order response statistics in the local population responses could contribute to the recovery, we shuffled the data by permutating the neurons recorded on a given day across FOVs and mice and again constructed similarity matrices from the newly generated average population response vectors on a given day for every experimental cohort. With this manipulation, we selectively removed local correlations in tuning, without changing other single-neuron response properties such as tuning width and response amplitude. We observed that the normalized correlations in the similarity matrices obtained from the shuffled data showed in all cohorts of mice a general tendency to decline throughout the experiment (Fig. 3g). In the sound-responsive cohort the correlation values were lower as compared to the other cohorts on day 7, consistent with our previous analyses. However, the correlations remained low for all the other days after microablation and no recovery was observed in the shuffled data. (Fig. 3g; see Methods; the normalized correlation (*r_n_*) of neuron-shuffled populations was below the lower boundary (CI_L_) of the shaded area for all days after ablation in sound responsive cohort: day 7: *r_n_* = 0.619, CI_L_ = 0.712; day 9: *r_n_* = 0.594, CI_L_ = 0.664; day 11: *r_n_* = 0.694, CI_L_ = 0.700; day 15: *r_n_* = 0.627, CI_L_ = 0.661; permutation test across groups by shuffling neurons between sound responsive cohort and control cohort: p < 0.006 for all post-ablation days; permutation test between non-sound responsive cohort and control cohort: p = 0.010, p = 0.862, p = 0.186, p = 0.738 for day 7, 9, 11, 15, respectively; Extended Data Fig. 5i-k). These results indicate that changes in higher-order statistics in local response patterns may mediate the recovery of the representational map during later days after microablation.

As a simple readout of local higher-order response statistics, we quantified the fraction of pairs of responsive neurons with high signal correlations (> 0.7) on a given day. We observed a delayed increase in the sound responsive cohort for the neuron pairs with large best response amplitudes (Fig. 3h, left; two-way ANOVA, F(6, 270) = 3.43, p = 0.0028 across days, F(3, 270) = 8.33, p = 2.55 × 10^-5^ across amplitude bins). The fraction of neuron pairs with high signal correlations remained unaltered throughout the experiment in the two other cohorts (Fig. 3h, middle and right; two-way ANOVA, non-sound responsive ablation: F(6, 269) = 1.26, p = 0.274 across days, F(3, 269) = 0.894, p = 0.445 across amplitude bins; control: F(6, 241) = 1.910, p = 0.080 across days, F(3, 241) = 2.010, p = 0.113 across amplitude bins). In direct comparison to the other cohorts, the delayed increase was prominent only in the sound response cohort (Fig. 3i; two-sample t-test between baseline days vs. days after ablation with FDR correction for sound responsive cohort: p = 0.286, p = 0.0167, p = 0.0017, p = 1.05×10^-4^ for day 7, 9, 11, 15; non-sound responsive cohort: p = 0.0621, p = 0.0621, p = 0.423, p = 0.0621 for day 7, 9, 11, 15, respectively; control: p > 0.136 for all post-ablation days; permutation test across groups: sound responsive cohort: p = 0.645, p = 0.23, p = 0.035, p = 0.005 for day 7, 9, 11, 15, respectively; non-sound responsive cohort: p > 0.20 for all post-ablation days; control: p > 0.575 for all post-ablation days). The changes in pairwise signal correlations indicate that tuning of neurons in the spared network became locally more homogeneous.

In summary, our analyses of single-neuron response properties indicate that the transient reduction in the response reliability and the narrowing of the neurons’ tuning primarily cause the disturbance of the representational map and that the delayed increase in local signal correlations compensates these partially longer-lasting effects and contribute to the recovery of the representational map.

### The recovery of the representational map is supported by neurons becoming sound-responsive after microablation

The above analyses of the changes in single-neuron response properties considered the neurons measured on a given imaging day. They did not take into account the history of their sound-evoked responses in previous imaging sessions, information that is available from our longitudinally obtained data.

Our initial analyses of mice without microablation (Fig. 1) showed that in basal conditions individual cells undergo changes in their responsiveness to the set of sound stimuli over the course of several days. Thus, we next asked whether and how the drift in responsiveness is influenced by the targeted loss of neurons. Quantifying the overlap of neurons showing significant sound responses on two subsequent imaging days, we found a significant reduction of with stable responsiveness after microablation of sound responsive neurons that was maintained throughout the rest of the observation period. In the other cohorts, the likelihood of a neurons to change responsiveness remained at baseline level (Fig. 4a; one-way ANOVA, F(5, 54) = 6.03, p = 0.0002; two-sample t-test between change during baseline days vs. change from day 5 to day 7, p = 9.47×10^-5^; two-sample t-test with FDR correction on day 5 → 7 between the sound responsive cohort vs. control, p = 0.064; the sound responsive cohort vs the non-responsive cohort, p = 0.0165; one-way ANOVA in the non-responsive cohort, F(5, 54) = 0.39, p = 0.85; one-way ANOVA in control cohort, F(5, 48) = 0.50, p = 0.78). Thus, microablation of sounds responsive neurons induces a longer-lasting increase in the drift rate of auditory responses.

**Fig 4.**
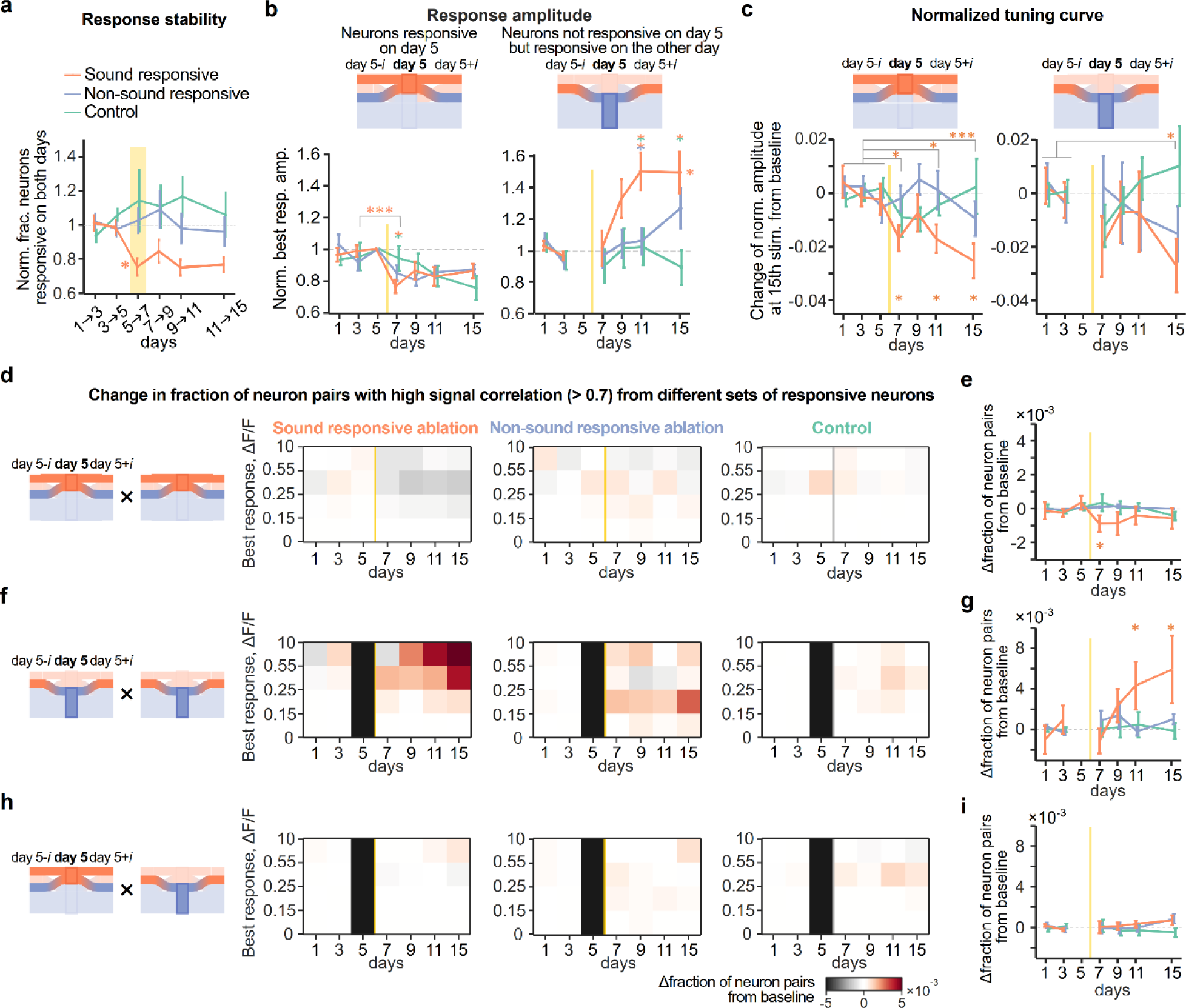
Major changes in single-neuron response properties are primarily mediated by neurons gaining responsiveness after microablation. **a.** Quantification of response stability as the fraction of neurons categorized as sound responsive on two consecutive imaging days. Color scheme for sound responsive cohort (orange), non-sound responsive ablation cohort (blue) and control cohort (gree) applies for this and the following panels. **b**. Top: Schematic illustrating the categorization of neurons according to their responsiveness on day 5 before microablation. One group considered all neurons responsive on day five and their responses on all other days, independent if they were also responsive or not. The other group considered neurons classified as responsive on a given day, that were however not responsive on day. Bottom: Average best response amplitude normalized to day 5 (left) or days 1 and 3 (right) in the three experimental cohorts. Note, that the strong increase in overall best response amplitudes as shown in Fig. 3d is mediated by neurons that were not responsive before microablation. **c**. Change of tuning width for neurons responsive on day 5 (left) and for neurons not responsive on day 5 but responsive on the other day (right) measured as normalized response amplitudes of the 15th largest stimulus in the sorted tuning curve (Same as Fig. 3f). **d**. Change in fraction of neuron pairs with high signal correlation that were categorized as responsive on day 5. Colormaps show difference in fraction normalized to baseline for four categories of neurons sorted their best response amplitude. Sound responsive cohort (left), non-sound responsive cohort (middle), control cohort (right). **e**. Change in fraction of neuron pairs with high signal correlations for neurons of the largest best response category in **d**. **f**-**g**. Same as **d**-**e**, but for pairs of neurons categorized as not responsive on day 5 but responsive on the other day. **h**-**i**. Same as **d**-**e**, but for pairs of neurons in which one neuron was categorized responsive on day 5 and the other categorized not responsive on day 5 but responsive on the other day. Note strong increase in neuron pairs with high signal correlations that were categorized as not responsive before microablation.

We next tested how microablation-induced changes in response properties relate to the history of responsiveness of individual neurons. We considered two categories of neurons: Neurons displaying significant sound responses on day 5 before the microablation and neurons with no significant sound responses on day 5, but that were responsive on another imaging day.

We revisited the effect of increased best response amplitudes after microablation of sound responsive neurons. For neurons of the first cateogory that were responsive before microablation, best response amplitudes on day 7 after microablation were significantly smaller in mice with sound responsive neurons ablated, and less prominently reduced in the non-sound responsive cohort (Fig 4b left; two-sample t-test between day 3 vs. day 7, for sound responsive cohort: p = 4.54 × 10^-4^; for non-sound responsive cohort, p = 0.33; for control, p = 0.63; group comparison by one-sample t-test with FRA correction between sound responsive cohort vs control on day 7, p = 0.039; sound responsive cohort vs non-sound responsive cohort, p = 0.096). When considering only those neurons that stayed responsive also after microablation, the best response amplitude maintained comparable to the baseline (Extended Data Fig. 6a), suggesting that the microablation-induced reduction in the average best amplitude is primarily caused by neurons losing responsiveness after microablation.

In the second category of neurons, those that did not show sound responses on day 5 before microablation but on other days before or after microablation^42^ (Extended Data Fig. 6b), best response amplitudes displayed a delayed but substantial increase after microablation of sound-responsive neurons (Fig. 4b right; one-way ANOVA across days for sound responsive cohort: F(5, 54) = 6.19, p = 0.0001; non-sound responsive cohort: F(5, 54) = 2.14, p = 0.075; control: F(5, 48) = 0.45, p = 0.81; two-sample t-test with FDR correction between sound responsive cohort vs. control: p = 0.74, p = 0.14, p = 0.017, p = 0.0096 for day 7, 9, 11, 15, respectively; sound responsive cohort vs. non-sound responsive cohort: p = 0.74, p = 0.14, p = 0.017, p = 0.23 for day 7, 9, 11, 15; non-sound responsive cohort vs. control: p = 0.74, p = 0.88, p = 0.82, p = 0.068 for day 7, 9, 11, 15). This indicates that the increase of best response amplitudes in responsive neurons on a given day shown in Fig. 3d is primarily caused by neurons that have gained responsiveness after microablation.

We wondered how far the microablation-induced effects on best response amplitude are propagated spatially in the cortical network (Extended Data Fig. 6c-f). We found that in the sound responsive ablation cohort, in which a change in the best response was observed, the change was primarily observed in neurons in the proximity (<∼200 μm) of ablated neurons.

When re-analyzing the narrowing of tuning width shown in Fig. 3f for neurons categorized as responsive or non-responsive on the day before microablation, we observed in the sound responsive ablation cohort a more pronounced narrowing in the neurons responsive before micoablation (Fig. 4c left; two-sample t-test between baseline days vs. days after ablation. day 7: p = 0.0113; day 9: p = 0.250; day 11: p = 0.0113; day 15: p = 0.0010). In both non-sound responsive ablation and control cohort, on the other hand, the tuning width did not change over days (Fig. 4c left; non-sound responsive cohort: p > 0.48 for all post-ablation days; control: p > 0.08 for all post-ablation days; permutation test of sound responsive ablation across groups: p = 0.040, p = 0.245, p < 0.005, p = 0.0150 for day 7, 9, 11, 15, respectively). Neurons unresponsive on day 5 but being responsive on any other day in all three experimental cohorts exhibited a less reduction in their tuning width (Fig. 4c right; two-sample t-test between baseline days vs. days after ablation. sound responsive cohort: p = 0.139, p = 0.573, p = 0.573, p = 0.0343 for day 7, 9, 11, 15, respectively; non-sound responsive cohort: p > 0.52 for all post-ablation days; control: p > 0.51 for all post-ablation days; permutation test of sound responsive ablation across groups: p = 0.170, p = 0.39, p = 0.405, p = 0.065 for day 7, 9, 11, 15, respectively; Extended Data Fig. 7a). This indicates that the major contributor for narrowing in tuning on a given day are neurons responsive before microablation.

We also revisited the increase in neuron pairs with high signal correlations in the sound responsive ablation cohort (Fig. 3h,i) by categorizing neurons, again, according to their responsiveness on the day before microablation. We observed a slight decrease in neuron pairs between neuron responsive before microablation for sound responsive cohort, but no change for the other cohorts (Fig. 4d left; sound responsive cohort, two-way ANOVA, F(6, 270) = 4.87, p = 9.52×10^-5^ across days; F(3, 270) = 5.57, p = 0.0010 across amplitude bins; two-sample t-test between baseline days and days after ablation (day 9-15), p = 8.68×10^-6^; Fig. 4d middle; non-sound responsive cohort, two-way ANOVA, F(6, 270) = 0.426, p = 0.861 across days; F(3,270) = 1.035, p = 0.378 across amplitude bins, t-test, p=0.098; Fig. 4d right; control cohort, two-way ANOVA, F(6, 242) = 2.25, p = 0.0390 across days; F(3, 242) = 0.025, p = 0.995 across amplitude bins; t-test, p = 0.405; Fig. 4e; permutation test across groups to sound responsive cohort: p = 0.010, p = 0.0350, p = 0.130, p = 0.205 for day 7, 9, 11, 15, respectively; non-sound responsive cohort: p > 0.71 for all post-ablation days; control: p > 0.41 for all post-ablation days). However, neuron pairs both unresponsive on day 5 but being responsive on any other day showed a stark increase in the fraction of highly correlated neuron pairs 3-9 days after sound responsive ablation, but no change after non-sound responsive ablation and control (Fig. 4f left; sound responsive cohort two-way ANOVA, F(5, 231) = 3.02, p = 0.0117 across days; F(3, 231) = 3.09, p = 0.0279 across amplitude bins; two-sample t-test between baseline days and days after ablation (day 9-15), p = 0.0337; Fig. 4f middle; non-sound responsive cohort F(5, 229) = 1.427, p = 0.215 across days; F(3,229) = 2.749, p = 0.0436 across amplitude bins; t-test, p = 0.222; Fig. 4f right; control cohort F(5, 206) = 0.583, p=0.713 across days; F(3, 206) = 0.508, p = 0.677 across amplitude bins; t-test, p = 0.792; Fig. 4g; permutation test across groups to sound responsive cohort: p = 0.905, p = 0.205, p = 0.040, p = 0.010 for day 7 , 9, 11, 15, respectively; non-sound responsive cohort: p > 0.11 for all post-ablation days; control cohort: p > 0.405 for all post-ablation days). Signal correlations in pairs of neurons in which one was categorized as responsive and the other unresponsive before microablation, did not change after ablation (Fig. 4h left; sound responsive cohort, two-way ANOVA: F(5, 217) = 0.334, p = 0.892 across days; F(3, 217) = 2.85, p = 0.038 across amplitude bins; Fig. 4h middle: non-sound responsive cohort: F(5, 221) = 0.25, p = 0.939 across days; F(3, 221) = 0.070, p = 0.976 across amplitude bins; Fig. 4h right; control cohort F(5, 195) = 0.424, p = 0.832 across days; F(3, 195) = 2.53, p = 0.059 across amplitude bins; Fig. 4i; permutation test across groups for all post-ablation days: p > 0.175, p > 0.11, p > 0.43 in sound responsive cohort, non-sound responsive cohort, control, respectively). However, in a subset of these pairs, namely between neurons maintaining responsiveness on both day 5 and another day and neurons unresponsive before microablation, a strong increase in the fraction of high signal correlations was observed after sound responsive ablation, but not after non-sound responsive ablation nor control (Extended Data Fig. 7b,c).

Together, our analysis of the increase in neuronal pairs with high signal correlations that takes the response history of neurons into account, revealed that again neurons gaining responsiveness after microablation have a disproportionally large impact.

When characterizing how these shifts in tuning related to those of the microablated neurons that were removed from the network, we observed that in the sound responsive cohort the tuning of spared neurons responsive before microablation became comparably less similar to that of the microablated neurons. In contrast, neurons gaining responsiveness in later days after microablation were more similar to those of the microablated neurons (Extended Data Fig. 7d,e).

### Excitatory and inhibitory neurons are differentially affected after microablation

Adjustments in the balance of activities in the populations of excitatory and inhibitory neurons have been described in a number of contexts where a reorganization of neuronal networks is induced^36,46^. To specifically assess activity in inhibitory and non-inhibitory neurons in our experiments, we co-injected in a subset of mice of each experimental cohort (n = 5 in sound responsive ablation; n = 5 in non-sound responsive ablation; n = 7 in control) an AAV vector labeling inhibitory cortical neurons (AAV mDlx-NLS-tagBFP) together with the AAV vectors mediating expression of GCaMP6m and H2B::mCherry (Fig. 5a). This led to a co-labeling of tagBFP in ∼10% of the nuclei showing mCherry expression. In control experiments using a GAD67-GFP pan-interneuron reporter mouse line, we observed highly specific expression of tagBFP in GFP positive neurons (Extended Data Fig. 8a,b).

**Fig 5.**
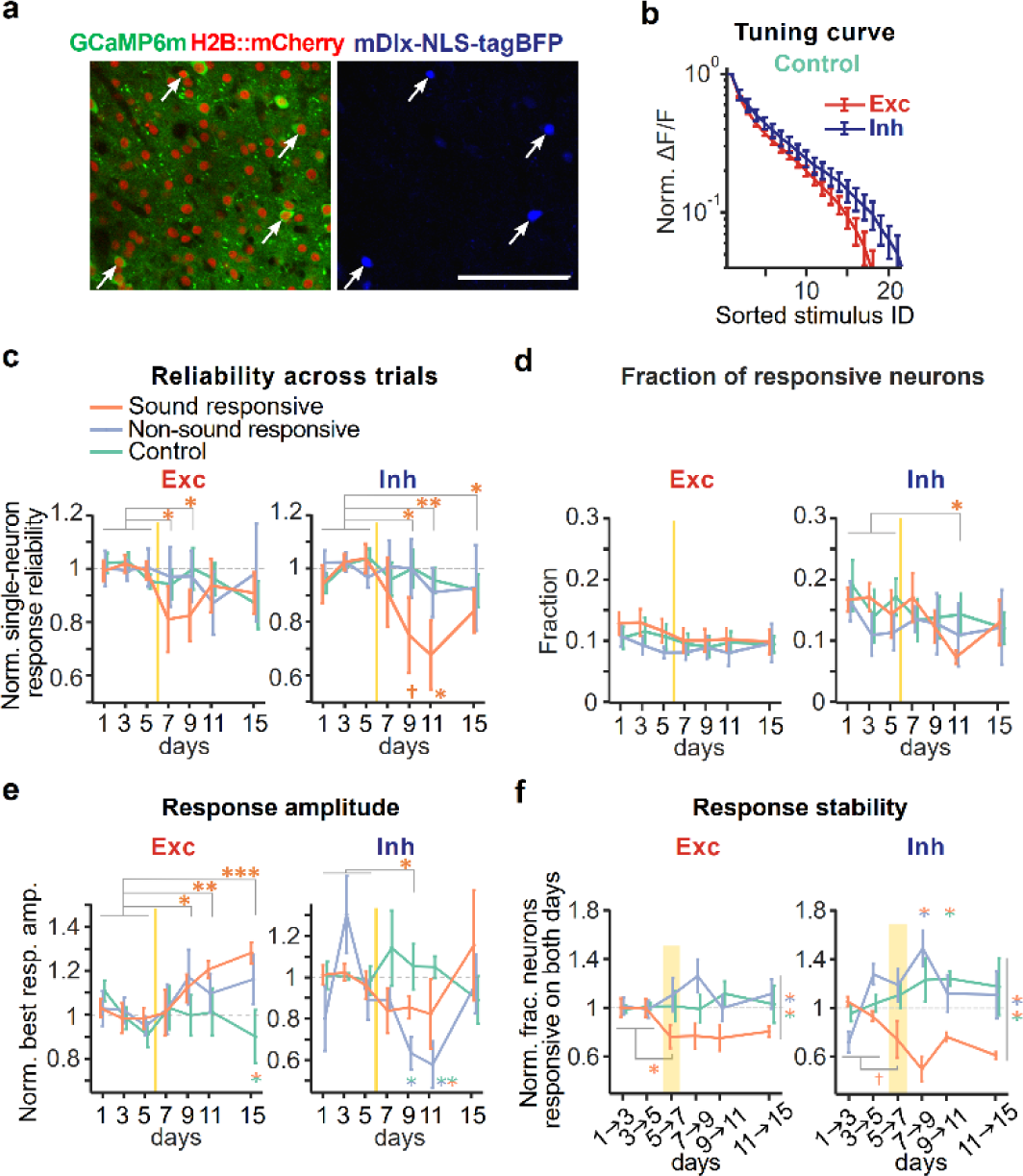
Microablation differentially affects population responses of excitatory and inhibitory neurons. **a**. A field of view in mouse auditory cortex showing broad expression of GCaMP6m and H2B::mCherry (left) and selective expression of nuclear localized tagBFP in interneurons (right). Scale: 100 μm. **b**. Averaged tuning curves of neurons responsive to sound stimuli normalized to the maximal response for excitatory (Exc: dark red line) and inhibitory neurons (Inh: dark blue line) during a baseline day in the control group. **c**. Same as Fig. 3a, but normalized response reliability across trials of single neurons averaged for all excitatory (left) and all inhibitory (right) neurons on each given day, in the three experimental cohorts. Data presented as mean ± s.e.m. across mice (5 mice for sound responsive ablation; 5 mice for non-sound responsive ablation; 7 mice for control). **d**. Fraction of sound responsive neurons for excitatory (Left) and inhibitory (Right) neurons for the three experimental cohorts. **e**. Same as Fig. 3d, but normalized best response amplitude of excitatory (left) and inhibitory (right) neurons responsive for each day. **f**. Same as Fig. 4a, but normalized fraction of neurons responsive on both days for excitatory (left) and inhibitory (right) neurons, in the three experimental cohorts.

Consistent with previous studies^47–49^, we observed during baseline conditions a moderately broader tuning in inhibitory neurons than in excitatory neurons (Fig. 5b and Extended Data Fig. 8c). We next asked if the microablation-induced effects observed in our previous analyses would affect excitatory and inhibitory neurons in the spared network to a similar or different degree. Similar to Fig. 3a, the response reliability across trials in excitatory neurons was lower after microablation of sound responsive neuron from day 7 and recovered during later days (Fig. 5c left; two-sample t-test of normalized reliability between baseline days vs. days after ablation with FDR correction. sound responsive cohort: p = 0.0413, p = 0.0413, p = 0.329, p = 0.139 for day 7, 9, 11, 15, respectively; non-sound responsive cohort: p > 0.60 for all post-ablation days; control: p > 0.14 for all post-ablation days; one-sided p-values in permutation test across groups, for sound responsive cohort: p = 0.0475, p = 0.0350, p = 0.263, p = 0.265 for day 7, 9, 11, 15, respectively; non-sound responsive cohort: p > 0.13 for all post-ablation days; control: p > 0.16 for all post-ablation days). Response reliability across trials in inhibitory neurons was also reduced in the sound responsive ablation cohort, but this effect appeared delayed and more pronounced (Fig. 5c right; two-sample t-test of normalized reliability between baseline days vs. days after ablation with FDR correction. sound responsive cohort: p = 0.323, p = 0.0325, p = 0.0074, p = 0.0486 for day 7, 9, 11, 15, respectively; non-sound responsive cohort: p > 0.66 for all post-ablation days; control: p > 0.36 for all post-ablation days; one-sided p-values from permutation test across groups, for sound responsive cohort: p = 0.145, p = 0.0125, p = 0.0025, p = 0.118 for day 7, 9, 11, 15, respectively; non-sound responsive cohort: p > 0.32 for all post-ablation days; control cohort: p > 0.27 for all post-ablation days). Microablation induced a transient disruption of the response reliability in both excitatory and inhibitory neurons, that appeared more pronounced in inhibitory neurons.

The fraction of neurons showing a response to the set of sounds on a given day was largely comparable between excitatory and inhibitory neurons, with the fraction of sound-responsive inhibitory neurons being slightly higher than that of sound-responsive excitatory neurons during baseline days (Fig. 5d). The fraction of neurons contributing to the population response on a given day was largely unaffected by the microablation procedure in both inhibitory and excitatory neurons, except for a reduction on day 11 in inhibitory neurons (Fig. 5d; two-sample t-test of fraction between baseline days vs. days after ablation with FDR correction; excitatory neurons in sound responsive cohort for all post-ablation days: p > 0.30; non-sound responsive cohort for all post-ablation days: p > 0.90; control cohort: p > 0.61; inhibitory neurons in sound responsive cohort: p = 0.022 on day 11, but p > 0.47 for the other post-ablation days; non-sound responsive cohort: p > 0.97; control: p > 0.57 for all post-ablation days, respectively).

Next, we investigated how best response amplitudes were affected in both cell types after microablation. While the best response amplitudes of excitatory neurons showed an increase after ablating sound responsive neurons, the best response amplitudes of inhibitory neurons decreased until day 9 but renormalized by day 15 (Fig. 5e left; one-sided t-test of best response between baseline days vs. days after ablation in excitatory neurons with FDR correction. sound responsive cohort: p = 0.446, p = 0.0449, p = 0.0017, p = 0.0003 for day 7, 9, 11, 15, respectively; permutation test of sound responsive ablation cohort to control cohort: p > 0.05, for day 7, 9, 11, p < 0.025 for day 15, respectively; Fig. 5e right; one-sided t-test between baseline days vs. days after ablation in inhibitory neurons with FDR correction. sound responsive cohort: p = 0.0529, p = 0.0434, p = 0.0778, p = 0.174 for day 7, 9, 11, 15, respectively; permutation test of sound responsive ablation cohort to control cohort: p > 0.05, for day 7, 9, 15, p < 0.05 for day 11, respectively; baseline normalized amplitude on day 15: 1.15 ± 0.26; Extended Data Fig. 8d). In the non-sound responsive cohort, the best response amplitudes did not clearly change in excitatory neurons, but significantly decreased with subsequent renormalization on day 15 in inhibitory neurons (Fig. 5e left; one-sided t-test of excitatory neurons in non-sound responsive cohort: p = 0.38, p = 0.076, p = 0.148, p = 0.076 for day 7, 9, 11, 15, respectively; permutation test of non-sound responsive ablation cohort to control cohort: p > 0.05, for all post-ablation days; Fig. 5e right; one-sided t-test of inhibitory neurons with FDR correction in non-sound responsive cohort: p = 0.37, p = 0.0591, p = 0.0591, p = 0.439 for day 7, 9, 11, 15, respectively; permutation test of non-sound responsive ablation cohort to control cohort: p > 0.05, for day 7, 15, p < 0.025 for day 9, 11, respectively) . In contrast, the best response amplitudes in the control group remained largely unaltered (Fig. 5e left; excitatory neurons: p > 0.48 for all post-ablation days; inhibitory neurons: p > 0.25 for all post-ablation days). In order to obtain an estimation of the total sound-evoked activity in excitatory and inhibitory neurons, we calculated the product of the fraction of responsive neurons and the average best response amplitude for both neuronal populations (see Methods). Assessing the balance of general excitation and inhibition levels in the local network, we observed a shift in the total sound-evoked activity towards the population of excitatory neurons in both sound responsive and non-sound responsive cohorts 3 to 5 days after microablation that returned to a level comparable to baseline on day 15 (Extended Data Fig. 8e). We did not observe altered decay time constants of the calcium transients throughout the experiment, corroborating the idea that the changes in best response amplitudes reflect microablation-induced differences in neuronal activity, rather than alterations in the expression levels of calcium binding proteins (Extended Data Fig. 8f,g).

When revisiting the microablation-induced changes in tuning width (Extended Data Fig. 8h), signal correlations (Extended Data Fig. 8i) and best response amplitudes in neurons categorized responsive or unresponsive before microablation (Extended Data Fig. 8j,k), we did not detect significant differences in excitatory or inhibitory neurons, likely also due to the smaller number of inhibitory neurons in the sample lowering the sensitivity of our analyses.

We next examined the rate at which individual excitatory or inhibitory neurons show changes in their responsiveness and analogous to Fig. 4a quantified the overlap of neurons showing significant sound responses on two consecutive imaging days. We found that the fraction of neurons showing stable tuning was reduced in both excitatory and inhibitory neurons during the days following the microablation of sound responsive neurons with a more pronounced reduction in inhibitory neurons (Fig. 5f; two-sided t-test between the overlap during baseline days (1→3, 3→5) vs. the overlap during post-ablation days (5→7, 7→9, 9→11, 11→15) for excitatory neurons: sound responsive ablation, p = 0.0024; non-sound responsive ablation, p = 0.21; control, p = 0.67; for inhibitory neurons: sound responsive ablation, p = 7.49 × 10^-4^; non-sound responsive ablation, p = 0.15; control, p = 0.15; two-sided t-test with FDR adjustment for group comparison in the overlap during post-ablation days, in excitatory neurons: sound responsive ablation vs. non-sound responsive ablation, p = 9.11 ×10^-5^; sound responsive ablation vs. control, p = 0.003; non-sound responsive ablation vs. control, p = 0.34; in inhibitory neurons: sound responsive ablation vs. non-sound responsive ablation, p = 1.79 × 10^-4^; sound responsive ablation vs. control, p = 3.99 ×10^-4^; non-sound responsive ablation vs. control, p = 0.76).

Together, our analysis showed for several response measures including response reliability, fraction of responsiveness and tuning width, similar microablation-induced effects in excitatory and inhibitory populations, suggesting a general degree of co-tuning. The reorganization of the local network after microablation was accompanied with a shift in the levels of sound-evoked responses towards excitatory neurons.

### A longer compensatory period after the selective ablation of inhibitory neurons

Interneurons have been suggested to play a particular important role in controlling the stability or plasticity of activity patterns in cortical networks^24,46,50^. We therefore wondered how the stability of the representational map would be affected by the selective removal of inhibitory neurons from the network. As the vast majority of the microablated neurons in both the sound responsive cohort and the non-sound responsive cohort were putatively excitatory neurons (Extended Data Fig. 4a and 8), we conducted another experimental series in which we used the mDlx-driven blue fluorescent reporter to selectively target inhibitory neurons for microablation (n = 6 mice, Fig. 6a). As before, we obtained a distribution of sound-evoked response amplitudes from the neurons of each FOV during baseline (day 3 and 5) and identified the inhibitory neurons (inset in Fig. 6b). We primarily targeted sound responsive inhibitory neurons, however, given the much lower density compared to excitatory neurons (∼10% of the local population, Extended Data Fig. 8a) we additionally microablated inhibitory neurons categorized as non-responsive, in order to remove in total a comparable number as in the other experimental cohorts (Extended Data Fig. 4a-c, see Methods).

**Fig 6.**
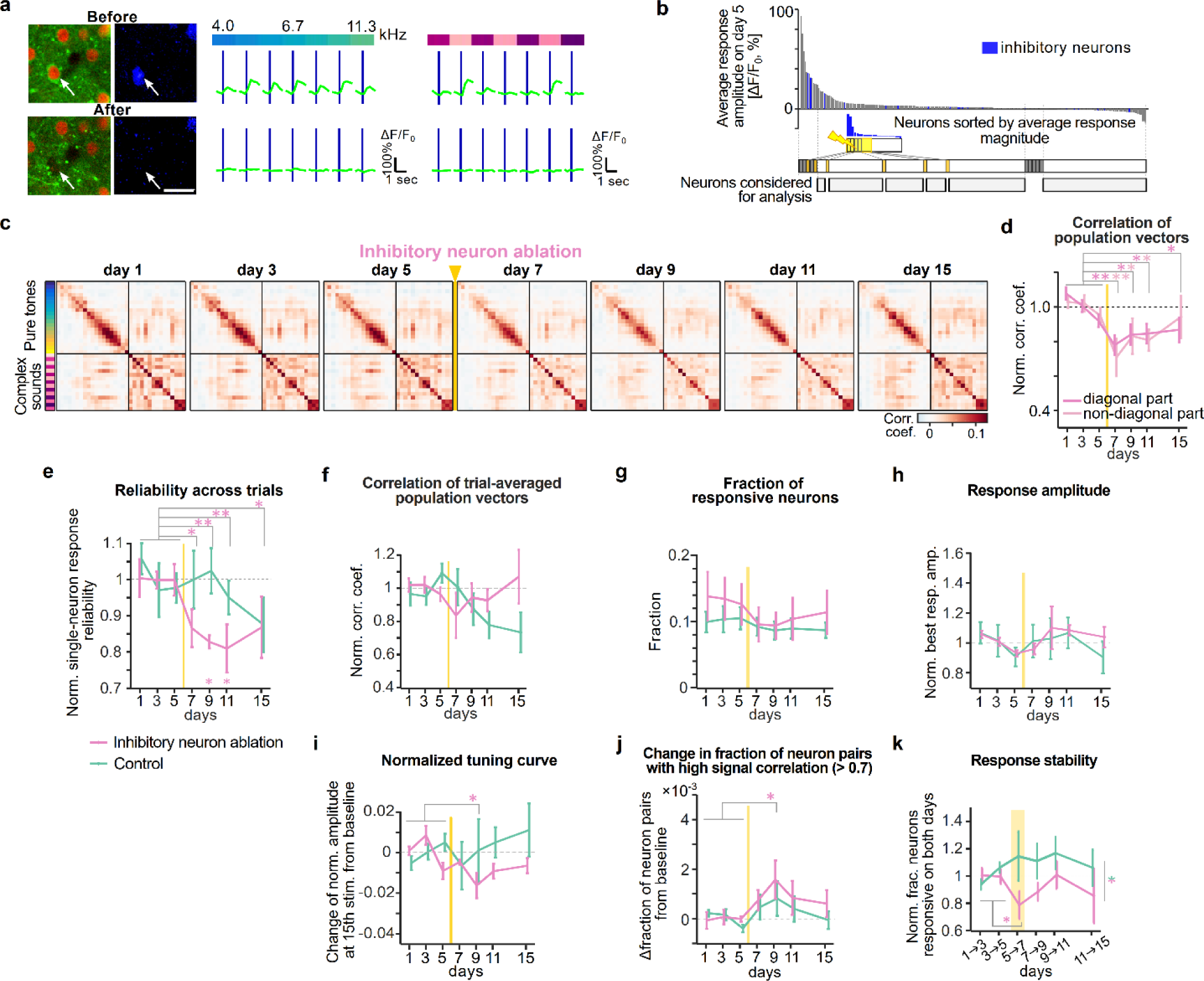
Targeted microablation of inhibitory neurons causes a longer-lasting disturbance of the representational map. **a**. Representative example of microablation of a targeted inhibitory neuron with PT & CS tuning. Scale: 20 μm. **b**. Target neurons for inhibitory neuron microablation. Top: Sorted sound-evoked amplitudes before microablation in a FOV. Interneurons are labeled as blue bars and excitatory neurons as gray bars. Middle: Sorted amplitudes of the same population above but only from interneurons. 4 - 8 neurons of highly responsive interneurons were targeted per FOV. Bottom: Experimentally targeted neurons and neurons corresponding to target groups in responsive and non-responsive cohorts were excluded from analysis. Analysis was conducted on remaining spared neurons (gray bars) **c**. Similarity matrix of population vectors for all stimuli averaged across all FOVs for each day in inhibitory neuron ablation cohort. **d**. Baseline-normalized correlations averaged across diagonal elements (dark pink) and off-diagonal elements (light pink) in the similarity matrices in **c**. **e**. Baseline-normalized response reliability across trials of single neurons averaged across all neurons on each given day (mean ± s.e.m. across mice). Pink line indicates data from inhibitory ablation cohort, green line indicates data from control group, as in the following panels. **f**. Baseline-normalized correlations averaged across off-diagonal elements in the similarity matrix constructed from trial-averaged population response vectors. **g.** Fraction of sound-responsive neurons over the imaging days. **h**. Baseline-normalized best response amplitude of neurons responsive for each day. **i**. Change of normalized response amplitudes from baseline at 15^th^ largest stimulus index in the tuning curve (Extended Data Fig. 9d) over days. **j**. Change in fraction of neuron pairs with high signal correlation from baseline at the largest best response category (Extended Data Fig. 9f). **k**. Quantification of response stability as the fraction of neurons categorized as sound responsive on two consecutive imaging days. Please compare panels **c**,**d** to Fig. 2, panels **e-j** to Fig. 3 and panel **k** to Fig 4.

We found that the removal of inhibitory neurons led to a disturbance in the structure of the representational map on the day after microablation, similar to that previously seen in the sound responsive ablation cohort. (Fig. 6c; two-sample t-test of normalized average correlation between baseline days vs. day 7, in diagonal elements: p = 4.84 × 10^-4^; in off-diagonal elements: p = 7.87 × 10^-4^). However, whereas in the sound responsive ablation cohort the structure of the map showed no longer statistical differences to the baseline 3 days after microablation, in the inhibitory neuron ablation cohort, this process appeared to be substantially delayed (Fig. 6c and d; two-sample t-test of normalized average correlation between baseline days vs. days after ablation with FDR correction, in diagonal elements: p = 0.0110, p = 0.0110, p = 0.0426 for day 9, 11, 15, respectively; off-diagonal elements: p = 0.0388, p = 0.0047, p = 0.500 for day 9, 11, 15, respectively; one-way ANOVA of normalized average correlation across days in diagonal elements: F(6, 34) = 4.11, p = 0.0033; off-diagonal elements: F(6, 34) = 2.19, p = 0.068; Extended Data Fig. 9a-c).

Next, we analyzed the single-neuron response properties associated with the removal of inhibitory neurons. Starting with the response reliability across trials, we observed a substantial reduction on day 7, that, in contrast to the sound responsive cohort, remained at low levels throughout the observation period (Fig. 6e; one-way ANOVA, F(6, 34) = 2.76, p = 0.0271, two-sample t-test between baseline days vs. day 7, 9, 11, and 15, p = 0.0158, 0.0014, 0.0036, 0.0482 after FDR correction; group comparison between inhibitory neuron cohort and control: p = 0.238, p = 0.0283, p = 0.0979, p = 0.964 for day 7, 9, 11, 15, respectively). To assess if additional changes in the tuning properties contribute to the disturbance of the representational map, we again calculated the average off-diagonal correlation in the similarity matrix constructed from trial-averaged population responses. This, however, did not exhibit microablation-related effects during the days after the manipulation (Fig. 6f; one-way ANOVA, F(6, 34) = 0.68, p = 0.66; two-sample t-test of normalized correlation coefficient between baseline days vs. days after ablation with FDR correction. p > 0.26 for all post-ablation days; permutation test across inhibitory neuron ablation and control cohorts: p = 0.21, p = 0.685, p = 0.865, p = 0.925 for day 7, 9, 11, 15, respectively). Therefore, microablation of inhibitory neurons appears to primarily affect the reliability of sound responses across trials in the local network and does not result in major effects on single-neuron tuning properties.

Consistently, repeating our analyses on the dataset from the inhibitory neuron cohort did not reveal changes in single-neuron tuning properties that were comparable to the sound responsive cohort (Fraction of responsive neurons: Fig. 6g, one-way ANOVA, F(6, 34) = 0.36, p = 0.90); Best response amplitudes: Fig. 6h, one-way ANOVA across days, F(6, 34) = 0.95, p = 0.47; two-sample t-test between baseline days vs. days after ablation. p = 0.427, p = 0.427, p = 0.131, p = 0.427 for day 7, 9, 11, 15, respectively; permutation test across groups: p = 0.68, p = 0.375, p = 0.355, p = 0.125 for day 7, 9, 11, 15, respectively; Width of tuning: Fig. 6i, two-sample t-test between baseline days vs. days after ablation with FDR correction: p = 0.337, p = 0.0428, p = 0.171, p = 0.337 for day 7, 9, 11, 15, respectively; permutation test for comparison between inhibitory neuron ablation and control: p = 0.485, p = 0.175, p = 0.0850, p = 0.155 for day 7, 9, 11, 15, respectively; see also Extended Data Fig. 9d; Fraction of pairs of neurons with high signal correlations: Fig. 6 j, two-sample t-test between baseline days vs. days after ablation with FDR correction: p = 0.079, p = 0.0174, p = 0.103, p = 0.147 for day 7, 9, 11, 15, respectively; permutation test across inhibitory neuron ablation and control: p = 0.365, p = 0.25, p = 0.265, p = 0.17 for day 7, 9, 11, 15, respectively; Extended Data Fig. 9e,f).

Quantifying the overlap of neurons being categorized as responsive on two consecutive imaging days, we found a reduction after microablation of inhibitory neurons, indicating increased changes in responsiveness of individual neurons across days (Fig. 6k; two-sided t-test of normalized fraction of stably responsive neurons during baseline days (1→3, 3→5) vs. day 5 → 7, p = 0.026; two-sample t-test between inhibitory neuron ablation cohort vs. control cohort during post-ablation days (7→9, 9→11), p = 0.0475). In addition, similarly in the sound responsive cohort, we found that neurons acquiring responsiveness after inhibitory neuron ablation, had a higher likelihood of sharing the tuning profile of those neurons that were removed from the network (Extended Data Fig. 9g, h).

Together, we observed that the targeted ablation of different neuron classes from the network caused specific changes in single-neuron properties at multiple timescales that underlay a transient impairment of the representational map, but also its subsequent recovery.

## Discussion

We used the representational map, formed by the population responses to a set of auditory stimuli, as a readout of auditory cortical function^5^. We observed that under constant environmental and behavioral conditions, individual neurons show a high degree of volatility in their functional tuning properties^12^, similar to previous reports in auditory cortex and other brain regions^9,12–15,19,51,52^. Interestingly, the relative distances of the individual stimuli in the neuronal coding space remain unaffected by the ongoing remodeling of tuning properties and thus the structure of the representational map is stably maintained^16^. Given that the overall responsiveness of neurons is largely stable, such changes are consistent with rotational changes in the geometry of the representation^45^.

To study the effect of the loss of neurons on network function, we selectively removed 30-40 sound responsive neurons. We found that this manipulation was sufficient to transiently disrupt the correlational structure of the population responses underlying the representational map. Linking this global change in sensory representation to properties of individual neurons^45^, we delineated the contributors to the disruption as well as the homeostatic recovery of the map: The removal of excitatory sound responsive neurons led to a transient decrease in the reliability of sound-evoked responses and a longer-lasting narrowing of tuning in the spared neurons, both associated with a disturbance of the representational map. However, within three days a recovery was mediated by neurons gaining responsiveness and showing a high degree of pairwise correlated tuning curves (Extended Data Fig. 10a). Interestingly, removal of inhibitory neurons led to a disturbance of the representational map taking more than five days to recover and that was primarily associated with a temporary decrease in the reliability of sound-evoked responses (Extended Data Fig. 10b)^53,54^.

The high sensitivity in the similarity matrix to the selective elimination of a small number of neurons suggests that the apparent stability of the representational map observed during basal conditions does not reflect a highly redundant or intrinsically stable network configuration, but relies on the ability of plasticity mechanisms to constantly keep the network in a functional configuration^55–59^. While such conservative mechanisms seem to be sufficiently effective during basal conditions when ongoing tuning changes occur, they can be at least temporarily challenged when specific neurons are removed from the network within a short time. Selective changes in circuit function and even behavior by subtle manipulations of neuronal activity have been demonstrated previously in various contexts^27,29,30,44,60,61^.

As mentioned above, theoretically three major scenarios were conceivable regarding the long-term dynamics of the map after its disruption: i) Given the permanent nature of cell ablation, the impairment of the map could have been long-lasting. ii) The ongoing volatility of tuning observed in individual neurons during baseline could have simply continued and have led to a gradual washout of the microablation-induced effects. iii) Specific homeostatic processes are engaged to support a recovery of the representational map. Our data showing a delayed increase in signal correlations in pairs of neurons (Fig. 3h) and a delayed imbalance of response amplitudes in excitatory and inhibitory neurons (Fig. 5e) only few days after the microablation are consistent with the third scenario. Interestingly, similar increases in pairwise correlations^62^ and shifts in the responses and the synaptic structures of excitatory and inhibitory neurons^50,63–66^ have been also observed in different conditions when cortical circuits needed to adapt to overall activity silencing, prolonged or permanent perturbations of sensory inputs.

In how far homeostatic mechanisms play a role in protecting higher-order brain functions in the cortex is less well understood, as classic schemes for homeostatic controllers^67,68^ with firing rate as a controlled variable are insufficient to explain homeostatic behaviors at different network scales^69^. Compensation of permanent manipulation and the setback of the controlled variable to its set point, represents hallmarks of a homeostatic control circuit. In the context of our observations, the compensation of the permanent removal of neurons from a local microcircuit leading to a dynamic reorganization of the network, indicates that the correlational structure of population activity underlying a representational map is not invariably stable, but could represent a controlled variable underlying homeostatic regulation^38^.

We speculate that the ability to compensate for the loss of microablated neurons by the selective recruitment of previously unresponsive neurons at the microcircuit level may also serve as a mechanism by which brain function may be upheld for extended periods of neuronal loss during aging and prodromal stages of neurodegenerative disease^1,2^.

## Acknowledgement

This work was supported by research grants Deutsche Forschungsgemeinschaft CRC1080-C05, Deutsche Forschungsgemeinschaft SPP 2041 Project #347573108, Deutsche Forschungsgemeinschaft/Agence nationale de la recherche Project #431393205 and Deutsche Forschungsgemeinschaft DIP “Neurobiology of Forgetting”.

## Author information

Yonatan Loewenstein is the incumbent of the David and Inez Myers Chair in Neural Computation.

## Author contributions

T.N. contributed to study conception, performed data collection, data analysis and contributed to manuscript writing; E.K. provided research materials; J.-B.E. provided data analysis tools and contributed to manuscript editing; D.F.A. provided data analysis tools; M.K. contributed to study conception and manuscript editing; Y.L. contributed to study conception and manuscript writing; S.R. conceived the study, provided funding and contributed to manuscript writing.

## Corresponding authors

Correspondence to Simon Rumpel.

## Data and code availability

Data will be made available upon reasonable request by the corresponding author. In addition, we are in the process in uploading data (G-node) and code (Zenodo) to publicly available repositories, as in our previous studies (Aschauer et al., 2022).

## Methods

### rAAV cloning and production

For rAAV genome encoding for GCaMP6m under the human SynapsinI promoter, and the rAAV encoding for H2B-mCherry, two plasmids were generated as described before^12^. To elaborate the plasmid containing a gene coding the blue fluorescent protein (tagBFP), preceded with three nuclear localization signals under the mDlx enhancer sequence (pAAV-mDlx-NLS-tagBFP), we started from the commercially available plasmid, AAV-mDlx-NLS-mRuby2 (Addgene plasmid #99130) and a plasmid for tagBFP in our lab stock (TRE-NLS-tagBFP, which is modified from Addgene plasmid #92202). The NLS-tagBFP sequence was PCR amplified from plasmid pAAV-TRE-NLS-tagBFP. The resulting PCR product was digested with the respective enzymes and purified. From AAV-mDlx-NLS-mRuby2, NLS-mRuby2 was excised with the restriction enzyme, and the NLS-BFP sequence was ligated into the cut and purified. The integrity of the final plasmid pAAV-mDlx-NLS-tagBFP was confirmed by sanger sequencing. All plasmids described above were packaged into AAV8 capsid described as previously described^70^.

### Mice

Experimental subjects were male C57BL/6JRj mice of 4-6 weeks of age obtained from Janvier laboratories. Before surgical procedures, mice were kept in groups of four, and housed in 530 cm^2^ cages on a 12h light/dark cycle with unlimited access to dry food and water. Experiments were carried out during the light period. All animal experiments were performed in accordance with the German laboratory animal law guidelines for animal research and had been approved by the Landesuntersuchungsamt Rheinland Pfalz (Approval 23 177-07/G 17-1-051).

### Stereotaxic injections and cranial window implantation

We followed the procedures described in detail previously^12^. Briefly, injections were performed perpendicular to the surface of the skull. Virus solution consisted of a mixture of 2 different rAAVs (rAAV8-hSyn-GCaMP6m-WPRE-hGHpolyA; titer: 1.35 × 10^11^ viral genomes (VG)/ml; rAAV8-hSyn-H2BmCherry-hGHpolyA; titer: 2 × 10^13^ VG/ml) in PBS. For another subset of experiments, rAAV8-mDlx-NLS-tagBFP-WPRE-hGHpolyA (titer: 2.16 × 10^11^ VG/ml) was additionally mixed with the foregoing rAAVs. 175 nL of the virus mixture loaded into a glass pipette was injected in 5 locations (20 nL/min, a total volume of 875 nL) along the anterior–posterior axis with coordinates: 4.4, −2.3/−2.6/−2.9/−3.2/−3.5, 2.5 (in mm, caudal, lateral, and ventral to Bregma) to cover the right auditory cortex. Several days after the injection, mice were anesthetized using isoflurane (maintained at 1.2-1.5 %) mounted on a stereotaxic with a custom-made v-shaped head holder. After the parietal and temporal bone was exposed, the skull above the auditory cortex (2 × 3 mm) was gently drilled and the bone was carefully lifted. The craniotomy was covered with a small cover glass, fixed with dental cement. To position the window plane perpendicular to the objective under the microscope, a custom-made titanium head post was mounted on the frontal bone and embedded with dental cement. After the surgical procedure, animals were single-housed and recovered for at least 1 week before further handling.

### Habituation to awake chronic two-photon imaging

We habituated mice to handling at the 2-photon microscope. Mice were fixated under the objective in a custom-made acrylic glass tube, using a custom-made head post. The mouse head was laterally tilted such that the surface of the auditory cortex aligns with the horizontal plane. Habituation was repeated for at least 7 days, the duration of which increased from around 10 min on the first day to around 2 hours on the last day, comparable to the duration of two-photon imaging in one day. Habituation ended when mice were accommodated to the head fixation apparatus. Since the sound stimulus set for longitudinal two-photon imaging was presented dozens of times each day, animal subjects repeatedly experienced all sound stimuli before data acquisition.

### Intrinsic imaging

We identified the functional organization of auditory fields to set the location of calcium imaging within the fields via intrinsic signal imaging. We followed the protocol previously reported^42^. Briefly, the mesoscopic optical imaging was conducted using a CCD camera with macroscope objectives. Intrinsic signals were recorded from near-infrared (780 nm) LEDs while presenting pure tone pips with different tone frequencies in 30 randomized trials. The change in light reflectance between pre- and post-stimulus image was computed and averaged across trials.

### Sound presentation

Sound design and delivery has been described in the previous study^12^. Briefly, all sounds were delivered free field by a ribbon loudspeaker 25 cm distance to the mouse head in a soundproof booth. The stimulus set consisted of 34 sound stimuli (19 pure-tone (PT) pips (50ms; 2 to 45 kHz separated by a quarter octave) and 15 complex sounds (CS) (70ms)) separated by one-second interval. The complex sounds were generated from arbitrary samples of music pieces or animal calls replayed at fourfold speed. In a subset of mice across all groups a set of 9 conspecific vocalizations (< 80ms) were presented during habituation and during imaging in addition. All stimulus on- and offsets were smoothened by a cosine ramp function with duration of 10 msec, respectively. The stimulus set was presented with 10 repetitions per stimulus in a pseudo-random order for each field of view at 70 dB sound pressure level.

### Two-photon imaging

The imaging configuration and procedure has been described in detail previously^12^. Briefly, two-photon calcium imaging was performed with a commercial microscope (Ultima IV, Bruker Corporation) with a 20×-objective (XLUMPlan Fl, NA = 0.95, Olympus) and a tunable pulsed laser (Chameleon Ultra, Coherent) in a soundproof chamber. Both GCaMP6m and mCherry were co-excited at 940 nm wavelength and separated by emission using a filter cube (U-MSWG2, Olympus). tagBFP was excited at 810 nm and separated by another filter cube (U-MWIB3, Olympus). Time-series imaging was performed using a field of view (FOV) of 367 × 367 μm (pixel size: 256 × 128) at 5 Hz frame rate. In a daily session, imaging was performed not only with presentation of the sound set, but also without any sound presentation to record spontaneous neuronal activities for 144 sec in a given FOV. In the longitudinal imaging experiments, 6-8 FOVs were imaged per mouse in layer 2/3 column (about 110 – 325 μm depth from cortical surface). FOVs with reliable sound responses were repeatedly imaged at a two-day interval, except for the last imaging day (4-day interval between 6^th^ and 7^th^ time point). Imaging days were counted according to the day of laser ablation in ablation experiments. Z-stack images of the layer 2/3 column were also acquired in different timepoints (before starting the experiment, just before and after microablation, and after finishing the imaging experiment).

### Protocol, quantification and validation of microablation

Targeted neurons were identified from a full frame scan. We switched to point scan mode with dwelling time of 8 us, scan position was set at the center of nucleus of the neurons. Microablation was performed with wavelength of 760 nm femtosecond laser pulses (140 fs) delivered through a 0.95 NA, 20× objective in mice that were lightly anesthetized with isoflurane (0.8-1.3%). Laser power was set between 100 – 220 mW according to the quality of fluorescence signal in the target neuron. We applied the pulsed laser and immediately terminated on the detection of the abrupt elevation of Ca^2+^ fluorescence. Otherwise, we kept the laser application for up to 30 sec. If we observed clear elevation of Ca^2+^ signal within a minute after laser exposure, we determined the procedure effective. We took an iterative approach for ablation to increase success rate but prevent off-target damage to the surrounding region at the same time (Extended Data Fig. 3a). If we did not observe such a Ca^2+^ elevation, we repeated the procedure with slight increase of the laser power, at most three times. In case the targeted neuron still does not show any observable increase of Ca^2+^ fluorescence, we waited for a few tens of minutes and again applied the laser once more. When the last application did not induce an elevated Ca^2+^ signal of the neuron, we categorized the neuron into procedures ineffective. As longer exposure times tended to produce off-target effects^44^, whenever we observe abnormal Ca^2+^ increase or strong bleaching in the surrounding region (approx. 50 μm in radius) during the iteration, we halted the laser application and categorized the neuron into procedures ineffective. In case accumulated laser ablation caused massive damage to the surrounding regions (approx. >100 μm in radius) of the target neurons, we terminated the experiment as failure.

We conducted in vivo and ex vivo experiments to evaluate the microablation protocol. To keep tracking neurons across days in vivo, we took a single image of each FOV with size of 367×367 μm (pixel size: 512 × 512) for each imaging session. We evaluated the effect of microablation by taking z-stack image of layer 2/3 column including the targeted neurons and comparing image of nucleus and somatic fluorescence between the time points just before and after microablation, after 2 and 4 days, and quantifying the loss of nucleus and calcium signal for the target neurons. The criteria for successful ablation of the target neurons are basically loss of nucleus signal in the neurons described as follows: a neuron was identified effectively ablated when the neuron demonstrated 1) complete loss of nucleus fluorescence and abnormality (complete loss or excessive filling) of Ca^2+^ somatic fluorescence, or 2) partial loss of nucleus signal with excessively filled Ca^2+^ fluorescence in soma. On the other hand, we classified a neuron as non-effectively ablated when the loss of nucleus fluorescence was unclear or when the loss was partially observed but with intact Ca^2+^ somatic fluorescence.

After 4 days of *in vivo* z-stack imaging, mice were anesthetized, the cranial windows were carefully removed, and the Evans blue (Sigma-Aldrich, E2129-10G) was injected at the corners of the imaged region on the cortical surface, so that the blue dye can serve as the landmark of imaged region after tissue extraction. Mice were perfused with a PBS/Heparin solution and subsequently with a 4% PFA solution following standard procedures. To keep the z-stack imaged volume intact as much as possible, the bulk brain section over 600 μm thickness was cut roughly parallel to the two-photon imaging plane on a vibratome (VT-1000, Leica Biosystems). To enable confocal imaging of the bulk section, the section was cleared with the CUBIC protocol^71^, washed with PBS and incubated in a DAPI solution (10 μg/mL) for 24 hours. Next, Images were acquired on a confocal microscope (DMi6000B CS TCS SP5, Leica) using 20x dry objective (HCX PL APO CS NA = 0.7) with frame size of 553.6 × 553.6 μm (pixel size: 1024 × 1024), as a z-stack with 0.5 μm resolution. The region for confocal imaging was estimated by the blue-dye landmarks. By slightly scaling and rotating roll, pitch, and yaw of the post-hoc 3 D confocal image, the in-vivo imaged population and microablated neurons were identified in the post-hoc image data.

### Validation of the AAV-vector labeling inhibitory interneurons in vivo

rAAV8-mDlx-NLS-tagBFP was injected into GAD67-GFP mice. After three weeks of injection, the mice were deeply anesthetized with Ketamine/Medetomidine (mg/kg), perfused with 10 ml PBS / Heparin and switched to 10 ml 4% PFA solution in the standard protocol. Brains were extracted and sliced for every 100 μm coronal section with Vibratone (VT-1000), then mounted on cover slips. Confocal images were acquired (DMi6000B CS TCS SP5) using 20× dry objective (HCX PL APO CS). From the images, colocalization of GFP and tagBFP signals was evaluated by custom-written MATLAB scripts.

## Data analysis

### Image processing pipeline for chronic two-photon data

Before applying an individual neuron tracking algorithm, global xy-image displacement induced by movement was corrected by cross-correlation based method^72^. Region of interests (ROIs) were semi-automatically selected by a custom-made MATLAB program, which were manually corrected later by a human expert and usually described by a set of several hundred points marking the centers of neuronal nuclei. To track individual ROIs across days, we followed a procedure described in the previous study^12^. Generally, two-dimensional affine transformation was optimally applied to register ROIs in each frame of the time series from the same FOV across several days. Optimization of the transformation was achieved by Nelder-Mead-Simplex algorithm with MATLAB function (fminsearch), and by iterating it in the two different spatial scales which first covered the entire frame and secondly focused on the four equally split image segments to adjust local movements during two-photon scanning. Lastly, to adjust to further local distortion, individual ROIs were allowed to move up to two-pixel (2.87 μm) and find a better matching with the image.

### ROI inclusion criteria and Calculation of ΔF/F_0_, deconvolution

To include only neurons in the analysis that had a reliably present nucleus signal in the H2B:mCherry channel, we applied four quality criteria described in our previous study^12^: Nearest Neighbor Distance, Normalized Soma Signal Intensity, Soma Signal to Noise ratio and Objective Function Value, which were applied on a frame-by-frame basis, so that a neuron was either reliably present or excluded at each time point. To correct fluctuating background fluorescence from somatic Ca^2+^ signal, the out-of-focus neuropil signal was calculated as the average fluorescence value of an area surrounding each individual regions of interest, and subtracted from the time series after multiplication with a contamination ratio as described before^73,74^. We calculated a contamination ratio, independently for each imaging plane in each experiment^73^. To calculate ΔF/F, the baseline F_0_ used to compute the ΔF/F_0_ was defined as a moving rank order filter, the 30th percentile of the 200 surrounding frames (100 before and 100 after). To estimate neuronal firing rate, this ΔF/F_0_ was then deconvolved using the conventional algorithm^75^.

### Stimulus-evoked sound responsiveness of single neurons

To classify single neurons as sound responsive or not, all the 10 trials from a given stimulus were compared in a signed-rank test against ten pre-stimulus spontaneous activity. A neuron was classified as significantly responsive, if the p value was below 0.05 after a Benjamini-Hochberg (or false discovery rate, FDR) correction for multiple comparisons against number of stimuli (34), for at least one stimulus^12^. The amplitude of sound-evoked Ca^2+^ transients was calculated by subtracting the average pre-stimulus baseline ΔF/F value (2 frames corresponding -200-0 ms) from the average post-stimulus peak ΔF/F value (2 frames corresponding 200-400 ms) of all trials^73^. We defined best response amplitude as the average response amplitude across trials elicited by the stimulus giving rise the significant maximal response among the 34 stimuli.

### FOV inclusion criteria

We included FOVs in our analysis that satisfied the following criteria: (A) FOVs needed to contain at least 100 ROIs (i.e. neurons) which fulfilled the quality criteria described above, (B) FOVs needed more than ten significantly sound responsive neurons during baseline days.

### Selection of sound responsive, non-sound responsive and inhibitory neurons for microablation

During the two baseline days before microablation (day 3 and day 5), we identified sound responsive neurons which showed significant sound-evoked response for at least one stimulus in each FOV. For each sound responsive neuron, we calculated the average amplitude across all the stimulus conditions which elicited significant responses. We derived the amplitude distribution of responsive neurons sorted in descending order for each day (day 3 and day 5). Then we created a single distribution after merging and sorting the two days distributions based on their amplitudes. In case identity of a neuron was duplicated because the neuron was responsive on both the days, the higher-rank neuron was selected to have unique neuron identities across the merged distribution. Thus, we prepared the sorted sound-responsive distribution merged for day 3 and day 5 in each FOV, or “the distribution of candidate responsive neurons”. On the day of microablation, we targeted neurons from the top of the distribution of candidate responsive neurons.

We identified non-sound responsive neurons, for day 3 and day 5 respectively, which did not show significant response to any of sounds in each FOV, and derived the amplitude distributions of these neurons sorted by absolute amplitude in ascending order. We selected the neurons responsive on neither of the days, which equal to the neurons existing in both the distributions. We calculated the average rankings of the neurons from both the distributions and sorted again the neurons based on the average ranking in ascending order to prepare “the distribution of candidate non-responsive neurons”. On the day of microablation, we targeted neurons from the highest rank (i.e., smallest absolute amplitude) of candidate non-responsive neurons.

For inhibitory neuron ablation group, we included the sound-responsive inhibitory interneurons based on the same criteria for the sound responsive ablation. To achieve comparable number of neurons to be microablated to the other groups of ablation experiment, we also included interneurons with non-significant but large transient amplitude after sound presentation (apparent sound responsive neurons), in case the number of sound responsive neurons does not reach the prerequisite number for each FOV. These apparent sound responsive neurons were also sorted based on their amplitude averaged across day 3 and day 5, then we selected neurons from the highest amplitudes. Practically in the experiment of microablation of sound responsive, non-responsive or inhibitory responsive neurons, we had to avoid neurons for microablation when the neurons showed unclear nucleus fluorescence due to optical occlusion by blood vessels, weak expression, out-of-focus position. In the end we were able to target 4-8 neurons for microablation in each FOV. Since the targeted neurons in these three experimental cohorts were determined based on either sound responsiveness and/or cell type, the spatial distribution of these neurons was generally random in each FOV.

In the control group, on the day when the animals would have undergone microablation in ablation experiments, mice (N = 7) were set in the microscope chamber under the comparable level of isoflurane anesthesia (∼3 hours) without any additional laser application except for monitoring Ca^2+^ and nucleus fluorescence for tens of minutes (sham procedure). Another 2 mice were set in the microscope chamber, but only for short anesthetic exposure (from half an hour to one hour) to acquire a few z stack images of the FOVs. Our offline analysis confirmed that the control mice exposed to long isoflurane anesthesia on day 6 and the other control mice with short isoflurane exposure on day 6 did not show any distinct difference in average best response amplitude across neurons on day 7 (baseline-normalized amplitude for long anesthetic exposure: 1.072 ± 0.086; for short anesthetic exposure: 1.063 ± 0.047; two-sample t-test, p = 0.96) nor fraction of responsiveness on day 7 (baseline-normalized fraction for long anesthetic exposure: 0.915 ± 0.057; for short anesthetic exposure: 1.114 ± 0.088; two-sample t-test, p = 0.085).

### Exclusion of neurons with unsuccessful ablation and neurons nearby ablated neurons

Targeted neurons include all the neurons to which we applied the laser for ablation. For any microablation experiment, a small proportion of the targeted neurons showed unclear effect of ablation, based on the criteria we described above. Since some of unclearly ablated neurons also demonstrated abnormal Ca^2+^ signals such as lower frequency of Ca^2+^ transients than before, or seizure-like bursting behavior, we excluded all the targeted neurons and nearby neurons for further analysis. Following a previous study^29^, we excluded targeted neurons (regardless of whether microablation was successful or not), as well as neurons within a sphere centered on ablated neurons having a radius of 15 μm, from any analysis.

### Definition of high and low category neurons

In addition, to make a fair comparison between different ablation experiments, we digitally filtered out highly sound responsive neurons and/or non-sound responsive neurons from spared population as ‘filtered condition’ for further analysis. To mimic response property and number of the ablated non-responsive neurons in non-responsive ablation group, we defined five non-responsive neurons with minimum absolute amplitudes per FOV in sound responsive ablation group, which we would have ablated in non-responsive ablation experiment (we call these non-responsive neurons, which were supposed to be physically or digitally excluded, “low category neurons”). Conversely, to mimic response property and number of the ablated sound responsive neurons in sound responsive ablation group, we defined five responsive neurons with highest amplitudes per FOV in non-responsive ablation group (we call these responsive neurons “high category neurons”). Then, we digitally excluded low category neurons from the spared population in sound responsive ablation group, high category neurons in non-responsive ablation group, and both low and high category neurons in control group, respectively. We verified that the number of ablated sound responsive neurons and other high category neurons were comparable between groups (Extended Fig. 4e), that the number of ablated non-responsive neurons and other low category neurons were also comparable (Extended Fig. 4f). We also confirmed that the best response amplitudes of high category neurons had no significant difference between groups (Extended Fig. 4h), the response amplitude of low category neurons was not significantly different (Extended Fig. 4i) and the number of spared neurons in the filtered condition were comparable between the different ablation groups (Extended Fig. 4j).

### Single-neuron tuning correlation and response reliability

For a given sound responsive neurons, the trials were pseudo-randomly split into two sets with the half number of trials. Response amplitudes along the 34 stimuli, i.e., tuning curve were averaged across each half of the trials. Then Pearson correlation of the averaged responses was calculated between the first and the second half set of trials. This procedure was repeated 10 times, and we averaged these correlation values as single-neuron tuning correlation for each day. To evaluate correlation to day 5, instead of calculating Pearson correlation of the averaged responses of half trials on the same day, we calculated correlation between the averaged response of half trials on day 5 and the averaged response of half trials on the other day. We used the same measure for all individual neurons to quantify the reproducibility of stimulus responses between different subsets of trials as single-neuron response reliability across trials^76,77^ in Fig. 3a, 5c, and 6e.

### Representational map

Pearson correlation of sound-evoked response vector among neurons in each FOV to different stimuli was calculated for each trial and averaged across trials for each day^11^. Then, a similarity matrix was constructed where each element in the matrix corresponds to trial-averaged correlation coefficient for each combination of stimuli, excluding pairs with the same trials. Note that each diagonal element, trial-averaged correlation to the same stimulus corresponds to the trial-to-trial reliability of population response to each stimulus. The representational map was generated by averaging all the similarity matrices across all the FOVs in each group of mice for each day. A dimension-reduced representational map was also formed by applying classical Multi-dimensional scaling (MDS), where a similarity matrix was mapped onto 2-dimensional (1st and 2nd) MDS space. Each data point in the space represents the relative distance of the correlation profile in the similarity matrix corresponding to one of the 34 standard stimuli.

### Decoding

A support vector machine a with linear kernel was trained to discriminate each pair of stimuli from the 34 stimuli as a binary classifier, using a built-in function in MATLAB, fitcecoc. In either case that training and testing was done on the same day, or done on different days, cross-validation was performed by 5-fold cross validation. Training and testing the classifier was conducted for a population in each FOV. The decoding performance was defined as the percentage of correctly classified trials, which was averaged across FOVs per mouse and averaged across mice for each experimental group.

### Categorization of neurons based on best response amplitude

Since various single-neuron response properties are considered to be dependent on the response amplitude or the best response amplitude for each neuron, we categorized sound responsive neurons into 4 categories according to their best response amplitudes on a given day (0-15%, 15-25%, 25-55%, 55-1000% in ΔF/F, respectively). Since the best response amplitude is largely log-normally distributed, the largest best amplitude category covers the extensive range in ΔF/F so that in each category roughly the same number of neurons are included for each experimental cohort (mean ± s.d., 25±6.5%, 25±7.4%, and 25±6.7% of neurons for each bin, in sound responsive cohort, non-sound responsive cohort and control cohort, respectively).

### Quantification of tuning width

For a given responsive neurons, the trial-averaged response amplitude across the 34 stimuli, i.e., tuning curve was sorted by descending order (from the largest amplitude for the 1^st^ stimulus to the smallest amplitude for the 34^th^ stimulus) and normalized by the largest amplitude. After categorizing individual neurons into bins according to their best response amplitudes described above, the sorted and normalized tuning curves were averaged for each bin and for each day (Fig. 3e). The tuning width was estimated as the average normalized amplitude at 15^th^ stimulus on a given day (Fig. 3f).

### Simulation for the effect of tuning width on similarity matrix

We simulated how the change in tuning width could affect the similarity matrix to probe what to extent the disturbance of the similarity matrix on day 7 could be derived from the reduction of tuning width. The tuning curves of neurons in each best amplitude bin during baseline days were scaled by multiplying experimentally observed normalized amplitudes across sorted stimuli for each bin of the best amplitude. We constructed again the similarity matrix from neurons with the artificially scaled tuning curves and compared the off-diagonal correlation between the simulated similarity matrix and the experimentally observed similarity matrix on day 7.

### Signal correlation in local population

To quantify similarity in tuning between neurons, we calculated signal correlation, i.e., Pearson correlation of the trial-averaged tuning curves between responsive neurons for each FOV on a given day. The distribution of signal correlation was again categorized into the best response amplitude bins described above according to the average best amplitude of a given neuron pair. For each best amplitude bin, we quantified the fraction of neuron pairs with high signal correlation beyond 0.7 out of all responsive neuron pairs on a given day.

### Balance of general excitation and inhibition levels in the local network

To investigate the balance of network-level activities between excitatory and inhibitory neurons, we calculated the product of the fraction of responsive neurons and the average best response amplitude in each mouse (Extended Data Fig. 8e), as a general estimate of the total amount of excitatory response and inhibtory response, i.e., “total responses” in the local network. The balance was quantified as the ratio of (total response in excitatory neurons)/(total response in excitatory neurons + total response in inhibitory neurons).

### Statistical analysis

All statistical analyses were performed using MATLAB (Mathworks, Natick, MA, USA). Most statistical comparisons were performed the two-sided t-test comparing means within individual mice for two different conditions like baseline days and a post-ablation day after false discovery rate (FDR) correction for multiple comparison to test changes after microablation for each experimental cohort. For comparison between different experimental cohorts in a given day or in a set of days, the two-sided t-test was applied after FDR correction. In all cases, we treated not neurons but mice as independent observations. When classifying individual neurons for sound-evoked responsiveness, since post-stimulus activities across trials were highly non-normally distributed, a signed-rank test was performed after Benjamini-Hochberg correction for multiple comparisons against number of stimuli. For the same reason, to compare the normalized correlation coefficient in the representational maps across the experimental cohorts (Extended Data Fig. 5 and 9), we used a non-parametric pairwise multiple comparisons, Dunn’s test for the different pairs of cumulative distribution of the normalized correlation coefficient, as well as Mann-Whitney U test after FDR correction to compare medians between the cohorts for each tested day. For comparison across the experimental cohorts in the other analyses from Fig. 3 to Fig. 6 in general, we performed permutation test on a given day after microablation by shuffling samples 200 times across the three cohorts and estimating p values for the experimentally observed data for each cohort. Regarding the change in similarity matrix constructed after shuffling neurons across FOVs (Fig. 3g), first we applied permutation test on a given day by shuffling neurons across the three experimental cohorts, randomly selecting the same number of neurons as that in the cohort to be compared and constructing the similarity matrix from the selected neurons. We repeated the procedure 500 times and estimated the distribution of the off-diagonal correlation from the cohort-permuted neurons by the 95% confidence interval (CI). Then, we compared the off-diagonal correlation of the similarity matrices from population vectors shuffled across FOVs in each experimental cohort with the CI from the corresponding cohort. Second, we further tested the change of normalized correlation in sound responsive cohort (or non-sound responsive cohort) from control cohort after ablation. We applied the same permutation procedure, but only with two experimental cohorts for comparison (sound responsive cohort and control, or non-sound responsive cohort and control). Sample sizes were largely comparable between conditions (number of mice between the different cohorts). No statistical tests were applied to determine sample sizes.

## Extended Data Figures

**Extended Data Fig. 1.**
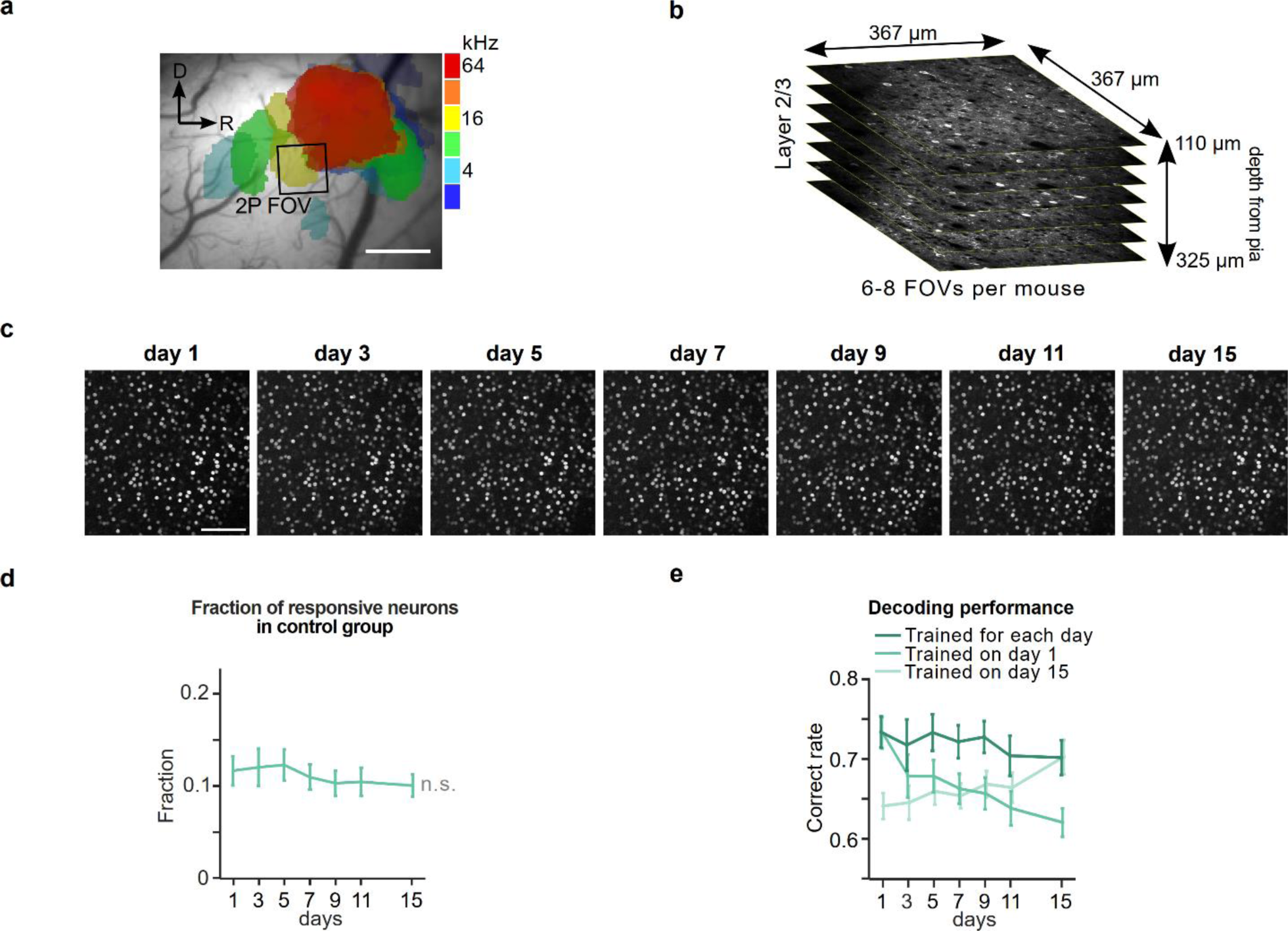
Longitudinal imaging of sound-evoked activity in the mouse auditory cortex. **a**. Example image of intrinsic signals on top of brain surface during burst of pure-tone (2 - 64 kHz) presentation. Intrinsic signals were recorded to functionally identify auditory fields, within which regions for two-photon calcium imaging were subsequently set. In this example, a region for two-photon imaging was selected as a black square. Scale bar: 500 μm. **b**. Schematics of sequential imaging across layer 2/3 column in the auditory cortex. 6 - 8 FOVs were imaged per mouse with average z distance of 24.2 μm (± 5.04 standard deviation). **c**. In vivo two-photon images of H2B::mCherry signal of an example FOV on all the seven imaging days. The distinct labeling in the red channel allows high-fidelity tracking of individual neurons using a signal that is independent of neuronal activity. Scale bar: 100 µm. **d**. Fraction of responsive neurons across days in control group without filtering high and low category neurons. **e**. Linear pairwise discriminability calculated by support vector machine averaged across all possible sound pairs and FOVs (mean ± s.e.m.) plotted across days. The classifier was trained with data from either given (dark green), first (green), or last (light green) imaging day.

**Extended Data Fig. 2.**
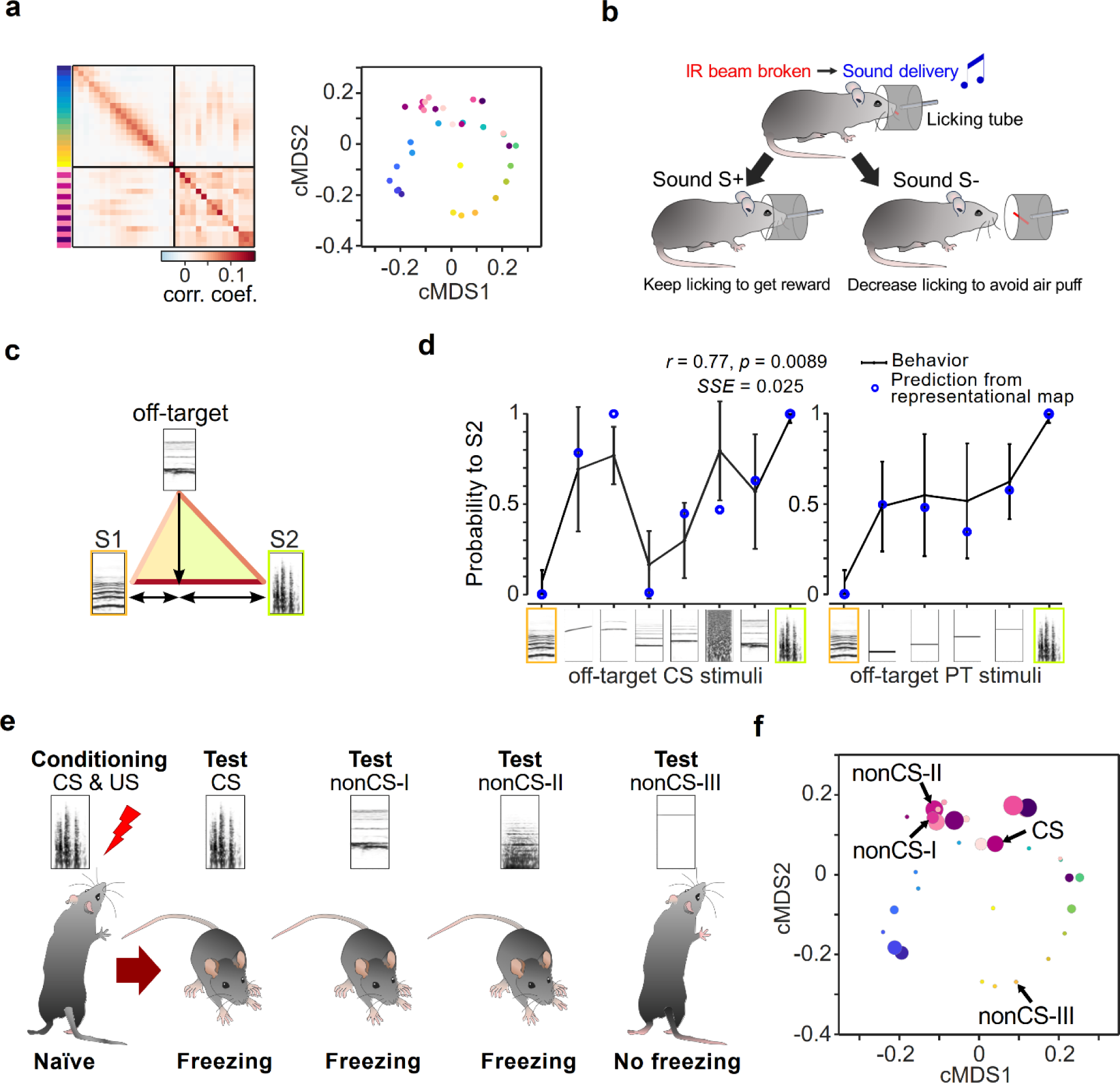
The structure of the representational map in the auditory cortex predicts stimulus generalization in a go/no-go task as well as in classical conditioning. **a**. Grand averaged representational similarity matrix and corresponding dimension-reduced classical Multidimensional scaling (cMDS) display constructed from the dataset acquired in the current study during baseline (n = 29 mice, 3 imaging days). **b**. Schematic of a go/no-go sound discrimination paradigm from a previous study^11^, in which the same set of sound stimuli was used as in the current study. In well-trained mice licking behavior was measured in response to reinforced S+ and S-stimuli. In addition, spontaneous behavioral categorization of non-reinforced off-target stimuli was used as an estimate of perceptual similarity of a given off-target in relation to the pair of target stimuli. **c**. Schematic of the application of the neurometric similarity matrix to estimate perceptual distances. The length of each side in the triangle of off-target, and reinforced stimuli S1 and S2 is determined by the correlation coefficients. The estimated relative representational similarity of the off-target stimulus to S1 or S2 is defined as the internal dividing point in the line connecting S1 and S2, orthogonally mapped from the off-target point. **d**. Solid black line: Behaviorally evaluated perceptual similarity as the go probability of off-target sounds in go no-go behavioral task (Adapted from Bathellier et al., 2012). Blue dots: Prediction of perceptual similarity from the representational similarity matrix in **a** with the metric in **b**. **e**. Stimulus generalization to conditioned and non-conditioned sound stimuli from another previously published study^12^, in which the same set of 34 sound stimuli as in the current study was used. One of the stimuli was used as a conditioned stimulus (CS) in a classical auditory-cued fear conditioning paradigm and was paired with a foot shock (US). Four days after conditioning, a generalization test was performed by presenting the CS and the three nonCS sounds (I, II, and III), without foot shock. The conditioned mice exhibited freezing behavior to CS, nonCS-I nonCS-II, but not to nonCS-III. **f**. Dimension-reduced display of the representational map obtained in the current study shown in **a**. Dot diameters depict a measure of neuronal plasticity (increased likelihood of a given sound stimulus to activate the same neuron assembly as the CS) obtained in the previous study^12^. Note, that the proximity between nonCS stimuli and the CS in the representational map allows us to predict the degree of plasticity in sound responses in the auditory cortex as well as the level of behaviorally measured stimulus generalization.

**Extended Data Fig. 3.**
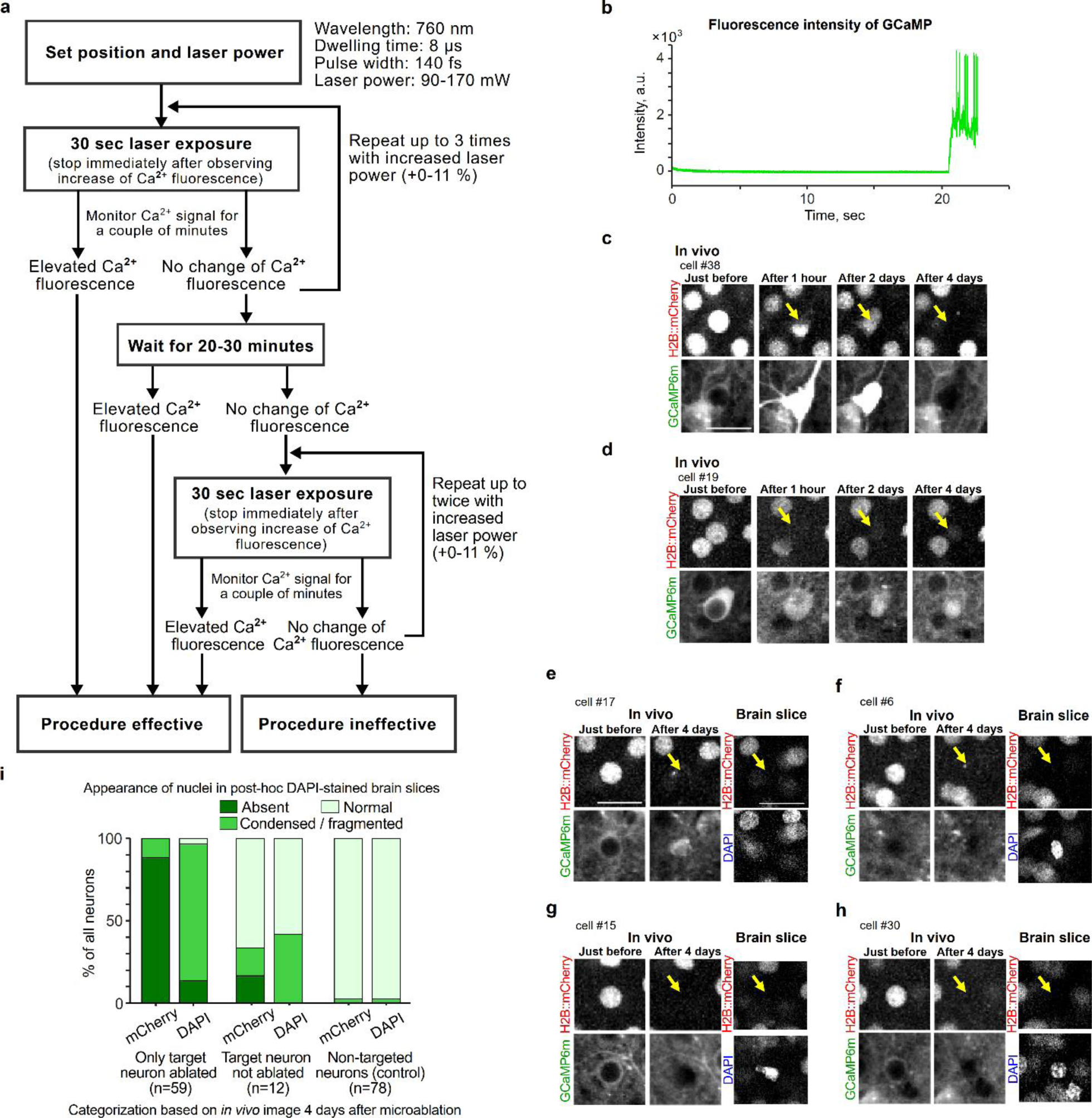
Protocol and validation of the microablation procedure. **a.** Protocol for microablation. See also Methods for further details. **b**. Example trace of Calcium fluorescence during laser exposure on a target neuron when doing point scanning. Ca^2+^ fluorescence was monitored while exposing the laser, and the time-lapse imaging was continued for a couple of seconds to a few tens of second until we observed abrupt elevation of the signal. In this example, abrupt increase of fluorescence intensity was observed around 21 sec from the onset of laser exposure. **c**-**d**. Examples of targeted neurons imaged in vivo over the time course of up to 4 days. In **c**, the targeted neuron showed a permanently elevated fluorescence signal one hour after the microablation procedure. After 4 days, both the nucleus fluorescence and the calcium fluorescence were undetectable. In another example targeted neuron in **d**, after 4 days of the microablation procedure, filled calcium fluorescence remained while the nucleus fluorescence was almost eliminated. When target neurons are successfully microablated, in most cases both signals disappeared (**c**, 89.4 %), in rare cases nucleus signal was lost with filled Ca^2+^ fluorescence (**d**, 10.6 %). **e-h**. Evaluation of the effectiveness of the procedure by re-identifying individual targeted neurons in fixed brain sections counter stained by 4’,6-diamidino-2-phenylindole (DAPI). Four days after the microablation procedure, neurons which lost the H2B::mCherry signal exhibited an absent DAPI signal (**e**) or an abnormally fragmented DAPI morphology (**f**-**h**). **i**. Summary bar plots of nucleus fluorescence signals and post-hoc DAPI signals in targeted neurons categorized based on in vivo images 4 days after the microablation procedure.

**Extended Data Fig. 4.**
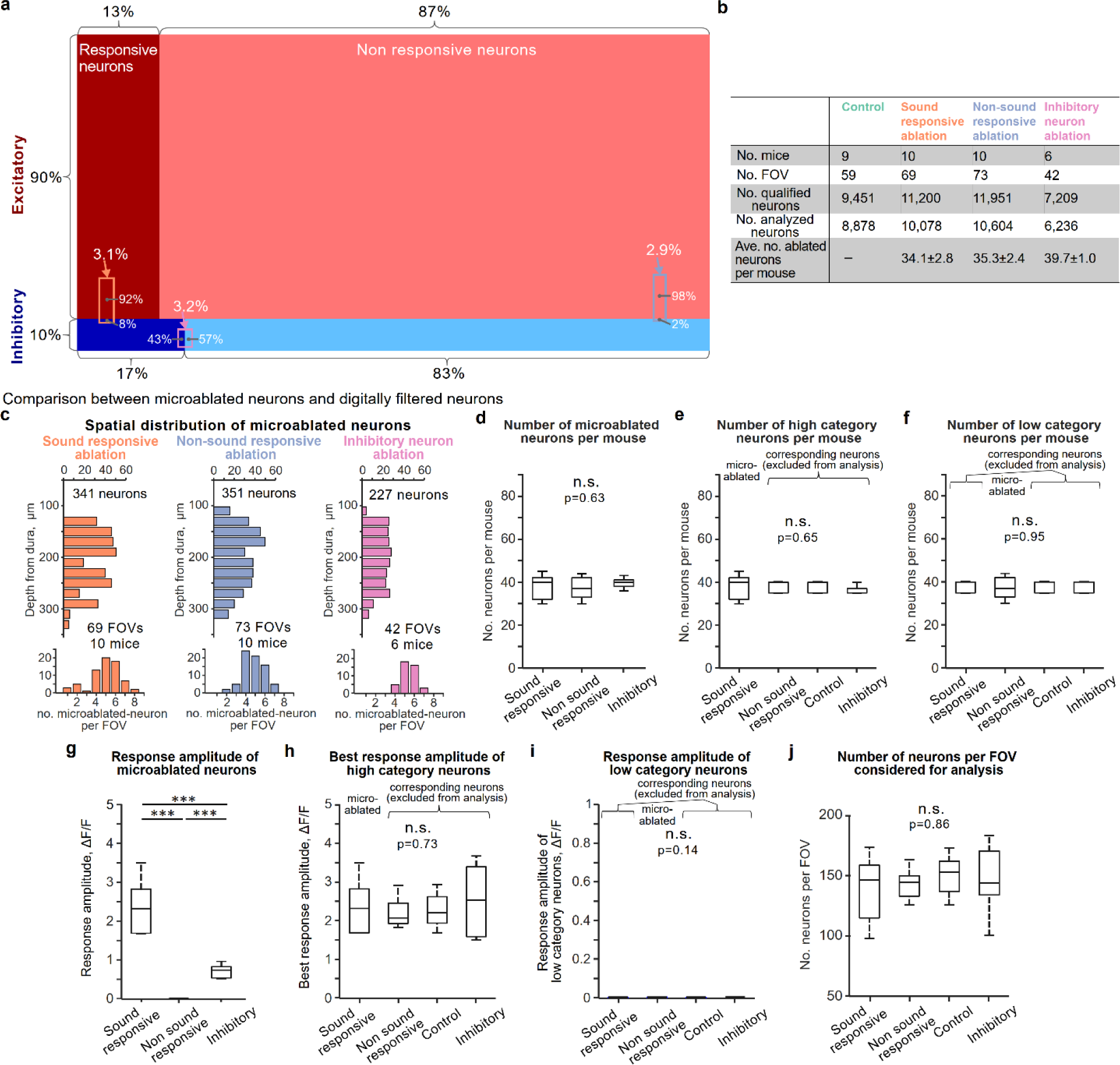
Response statistics of the microablated neurons. **a**. Summary schematic indicating the average fraction of neurons with their categorization of responsiveness and different cell types. For a subset of mice in each experimental group (5/10, 5/10, 7/9 mice in sound responsive cohort, non-sound responsive cohort, control cohort, respectively), we also labeled inhibitory cortical neurons by co-injecting an AAV vector encoding the mDlx enhancer system (AAV-mDlx-NLS-tagBFP) with the AAV vectors expressing GCaMP6m and H2B::mCherry (see also Fig. 5 and Extended Data Fig. 8). Therefore, we were able to identify the excitatory and inhibitory types of microablated and spared neurons. Small rectangles indicate the fraction of microablated neurons for each experimental group. The length of horizontal side refers to the fraction of microablated neurons out of all qualified neurons per FOV. The fraction along the vertical side refers to the ratio of excitatory and inhibitory neurons in the microablated neurons. The categorization of responsiveness is also given for an additional experimental cohort in which inhibitory neurons were microablated (see also Fig. 6). Orange: sound responsive ablation; Blue: non-sound responsive ablation; Pink: inhibitory neuron ablation. The fraction along the horizontal side in the pink rectangle refers to the ratio of significantly responsive and unresponsive neurons in microablated inhibitory neurons per FOV. **b**. Summary table of number of mice, number of FOV, number of neurons and number of averaged microablated neurons per FOV acquired for each experimental group. For sound responsive ablation and non-sound responsive ablation, 41.5 ± 1.75 neurons and 40.3 ± 2.40 neurons were targeted per mouse on average, meaning that 81.9 ± 5.5% and 87.5 ± 2.4% were successfully ablated out of the total number of target neurons respectively, based on the criteria described in Methods. **c**. Spatial distribution of microablated neurons for each experimental group. Top: distribution along depth from dura. Bottom: Distribution of number of microablated neurons per FOV. **d**. Box plots of number of microablated neurons per mouse across experimental groups. One-way ANOVA test, F(2, 23) = 0.47, p = 0.63. **e**. Number of high category neurons per mouse, which means number of microablated neurons for sound responsive ablation and number of corresponding neurons for the other groups. One-way ANOVA test, F(3, 30) = 0.56, p = 0.648. **f**. Number of low category neurons per mouse, which means number of microablated neurons for non-sound responsive ablation and number of corresponding neurons for the other groups. One-way ANOVA test, F(3, 30) = 0.12, p = 0.95. **g**. Response amplitude of microablated neurons averaged during baseline days (3 and 5). The amplitude of non-sound responsive neurons was defined as the average amplitude across stimuli for each neuron. t-test with FDR correction between sound responsive ablation vs. non-responsive ablation: p < 0.001; sound responsive ablation vs. inhibitory neuron ablation: p < 0.001; non-responsive ablation vs. inhibitory neuron ablation: p < 0.001. **h**. Best response amplitude of high category neurons during baseline days. One-way ANOVA test, F(3, 30) = 0.44, p = 0.73. **i**. Response amplitude of low category neurons during baseline days. The amplitude of non-sound responsive neurons was defined as the average amplitude across stimuli for each neuron. One-way ANOVA test, F(3, 30) = 1.98, p = 0.14. **j**. Number of spared neurons per FOV considered for analysis. One-way ANOVA test, F(3, 31) = 0.25, p = 0.86.

**Extended Data Fig. 5.**
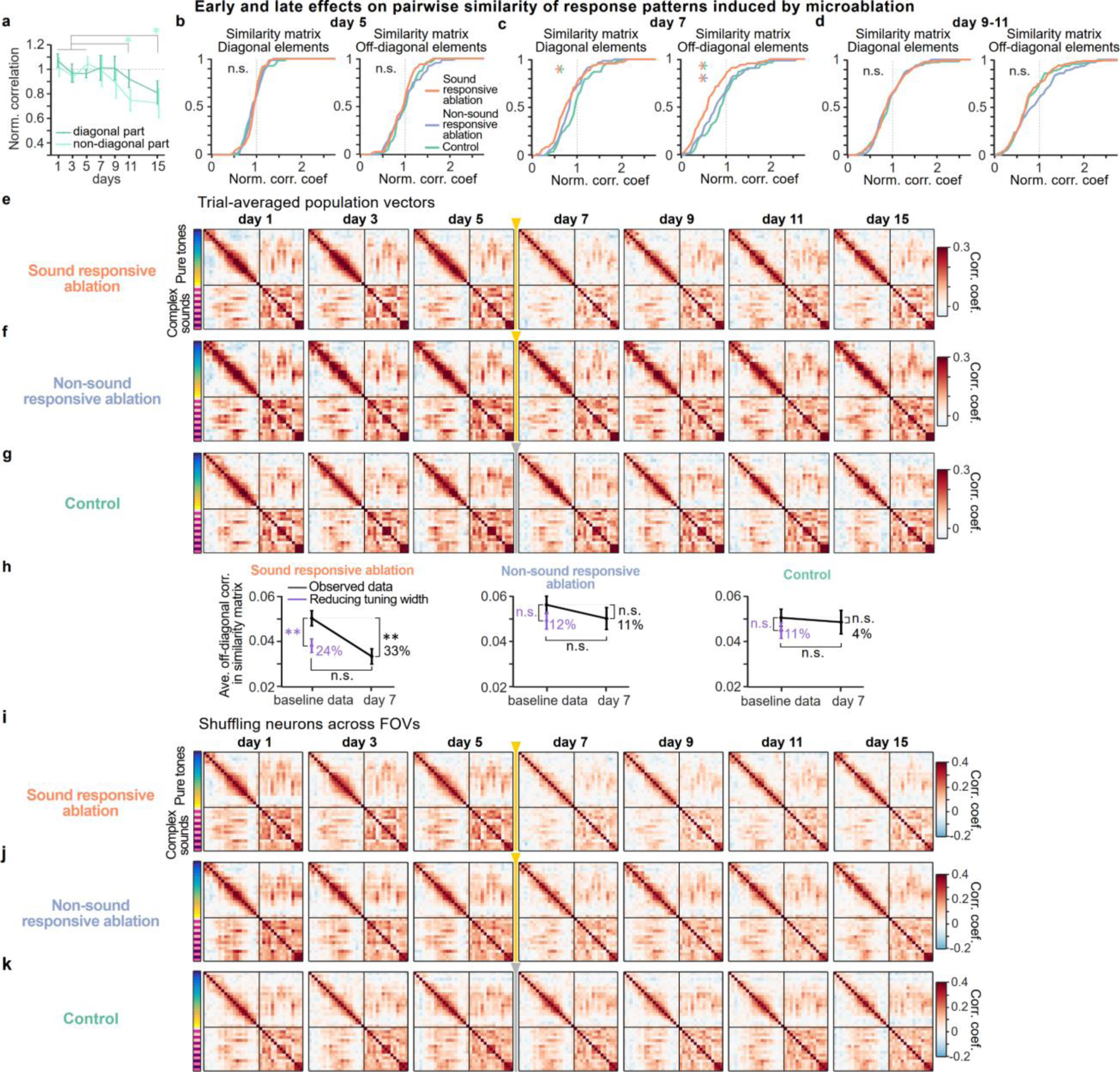
Additional analyses of the effects on the representational maps induced by microablation. **a**. Normalized correlations averaged across diagonal elements (dark green) and off-diagonal elements (light green) in the similarity matrices from the filtered data without the high and low category neurons. Two-sample t-test between baseline days vs. days after ablation with FDR correction, in average of diagonal elements: p = 0.926, p = 0.926, p = 0.534, p = 0.0838, for day 7, 9, 11, 15, respectively; average of off-diagonal elements: p = 0.882, p = 0.277, p = 0.0072, p = 0.0072 for day 7, 9, 11, 15, respectively. **b**. Cumulative distribution of baseline normalized correlation coefficient along diagonal elements (left) and along off-diagonal elements (right) on day 5 in the representational map for each experimental group. To make sure that the distributions of normalized correlation in the representational map during baseline were comparable between the experimental groups, difference in these cumulative distributions was tested by Dunn’s test: Q_critical_ = 2.39, all Q values between groups for both diagonal and off-diagonal components in the similarity matrices, Q < Q_critical_. Mann-Whitney *U* test between groups with FDR correction for both the diagonal and the off-diagonal components, all p values, p > 0.5. **c**. Cumulative distribution of baseline normalized correlation coefficient on day 7 in the representational map. Difference in these cumulative distributions between the experimental groups was again tested by Dunn’s test as well as Mann-Whitney *U* test. The cumulative distribution in sound responsive ablation cohort significantly relatively shifted to lower correlation than the other groups: For diagonal components, Q_critical_ = 2.39; Sound responsive ablation vs. Control: Q = 2.93; Sound responsive ablation vs. Non-responsive ablation: Q < Q_critical_; Non-responsive ablation vs. Control: Q < Q_critical_; Mann-Whitney *U* test with FDR correction between Sound responsive ablation vs. Control: p=0.016; Sound responsive ablation vs. Non-responsive ablation: p = 0.077; Non-responsive ablation vs. Control: p = 0.20; For off-diagonal components, Sound responsive ablation vs. Control: Q = 3.44; Sound responsive ablation vs. Non-responsive ablation: Q = 2.57; Non-responsive ablation vs. Control: Q < Q_critical_; Mann-Whitney *U* test with FDR correction between Sound responsive ablation vs. Control: p = 0.0018; Sound responsive ablation vs. Non-responsive ablation: p = 0.015; Non-responsive ablation vs. Control: p = 0.33. **d**. Cumulative distribution of baseline normalized correlation coefficient during day 9-11 in the map. Dunn’s test: Q_critical_ = 2.39, all Q values between groups for both diagonal and off-diagonal components in the similarity matrices, Q < Q_critical_. Mann-Whitney *U* test between groups with FDR correction for both the diagonal and the off-diagonal components, all p values, p > 0.08. **e**-**g**. Similarity matrices from trial-averaged population response vectors across days in sound responsive ablation condition, non-sound responsive ablation condition and control, respectively. **h**. Simulated off-diagonal correlation of similarity matrix by experimentally derived downscaled tuning curve in individual neurons. Two-sample t test between baseline correlation vs. simulated correlation after downscaling, p = 0.0083 for sound responsive cohort; p = 0.190 for non-sound responsive cohort; p = 0.270 for control; Two-sample t test between simulated correlation after downscaling vs. correlation on day 7: p = 0.419, p = 0.888, p= 0.575 for sound responsive, non-sound responsive, and control cohort, respectively. **i-k**. Similarity matrices across days from population response vectors after shuffling neurons across FOVs for each experimental cohort.

**Extended Data Fig. 6.**
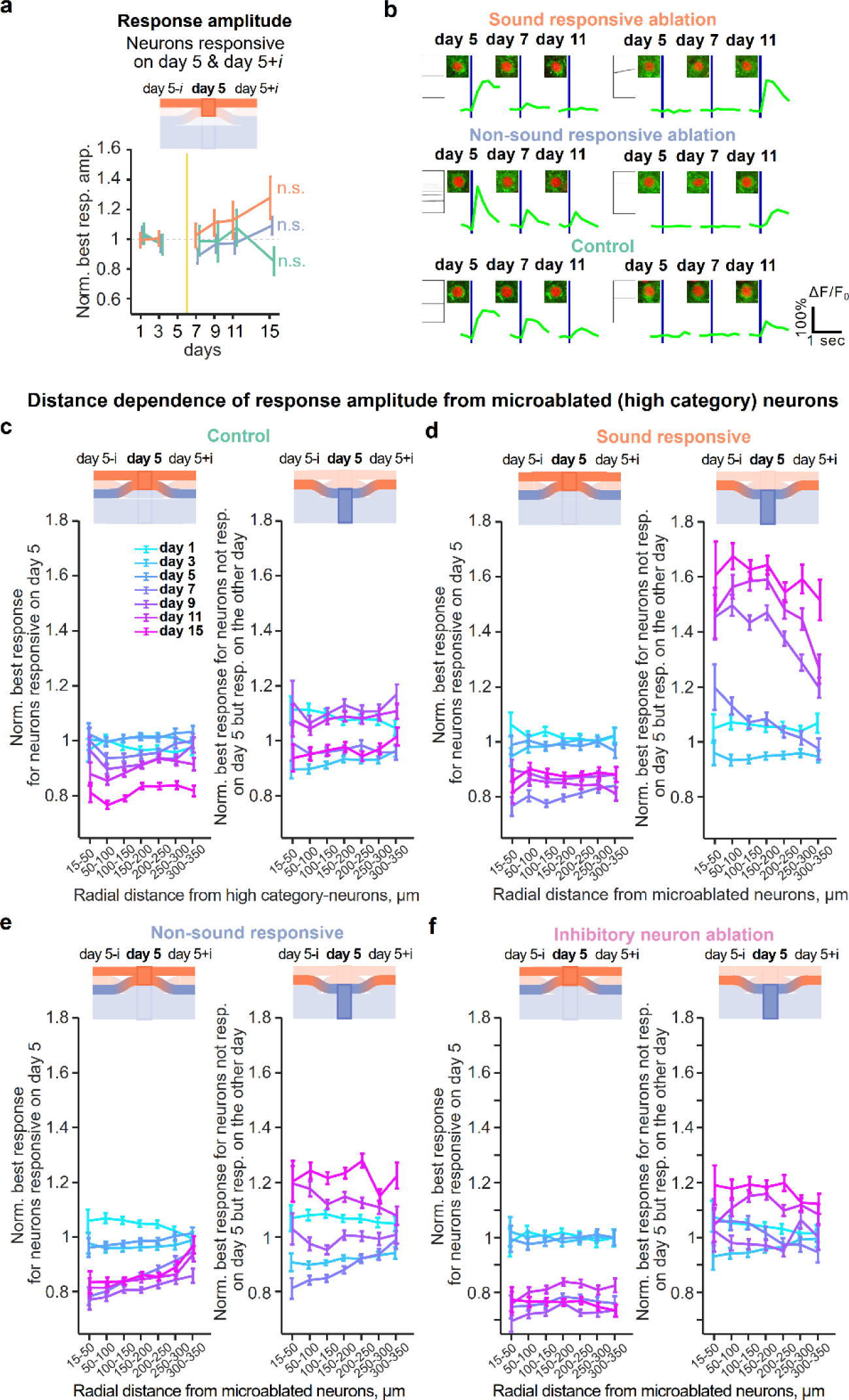
Additional analyses of the microablation-induced effects on response amplitudes. **a.** Best response amplitude of neurons responsive on both day 5 and day 5±i normalized by the average best response during early baseline days (1 and 3). One-way ANOVA test, sound responsive ablation: F(5, 54) = 1.29, p = 0.28; Non-responsive ablation: F(5, 54) = 1.35, p = 0.26; Control: F(5, 48) = 0.58, p = 0.72; **b.** Exemplary sound-evoked Ca^2+^ transients of neurons which responded to sounds on day 5 and reduced the amplitudes in these responses in following days after ablation in sound responsive ablation (top left), non-sound responsive ablation (middle left), and control (bottom left), respectively. Right: Exemplary sound-evoked Ca^2+^ transients of neurons unresponsive on day 5, but becoming responsive in the following days after ablation. **c**. Assessing the spatial extent of the microablation-induced effect on normalized best response amplitudes in the spared neurons. In the control group, the high category neurons with strong sound-evoked responses that were excluded from the further analysis were considered as reference (see Methods). Spared neurons were split into neurons responsive on day 5 (Left) and newly responsive neurons (Right). In a given mouse, the three-dimensional distance of a spared neuron to each of the 30-40 microablated neurons (or high category neurons in the control group) was calculated, i.e. contributed multiple distance measurements associated with the same effect on the best response amplitude. All distance combinations were categorized into distance bins ranging between 15 and 350 μm. For each distance bin, the corresponding best response amplitude in each neuron was normalized to the average baseline amplitude for each mouse, the normalized best response was averaged across neurons (error bars are shown as s.e.m.). In the control group, normalized best responses of neurons responsive on day 5 gradually decreased over days with generally constant amplitude across distance, but exhibited modest reduction of normalized amplitude on later (day 11 and 15) days (Two-sample t-test between short and long distance, day 1 to day 15: p = 0.129, p = 0.823, p = 0.138, p = 0.0486, p = 0.0612, p = 0.0014, p = 0.0012). Normalized best responses of newly responsive neurons remained rather flat across distance and did not show any systematic change during and after baseline days (Two-sample t-test between short and long distance, day 1, p = 0.0656; day 3, p = 0.014; day 7, p = 0.957; day 9, p = 0.127; day 11, p = 0.0797; day 15, p = 0.194). **d**-**f**. Same as **c**, but from microablated neurons for sound responsive ablation, for non-responsive ablation, and for inhibitory neuron ablation, respectively. In sound responsive ablation, normalized best responses of neurons responsive on day 5 decreased over days, but with constant amplitude across distance (Two-way ANOVA, F(6, 244429) = 0.451, p = 0.845 across distance bins; F(6, 244429) = 151.58, p < 1.0×10^-10^ across days). On the other hand, normalized best responses of newly responsive neurons strongly increased the amplitudes later days (Two-way ANOVA, F(6, 121590) = 10.9, p = 3.56×10^-12^ across distance bins; F(5, 121590) = 543.19, p < 1.0×10^-12^ across days). Three days after microablation, the normalized amplitude with nearby distance exhibited larger amplitude than the amplitude distant from ablated neurons (Two-sample t-test between short (< 100 μm) and long distance (250-350 μm), day 1, p = 0.488; day 3, p = 0.419; day 7, p = 0.0001; day 9, p < 1.0×10^-4^; day 11, p = 0.0002; day 15, p = 0.124). For non-sound responsive ablation, normalized best responses of neurons responsive on day 5 decreased over days and the reduction was more prominent at shorter distance from ablated neurons (Two-way ANOVA, F(6, 284429) = 9.47, p = 1.99×10^-10^ across distance bins; F(6, 284429) = 230.36, p < 1.0×10^-10^ across days; Two-sample t-test between short and long distance, day 1, p = 0.0116; day 3, p = 0.495; day 5, p = 0.0313; day 7, p < 1.0×10^-4^; day 9, p = 0.0005; day 11, p = 0.0003; day 15, p = 0.0004). On the other hand, normalized best responses of newly responsive neurons did not change over distance, but showed slight increase in amplitude later days after ablation (Two-way ANOVA, F(6, 146974) = 1.85, p = 0.0853 across distance bins; F(5, 146974) = 274.42, p = 1.0×10^-10^ across days). The normalized amplitude on day 7 was suppressed at shorter distance, but conversely increased later days (Two-sample t-test between short and long distance: day 1, p = 0.26; day 3, p = 0.0359; day 7, p < 1.0×10^-5^; day 9, p = 0.63; day 11, p = 0.0122; day 15, p = 0.0846).

**Extended Data Fig. 7.**
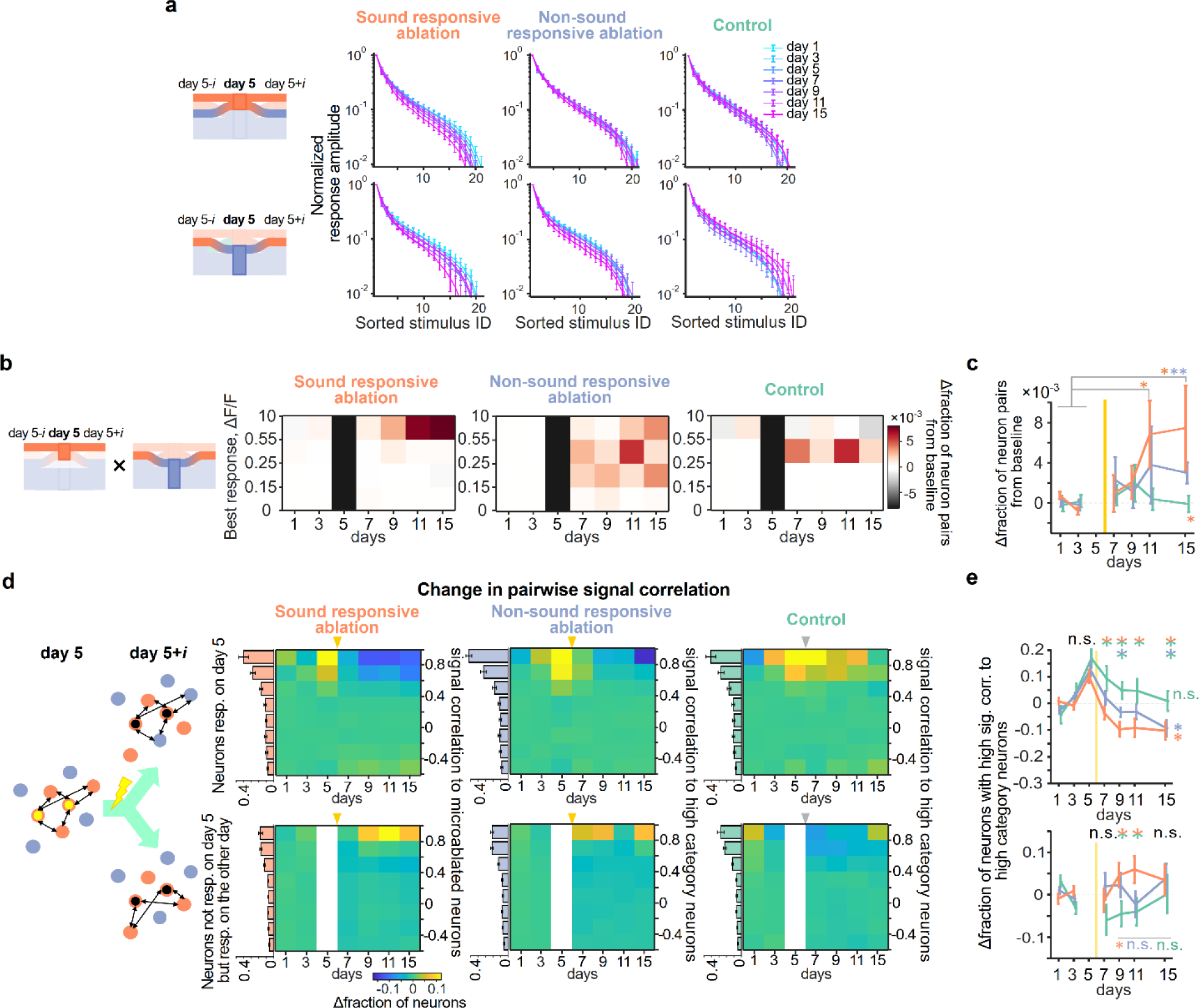
Additional analyses of single-neuron response dynamics contributing to the change of the representational maps. **a**. Same as Fig. 3e bottom, but normalized tuning curves of day 5 responsive neurons (top) and of neurons unresponsive on day 5 but responsive on the other day (bottom), for sound responsive cohort (left), non-sound responsive cohort (middle) and control cohort (right), respectively. **b**. Same as Fig. 4c, but the colormaps displaying the change in the fraction of high-signal-correlation neuron pairs, one of which was from neurons responsive on both day 5 and any other day, and the other of which was from neurons unresponsive on day 5 but responsive on any other day. **c**. Change in the fraction of neuron pairs with high signal correlation from baseline at the largest response amplitude bin in the corresponding colormaps in **b**. Two-sample t-test between baseline days and days after ablation with FDR correction: p = 0.243, p = 0.0667, p = 0.0146, p = 0.0175 for day 7, 9, 11, 15 in sound responsive cohort; p = 0.120, p = 0.188, p = 0.120, p = 0.0055 for day 7, 9, 11, 15 in non-sound responsive cohort; p > 0.398 for all post-sham ablation days in control cohort. Permutation test for group comparison: p > 0.05 from day 7 - 11 for all the three cohorts, but p < 0.05 on day 15 for sound responsive cohort. **d**. Left: Scheme of signal correlation between microablated neurons and spared neurons, pairs of which are defined based on the responsiveness on day 5 and day 5±i. Right: Colormaps show baseline-subtracted fraction of neurons with signal correlation between microablated neurons (or high category neurons except for the sound responsive cohort) and spared neurons, which are responsive on day 5 (top) and not responsive on day 5 but responsive on the other day (bottom). Bar plots next to the colormap are fractions of neurons for signal correlation bins averaged across baseline days (day 1 and day 3) in each split group. **e**. Baseline normalized change in fraction of spared neurons with high signal correlation toward high category neurons. One-way ANOVA, sound responsive ablation: F(5,114) = 3.58, p = 0.0048; non-responsive ablation: F(5,114) = 3.7, p = 0.0039), the fraction did not reduce at all for control group (One-way ANOVA, F(5,101) = 1.44, p = 0.21, comparison across groups by two-sample t-test with FDR correction between responsive ablation vs. control on day 7, p = 0.042; day 9, p = 0.0051; day 11, p = 0.049; day 15, p = 0.061; responsive ablation vs. non responsive ablation on day 7, p = 0.15; day 9, p = 0.057; day 11, p = 0.18; day 15, p = 0.85; non-responsive ablation vs. control on day 7, p = 0.16; day 9, p = 0.047; day 11, p=0.18; day 15, p = 0.061; For newly responsive neurons, comparison across groups by one-sample t-test with FDR correction between responsive ablation vs. control on day 7, p = 0.21; day 9, p = 0.055; day 11, p = 0.049; day 15, p = 0.43; responsive ablation vs. non-responsive ablation on day 7, p = 0.21; day 9, p = 0.29; day 11, p = 0.054; day 15, p = 0.47; non-responsive ablation vs. control on day 7, p = 0.15; day 9, p = 0.086; day 11, p = 0.35; day 15, p = 0.43.

**Extended Data Fig. 8.**
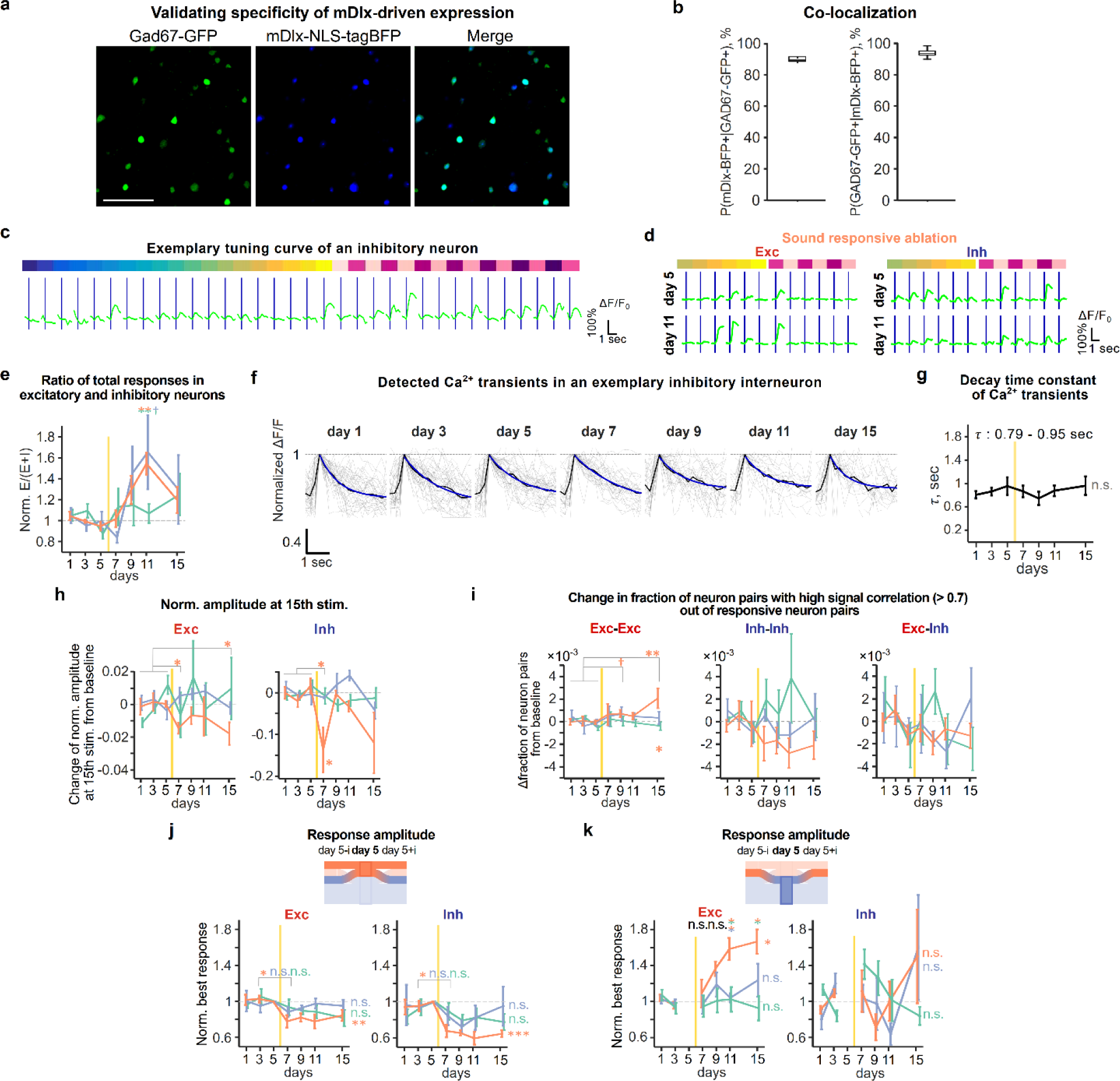
Validation of the viral construct labeling inhibitory neurons and additional analyses of the microablation-induced effects on excitatory and inhibitory neurons. **a**. Representative example of colocalization between mDlx-driven tagBFP expression and Gad67. Scale: 100 μm. **b**. Quantification of colocalization. The fraction of interneuron labeled with the blue fluorescent protein marker out of the entire quality qualified neurons was around 10% for each group (10.5 ± 0.5% (mean ± s.e.m. across mice) in sound responsive ablation, 9.8 ± 0.6% in non-sound responsive ablation, and 7.8 ± 0.4% in control, respectively), in line with previous works with auditory cortex in mature adult mice^82^. When we revisited the profile of cell types in microablated neurons, nearly all the ablated neurons were excitatory neurons (92.3 ± 2.0% for sound-responsive ablated neurons, 98.4 ± 0.9% for non-sound responsive ablated neurons), the fraction of which is higher than the average fraction of excitatory neurons in the population. **c**. Exemplary tuning curve of inhibitory neuron. **d**. Representative excitatory (left) and inhibitory (right) neurons, which change response amplitude to sound stimuli from day 5 to day 11, in sound responsive ablation. **e**. Baseline-normalized ratio of the total sound-evoked activity in excitatory and inhibitory neurons (see Methods). Data presented as mean ± s.e.m. across mice. Two-sided t-test with FDR adjusted p-values, sound responsive cohort vs. control, non-sound responsive cohort vs. control and sound responsive cohort vs. non-sound responsive cohort on day 7: p > 0.5 for all combinations; day 9: p = 0.039, p = 0.039, p = 0.16; day 11: p = 0.045, p = 0.032, p = 0.27; day 15: p > 0.5 for all combinations, respectively; the normalized ratio of best amplitudes on day 15, 1.09 ± 0.088 for sound responsive ablation, 1.10 ± 0.11 for non-sound responsive ablation. **f**. Traces of detected calcium transients of an inhibitory neuron over days. We verified that the changes of response amplitude in inhibitory neurons after microablation (Fig. 5e) were not due to another feature like Ca^2+^ kinetics in somas of inhibitory neurons characterized by the decay time constant^83^. **g**. Decay time constant of calcium transients of all identified interneurons over days across mice (mean ± s.e.m.). One-way ANOVA test, F(6, 28) = 0.48; p = 0.82. **h**. Same as Fig. 3f, but change of tuning width by using normalized response amplitudes at 15th largest stimulus index in the tuning curve over days for excitatory (left) and inhibitory (right) neurons. Two-sample t-test of Δnormalized amplitude for excitatory neurons between baseline days vs. days after ablation with FDR correction. Sound responsive cohort: p = 0.0106, p = 0.253, p = 0.346, p = 0.0106, for day 7, 9, 11, 15, respectively; non-sound responsive cohort: p > 0.474 for all post-ablation days; Control: p > 0.67 for all post-ablation days. For inhibitory neurons, sound responsive cohort: p = 0.213, p = 0.88, p = 0.330, p = 0.0717 for day 7, 9, 11, 15, respectively; non-sound responsive cohort: p > 0.20 for all post-ablation days; Control: p > 0.24 for all post-ablation days. **i**. Change in fraction of neuron pairs with high signal correlation and with large response amplitude from baseline among excitatory-excitatory neurons (left), inhibitory-inhibitory neurons (middle), and excitatory-inhibitory neurons (right). Two-sample t-test of Δfraction of excitatory-excitatory neuron pairs with FDR correction in sound responsive cohort: p = 0.15, p = 0.051, p = 0.11, p = 0.0034 for day 7, 9, 11, 15; non-sound responsive cohort: p > 0.29 for all post-ablation days; control: p > 0.43 for all post-ablation days. Permutation test for group comparison: p > 0.05 for day 7, 9, 11, p < 0.025 on day 15 in sound responsive cohort; p > 0.05 for all post-ablation days in both non-sound responsive and control cohorts. For inhibitory-inhibitory neuron pairs, sound responsive cohort: p > 0.15 for all post-ablation days; non-sound responsive cohort: p > 0.28 for all post-ablation days; control: p > 0.16 for all post-ablation days. For excitatory-inhibitory neuron pairs, sound responsive cohort: p > 0.25 for all post-ablation days; non-sound responsive cohort: p > 0.30 for all post-ablation days; control: p > 0.26 for all post-ablation days. **j**. Baseline normalized best response amplitude of neurons responsive on day 5 in excitatory (left) and inhibitory (right) neurons for the three experimental groups. When splitting excitatory and inhibitory neurons based on the responsiveness on day 5, as observed in Fig. 4b, the normalized best response amplitude of neurons responsive on day 5 significantly decreased on day 7 for both the excitatory and inhibitory neurons in the sound responsive ablation cohort, but not in the other cohorts (One-way ANOVA across days, Excitatory neurons in sound responsive ablation, F(6, 28) = 4.21, p = 0.0038; non-responsive ablation, F(6, 28) = 1.01, p = 0.43; control, F(6, 42) = 1.36, p = 0.25; Inhibitory neurons in sound responsive ablation, F(6, 21) = 7.44, p = 2.3×10^-4^; non-responsive ablation, F(6, 28) = 0.51, p = 0.80; control, F(6, 42) = 1.02, p = 0.42; Two-sample paired t-test of the normalized best amplitude of excitatory neurons between day 3 vs. day 7: sound responsive ablation, p = 0.023; non-responsive ablation, p = 0.34; control, p = 0.38; inhibitory neurons between day 3 vs. day 7: sound responsive ablation, p = 0.049; non-responsive ablation, p = 0.38; control, p = 0.59). **k**. Same as **j**, but for newly responsive neurons. For the neurons unresponsive on day 5 but being responsive on the other day, the normalized best response amplitude of excitatory neurons exhibited a delayed but substantial increase after microablation in the sound responsive ablation (One-way ANOVA across days, F(5, 24) = 7.97, p = 1.5×10^-4^) like Fig. 4b right, but inhibitory neurons did not (F(5, 24) = 1.01, p = 0.43). For the non-responsive ablation and the control cohorts, both the excitatory and inhibitory neurons did not change the normalized best response amplitude over days (Excitatory neurons in non-responsive ablation: F(5, 24) = 1.12, p = 0.37; control: F(5, 36) = 0.28, p = 0.92; Inhibitory neurons in non-responsive ablation: F(5, 23) = 1.18, p = 0.35; control: F(5, 25) = 1.44, p = 0.24). Group comparison revealed that the normalized amplitude of the newly responsive excitatory neurons in sound responsive ablation was significantly larger than those in non-responsive ablation and in control late after microablation (Two-sample t-test with FDR correction, day 7 and day 9: p > 0.2 for either combination of the experimental groups; day 11: sound responsive ablation vs. non-responsive ablation, p = 0.011; sound responsive ablation vs. control, p = 0.024; non-responsive ablation vs. control, p = 0.93; day 15: sound responsive ablation vs. non-responsive ablation, p = 0.14; sound responsive ablation vs. control, p = 0.011; non-responsive ablation vs. control, p = 0.19).

**Extended Data Fig. 9.**
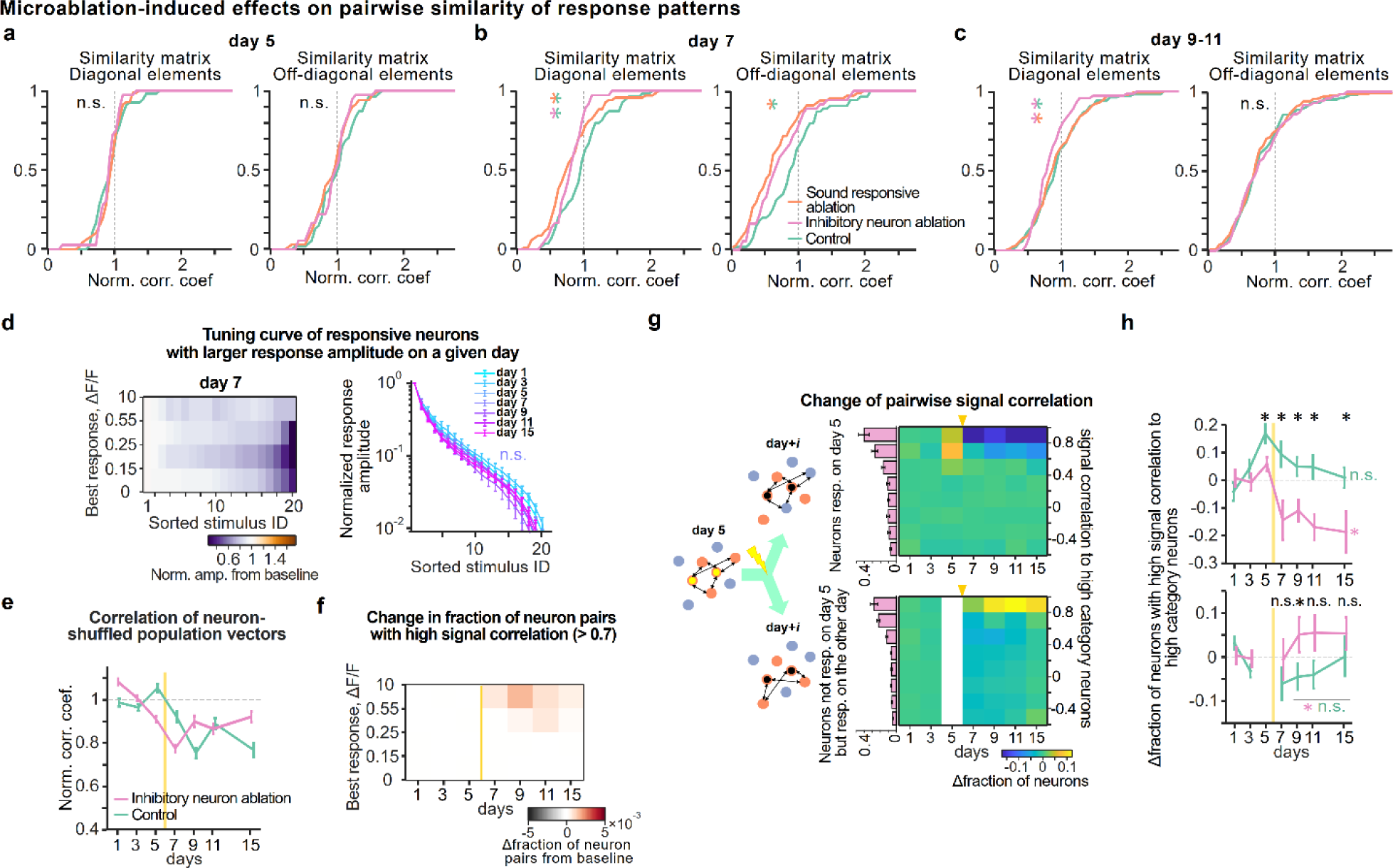
Additional analyses of the effects induced by selective microablation of inhibitory neurons on the representational map and response characteristics of individual neurons. **a**. Cumulative distribution of baseline normalized correlation coefficient on day 5 in the representational map for each experimental group. In inhibitory neuron ablation experiment, 91.7 ± 1.5% was successfully ablated out of the targeted neurons, based on the criteria we already described in Methods. Same as Extended Data Fig. 5a, to compare the effect of ablation between inhibitory neuron ablation and sound responsive ablation groups on the day after microabtaion, difference in these cumulative distributions was tested by Dunn’s test as well as Mann-Whitney *U* test with FDR correction. These cumulative distributions on day 5 were not statistically different by definition between the groups (Dunn’s test for all group combinations in both diagonal and off-diagonal elements, Q < Qcritical; Mann-Whitney *U* test, adjusted p values > 0.5). **b**. Same as **a**, but the cumulative distributions of normalized correlation on day 7. Especially for diagonal elements, the distributions in both sound responsive ablation and inhibitory neuron ablation cohort shifted toward lower correlation than control (Mann-Whitney *U* test, adjusted p-values, sound responsive ablation vs. control, p = 0.016; inhibitory vs. control, p = 0.020; sound responsive ablation vs. inhibitory, p = 0.51; Dunn’s test, sound responsive ablation vs. control, Q = 3.00 (> Qcritical); inhibitory neuron ablation vs. control, Q < Qcritical). For off-diagonal elements, the distributions in sound responsive ablation cohort was lower than that in control (Mann-whitney *U* test, with adjusted p values, sound responsive ablation vs. control, p = 0.0018; inhibitory neuron ablation vs. control, p = 0.06; p = 0.29; sound responsive ablation vs. inhibitory neuron ablation; Dunn’s test, sound responsive ablation vs. control, Q = 3.48 (> Qcritical); Q < Qcritical for the other group comparison). **c**. Same as **a**, but during day 9-11. On the other hand, late after microablation (day 9-11), while the distribution in sound responsive ablation went back to baseline level, the distribution in inhibitory neuron ablation, especially the diagonal elements maintained lower correlation than the other groups (Dunn’s test, inhibitory neuron ablation vs control, Q = 2.44 (> Qcritical); sound responsive ablation vs. inhibitory neuron ablation, Q = 2.43 (> Qcritical); Mann-Whitney *U* test, adjusted p-values, sound responsive ablation vs. control, p = 0.87; inhibitory neuron ablation vs. control, p = 0.056; sound responsive ablation vs. inhibitory neuron ablation, p = 0.056). **d**. Same as Fig. 3e, but normalized tuning curve of responsive neurons in inhibitory neuron ablation. Left: the colormap of the baseline normalized tuning curve on day 7 according to the best response amplitude bins. Right: normalized tuning curves overlaid across days. Two-sample t test of average of normalized amplitude across the stimulus with 2nd largest amplitude to the last stimulus between baseline days vs. day 7: p = 0.202. **e**. Same as Fig. 3g, but normalized off-diagonal correlations in the similarity matrix constructed from the population vectors shuffled across FOVs in inhibitory neuron ablation. Two-sample t test of normalized correlation during all post-ablation days between inhibitory neuron ablation and control. p = 0.33. **f**. Same as Fig. 3h, but the colormap of the change in fraction of responsive neuron pairs with high signal correlation from baseline according to the best response amplitude in inhibitory neuron ablation cohort. Two-way ANOVA, F(6, 154) = 2.05, p = 0.0622 across days; F(3, 154) = 5.097, p = 0.0022 across amplitude bins. **g**. Same as Extended Data Fig. 6f, but baseline-subtracted fraction of neurons with signal correlation between high category neurons and spared neurons in inhibitory neuron ablation condition. **h**. Same as Extended Data Fig. 6g, but baseline-subtracted change in fraction of spared neurons with high signal correlation toward high category neurons. Change in fraction of spared neurons with high signal correlation toward high category neurons in inhibitory neuron ablation and control groups, where spared neurons were split into those responsive on day 5 (top) and those unresponsive on day 5 but being responsive on the other day (bottom). For spared neurons responsive on day 5, the fraction of neurons with high signal correlation in inhibitory neuron ablation was relatively lower than control (Top, comparison between inhibitory neuron ablation vs. control on day 5, p = 0.025, day 7, p = 0.0061, day 9, p = 0.0053, day 11, p = 0.0028, day 15, p = 0.018). On the other hand, for spared neurons newly responsive on the other day, the fraction of neurons with high signal correlation in inhibitory neuron ablation was higher than control (Bottom, two-sample t-test between baseline days vs. days during 9 - 15, p = 0.032 in inhibitory neuron ablation, p = 0.31 in control, comparison between inhibitory neuron ablation vs. control on day 7, p = 0.35; day 9, p = 0.028; day 11, p = 0.072; day 15, p = 0.24).

**Extended Data Fig. 10.**
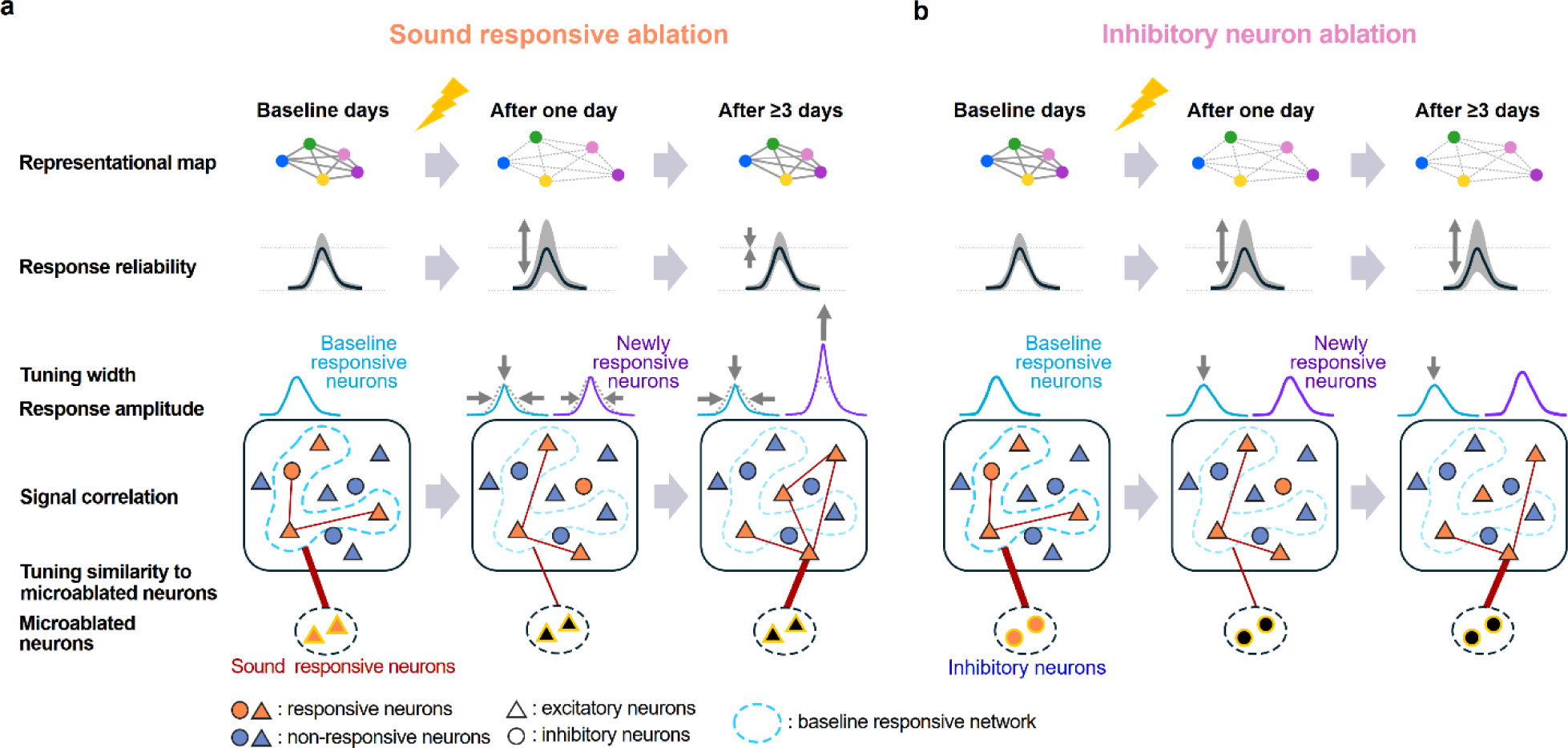
Schematic summary of homeostasis in the representational map and the underlying single-neuron mechanisms. **a.** Schematics in ablation of sound responsive neurons. After microablation of sound responsive neurons, the representational map undergoes temporal disturbance and recovery in 3-5 subsequent days. Here the representational map is depicted as the colored dots corresponding to different sound identities in representational space from population activities (top row). Longer distance between dots in this space indicates lower correlation i.e., dissimilar relationship of population activities between the sounds. Response reliability across trials in individual neurons is reduced on the day after ablation shown as the example tuning curve with a shaded gray band (the 2^nd^ row). The broader band indicates a larger variability in responses across trials. The width of tuning curve in the individual neurons is reduced on the day after ablation and remains narrowed over several days (the 3^rd^ row). The narrowing of the width is more pronounced in baseline responsive neurons (the 3^rd^ row, light blue curve). The average best response amplitude for the baseline responsive neurons keeps reduced after microablation (the 3^rd^ row, light blue curve), while the best amplitude for the newly responsive neurons, which are unresponsive on the day before ablation but become responsive on any other day, increased a few days after microablation (the 3^rd^ row, purple curve). The rate of response turnover, where some responsive neurons lose their responsiveness and other unresponsive neurons gain their responsiveness, is accelerated after microablation. The more neurons in the baseline responsive network (rounded squares in the 4^th^ row, light blue dotted circle) lose their responsiveness, the more neurons outside the baseline network gain responsiveness. The fraction of neuron pairs with high signal correlation, i.e., highly similar tunings increase later days after microablation (the number of red edges in the population inside the rounded squares in the 4^th^ row). The increase of fraction is specifically driven by the newly responsive neurons, which are located outside of the baseline responsive network. Similarity in tuning between the microablated neurons and the spared neurons increased later days after microablation, especially between the microablated neurons and the newly responsive neurons (bottom, shown as the thickness of red edges between these neurons). **b**. Same as **a**, but in ablation of inhibitory neurons. Different from microablation of sound responsive neurons, microablation of inhibitory neurons induces long-lasting disturbance of the representational map, which is depicted as the more widely distributed color dots in representational space (top row). The single-neuron response reliability across trials maintains a lower level than the baseline level (the 2^nd^ row) with the broader shaded gray band. Since the reduction of similarity matrix is mostly due to the destabilization of response across trials, the tuning width (the 3^rd^ row), best response amplitude (the 3^rd^ row) and tuning similarity between neurons (rounded squares in the 4^th^ row) do not change after microablation. Interestingly, the tuning similarity between the microablated neurons (bottom, dotted circle) and the newly responsive neurons increases also for inhibitory neuron ablation.

