## Extended data figures for "Homeostasis of a representational map in the neocortex"

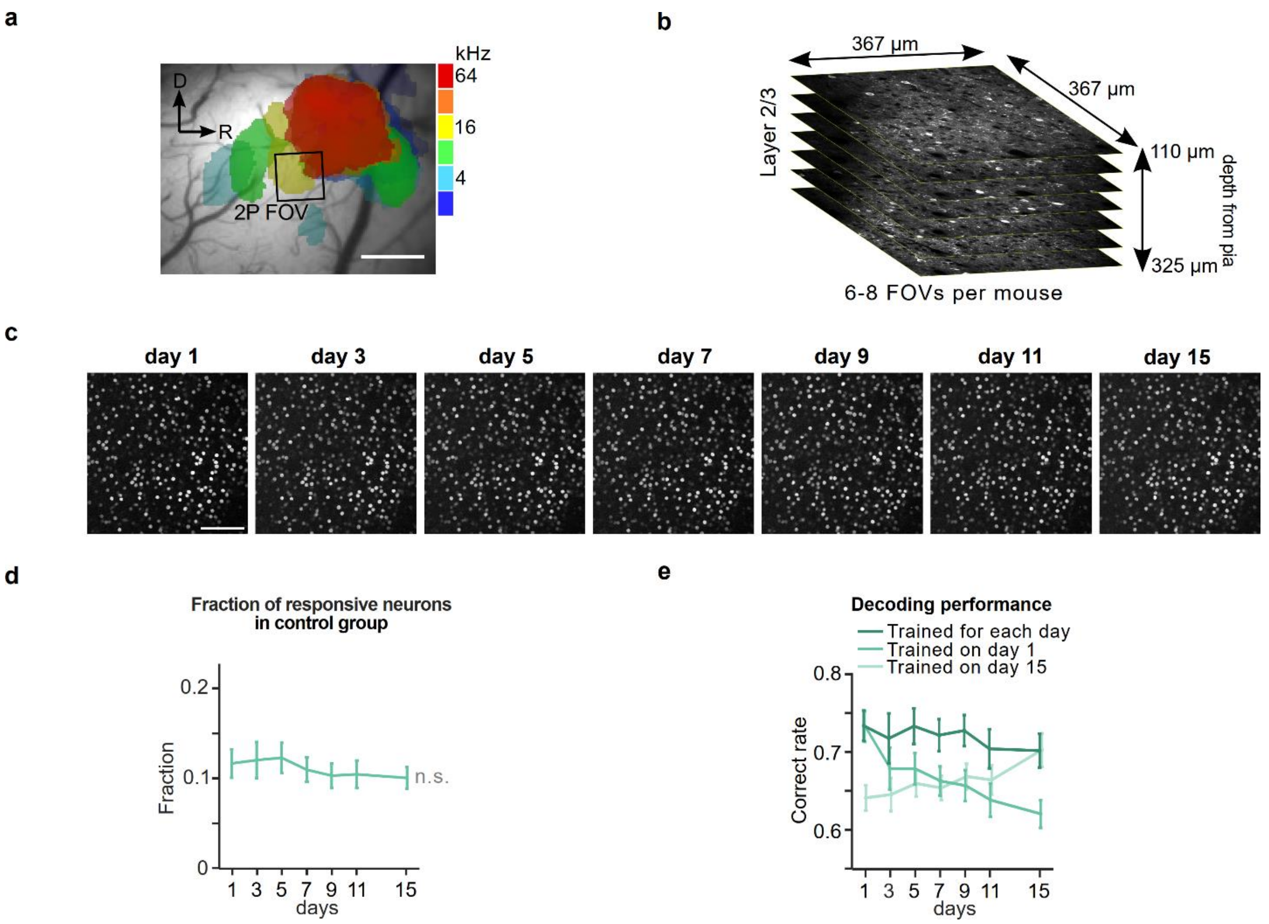


**Extended Data Fig. 1** Longitudinal imaging of sound-evoked activity in the mouse auditory cortex.

**a**. Example image of intrinsic signals on top of brain surface during burst of pure-tone (2 - 64 kHz) presentation. Intrinsic signals were recorded to functionally identify auditory fields, within which regions for two-photon calcium imaging were subsequently set. In this example, a region for two-photon imaging was selected as a black square. Scale bar: 500 µm. **b**. Schematics of sequential imaging across layer 2/3 column in the auditory cortex. 6 - 8 FOVs were imaged per mouse with average z distance of 24.2 µm (± 5.04 standard deviation). **c**. In vivo two-photon images of H2B::mCherry signal of an example FOV on all the seven imaging days. The distinct labeling in the red channel allows high-fidelity tracking of individual neurons using a signal that is independent of neuronal activity. Scale bar: 100 µm. **d**. Fraction of responsive neurons across days in control group without filtering high and low category neurons. **e**. Linear pairwise discriminability calculated by support vector machine averaged across all possible sound pairs and FOVs (mean ± s.e.m.) plotted across days. The classifier was trained with data from either given (dark green), first (green), or last (light green) imaging day.

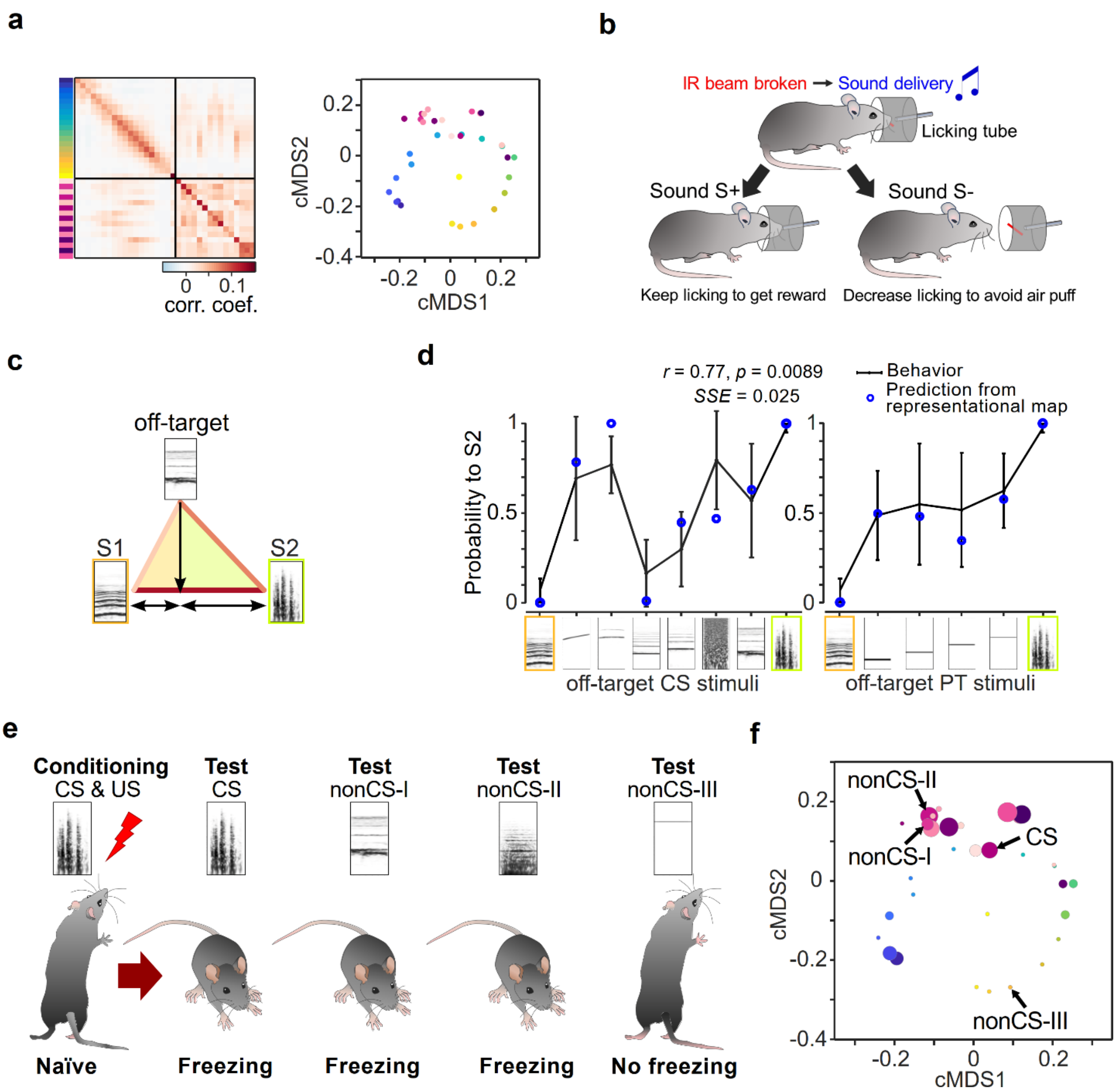


**Extended Data Fig. 2** The structure of the representational map in the auditory cortex predicts stimulus generalization in a go/no-go task as well as in classical conditioning.

**a**. Grand averaged representational similarity matrix and corresponding dimension-reduced classical Multidimensional scaling (cMDS) display constructed from the dataset acquired in the current study during baseline (n = 29 mice, 3 imaging days). **b**. Schematic of a go/no-go sound discrimination paradigm from a previous study[11], in which the same set of sound stimuli was used as in the current study. In well-trained mice licking behavior was measured in response to reinforced S+ and S- stimuli. In addition, spontaneous behavioral categorization of non-reinforced off-target stimuli was used as an estimate of perceptual similarity of a given off-target in relation to the pair of target stimuli. **c**. Schematic of the application of the neurometric similarity matrix to estimate perceptual distances. The length of each side in the triangle of off-target, and reinforced stimuli S1 and S2 is determined by the correlation coefficients. The estimated relative representational similarity of the off-target stimulus to S1 or S2 is defined as the internal dividing point in the line connecting S1 and S2, orthogonally mapped from the off-target point. **d**. Solid black line: Behaviorally evaluated perceptual similarity as the go probability of off-target sounds in go no-go behavioral task (Adapted from Bathellier et al., 2012). Blue dots: Prediction of perceptual similarity from the representational similarity matrix in **a** with the metric in **b**. **e**. Stimulus generalization to conditioned and non-conditioned sound stimuli from another previously published study[12], in which the same set of 34 sound stimuli as in the current study was used. One of the stimuli was used as a conditioned stimulus (CS) in a classical auditory-cued fear conditioning paradigm and was paired with a foot shock (US). Four days after conditioning, a generalization test was performed by presenting the CS and the three nonCS sounds (I, II, and III), without foot shock. The conditioned mice exhibited freezing behavior to CS, nonCS-I nonCS-II, but not to nonCS-III. **f**. Dimension-reduced display of the representational map obtained in the current study shown in **a**. Dot diameters depict a measure of neuronal plasticity (increased likelihood of a given sound stimulus to activate the same neuron assembly as the CS) obtained in the previous study[12]. Note, that the proximity between nonCS stimuli and the CS in the representational map allows us to predict the degree of plasticity in sound responses in the auditory cortex as well as the level of behaviorally measured stimulus generalization.

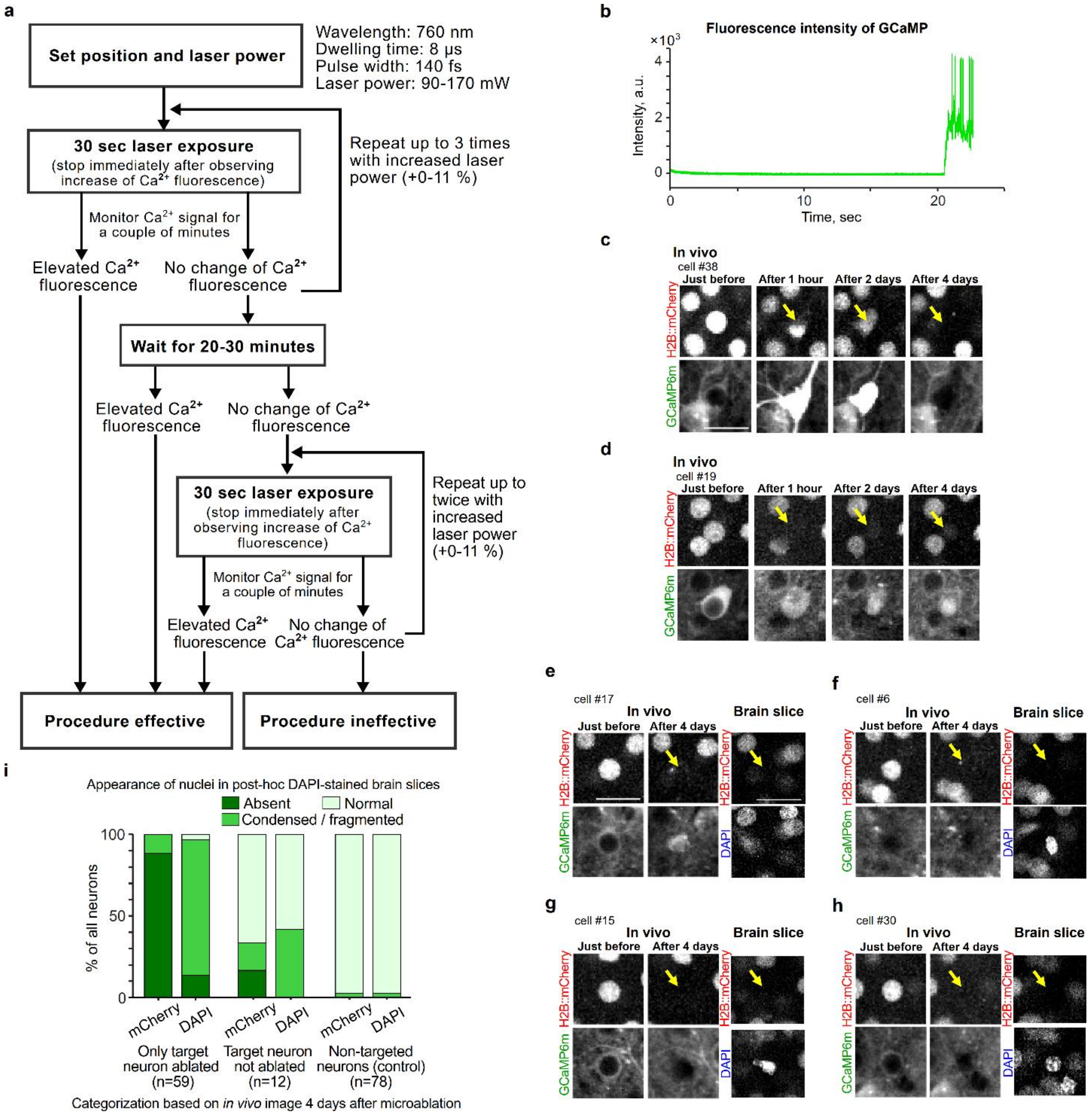


**Extended Data Fig. 3** Protocol and validation of the microablation procedure.

**a.** Protocol for microablation. See also Methods for further details. **b**. Example trace of Calcium fluorescence during laser exposure on a target neuron when doing point scanning. $Ca^{2+}$ fluorescence was monitored while exposing the laser, and the time-lapse imaging was continued for a couple of seconds to a few tens of second until we observed abrupt elevation of the signal. In this example, abrupt increase of fluorescence intensity was observed around 21 sec from the onset of laser exposure. **c-d**. Examples of targeted neurons imaged in vivo over the time course of up to 4 days. In **c**, the targeted neuron showed a permanently elevated fluorescence signal one hour after the microablation procedure. After 4 days, both the nucleus fluorescence and the calcium fluorescence were undetectable. In another example targeted neuron in **d**, after 4 days of the microablation procedure, filled calcium fluorescence remained while the nucleus fluorescence was almost eliminated. When target neurons are successfully microablated, in most cases both signals disappeared (**c**, 89.4 %), in rare cases nucleus signal was lost with filled $Ca^{2+}$ fluorescence (**d**, 10.6 %). **e-h**. Evaluation of the effectiveness of the procedure by re-identifying individual targeted neurons in fixed brain sections counter stained by 4’,6-diamidino-2-phenylindole (DAPI). Four days after the microablation procedure, neurons which lost the H2B::mCherry signal exhibited an absent DAPI signal (**e**) or an abnormally fragmented DAPI morphology (**f**-**h**). **i**. Summary bar plots of nucleus fluorescence signals and post-hoc DAPI signals in targeted neurons categorized based on in vivo images 4 days after the microablation procedure.

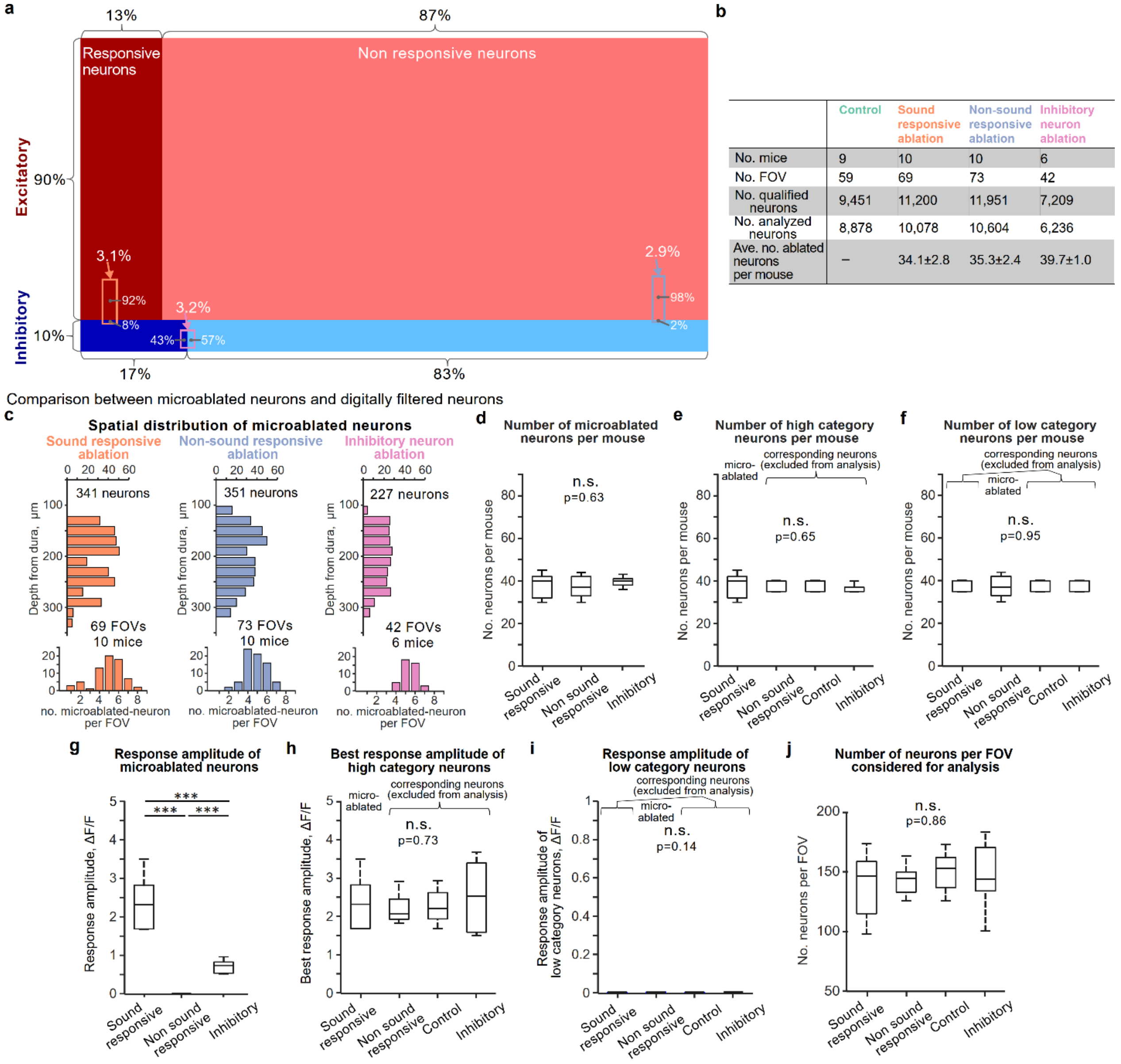


| | Control | Sound responsive ablation | Non-sound responsive ablation | Inhibitory neuron ablation |
|---|---|---|---|---|
| No. mice | 9 | 10 | 10 | 6 |
| No. FOV | 59 | 69 | 73 | 42 |
| No. qualified neurons | 9,451 | 11,200 | 11,951 | 7,209 |
| No. analyzed neurons | 8,878 | 10,078 | 10,604 | 6,236 |
| Ave. no. ablated neurons per mouse | – | 34.1±2.8 | 35.3±2.4 | 39.7±1.0 |

**Extended Data Fig. 4** Response statistics of the microablated neurons.

**a**. Summary schematic indicating the average fraction of neurons with their categorization of responsiveness and different cell types. For a subset of mice in each experimental group (5/10, 5/10, 7/9 mice in sound responsive cohort, non-sound responsive cohort, control cohort, respectively), we also labeled inhibitory cortical neurons by co-injecting an AAV vector encoding the mDlx enhancer system (AAV-mDlx-NLS-tagBFP) with the AAV vectors expressing GCaMP6m and H2B::mCherry (see also Fig. 5 and Extended Data Fig. 8). Therefore, we were able to identify the excitatory and inhibitory types of microablated and spared neurons. Small rectangles indicate the fraction of microablated neurons for each experimental group. The length of horizontal side refers to the fraction of microablated neurons out of all qualified neurons per FOV. The fraction along the vertical side refers to the ratio of excitatory and inhibitory neurons in the microablated neurons. The categorization of responsiveness is also given for an additional experimental cohort in which inhibitory neurons were microablated (see also Fig. 6). Orange: sound responsive ablation; Blue: non-sound responsive ablation; Pink: inhibitory neuron ablation. The fraction along the horizontal side in the pink rectangle refers to the ratio of significantly responsive and unresponsive neurons in microablated inhibitory neurons per FOV. **b**. Summary table of number of mice, number of FOV, number of neurons and number of averaged microablated neurons per FOV acquired for each experimental group. For sound responsive ablation and non-sound responsive ablation, 41.5 ± 1.75 neurons and 40.3 ± 2.40 neurons were targeted per mouse on average, meaning that 81.9 ± 5.5% and 87.5 ± 2.4% were successfully ablated out of the total number of target neurons respectively, based on the criteria described in Methods. **c**. Spatial distribution of microablated neurons for each experimental group. Top: distribution along depth from dura. Bottom: Distribution of number of microablated neurons per FOV. **d**. Box plots of number of microablated neurons per mouse across experimental groups. One-way ANOVA test, $F(2, 23) = 0.47$, $p = 0.63$. **e**. Number of high category neurons per mouse, which means number of microablated neurons for sound responsive ablation and number of corresponding neurons for the other groups. One-way ANOVA test, $F(3, 30) = 0.56$, $p = 0.648$. **f**. Number of low category neurons per mouse, which means number of microablated neurons for non-sound responsive ablation and number of corresponding neurons for the other groups. One-way ANOVA test, $F(3, 30) = 0.12$, $p = 0.95$. **g**. Response amplitude of microablated neurons averaged during baseline days (3 and 5). The amplitude of non-sound responsive neurons was defined as the average amplitude across stimuli for each neuron. t-test with FDR correction between sound responsive ablation vs. non-responsive ablation: $p < 0.001$; sound responsive ablation vs. inhibitory neuron ablation: $p < 0.001$; non-responsive ablation vs. inhibitory neuron ablation: $p < 0.001$. **h**. Best response amplitude of high category neurons during baseline days. One-way ANOVA test, $F(3, 30) = 0.44$, $p = 0.73$. **i**. Response amplitude of low category neurons during baseline days. The amplitude of non-sound responsive neurons was defined as the average amplitude across stimuli for each neuron. One-way ANOVA test, $F(3, 30) = 1.98$, $p = 0.14$. **j**. Number of spared neurons per FOV considered for analysis. One-way ANOVA test, $F(3, 31) = 0.25$, $p = 0.86$.

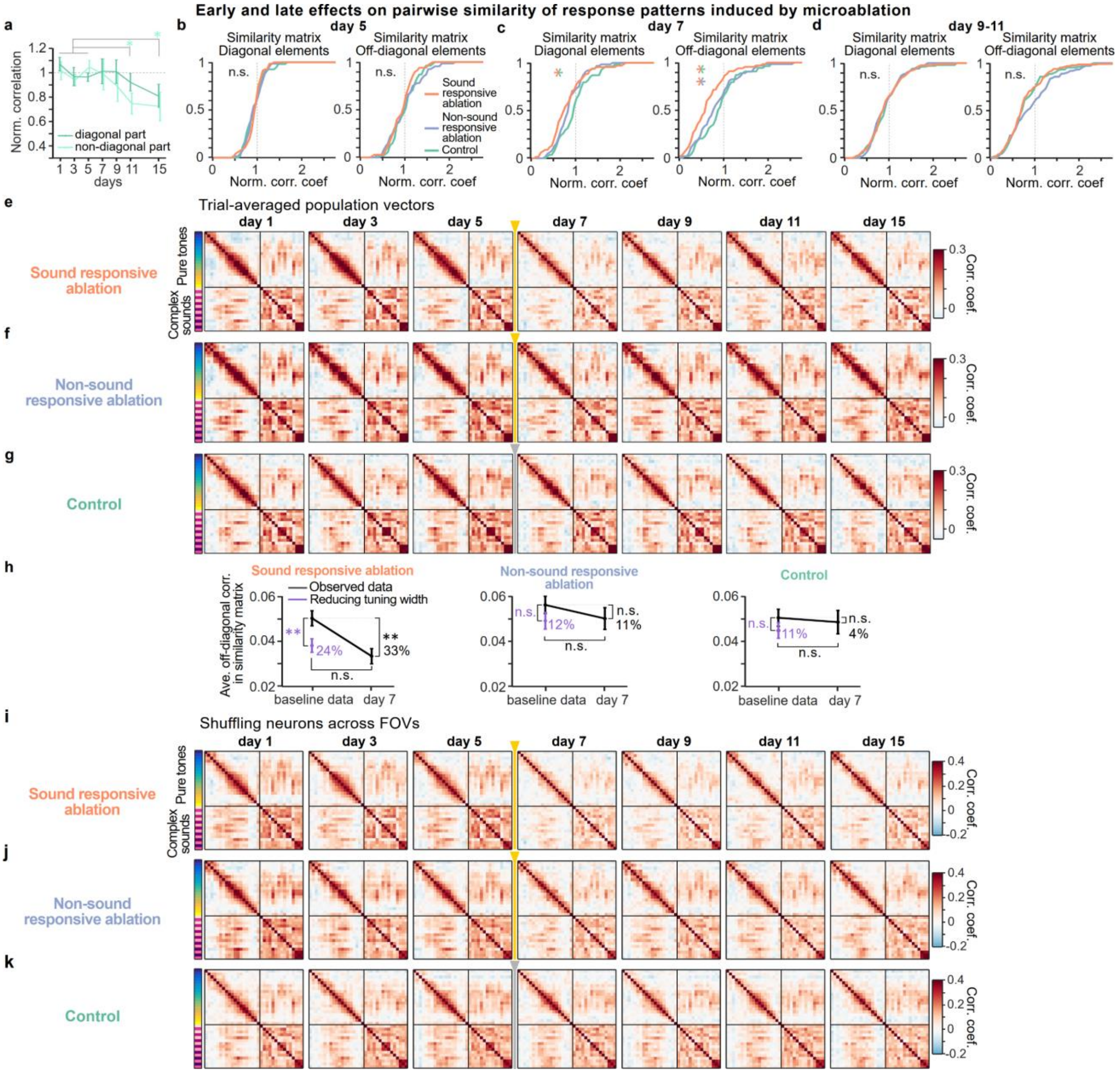


**Extended Data Fig. 5** Additional analyses of the effects on the representational maps induced by microablation.

**a**. Normalized correlations averaged across diagonal elements (dark green) and off-diagonal elements (light green) in the similarity matrices from the filtered data without the high and low category neurons. Two-sample t-test between baseline days vs. days after ablation with FDR correction, in average of diagonal elements: $p = 0.926$, $p = 0.926$, $p = 0.534$, $p = 0.0838$, for day 7, 9, 11, 15, respectively; average of off-diagonal elements: $p = 0.882$, $p = 0.277$, $p = 0.0072$, $p = 0.0072$ for day 7, 9, 11, 15, respectively. **b**. Cumulative distribution of baseline normalized correlation coefficient along diagonal elements (left) and along off-diagonal elements (right) on day 5 in the representational map for each experimental group. To make sure that the distributions of normalized correlation in the representational map during baseline were comparable between the experimental groups, difference in these cumulative distributions was tested by Dunn's test: $Q_{critical} = 2.39$, all Q values between groups for both diagonal and off-diagonal components in the similarity matrices, $Q < Q_{critical}$. Mann-Whitney *U* test between groups with FDR correction for both the diagonal and the off-diagonal components, all p values, $p > 0.5$. **c**. Cumulative distribution of baseline normalized correlation coefficient on day 7 in the representational map. Difference in these cumulative distributions between the experimental groups was again tested by Dunn's test as well as Mann-Whitney *U* test. The cumulative distribution in sound responsive ablation cohort significantly relatively shifted to lower correlation than the other groups: For diagonal components, $Q_{critical} = 2.39$; Sound responsive ablation vs. Control: Q = 2.93; Sound responsive ablation vs. Non-responsive ablation: $Q < Q_{critical}$; Non-responsive ablation vs. Control: $Q < Q_{critical}$; Mann-Whitney *U* test with FDR correction between Sound responsive ablation vs. Control: p=0.016; Sound responsive ablation vs. Non-responsive ablation: p = 0.077; Non-responsive ablation vs. Control: p = 0.20; For off-diagonal components, Sound responsive ablation vs. Control: Q = 3.44; Sound responsive ablation vs. Non-responsive ablation: Q = 2.57; Non-responsive ablation vs. Control: $Q < Q_{critical}$; Mann-Whitney *U* test with FDR correction between Sound responsive ablation vs. Control: p = 0.0018; Sound responsive ablation vs. Non-responsive ablation: p = 0.015; Non-responsive ablation vs. Control: p = 0.33. **d**. Cumulative distribution of baseline normalized correlation coefficient during day 9-11 in the map. Dunn's test: $Q_{critical} = 2.39$, all Q values between groups for both diagonal and off-diagonal components in the similarity matrices, $Q < Q_{critical}$. Mann-Whitney *U* test between groups with FDR correction for both the diagonal and the off-diagonal components, all p values, p > 0.08. **e-g**. Similarity matrices from trial-averaged population response vectors across days in sound responsive ablation condition, non-sound responsive ablation condition and control, respectively. **h**. Simulated off-diagonal correlation of similarity matrix by experimentally derived downscaled tuning curve in individual neurons. Two-sample t test between baseline correlation vs. simulated correlation after downscaling, p = 0.0083 for sound responsive cohort; p = 0.190 for non-sound responsive cohort; p = 0.270 for control; Two-sample t test between simulated correlation after downscaling vs. correlation on day 7: p = 0.419, p = 0.888, p= 0.575 for sound responsive, non-sound responsive, and control cohort, respectively. **i-k**. Similarity matrices across days from population response vectors after shuffling neurons across FOVs for each experimental cohort.

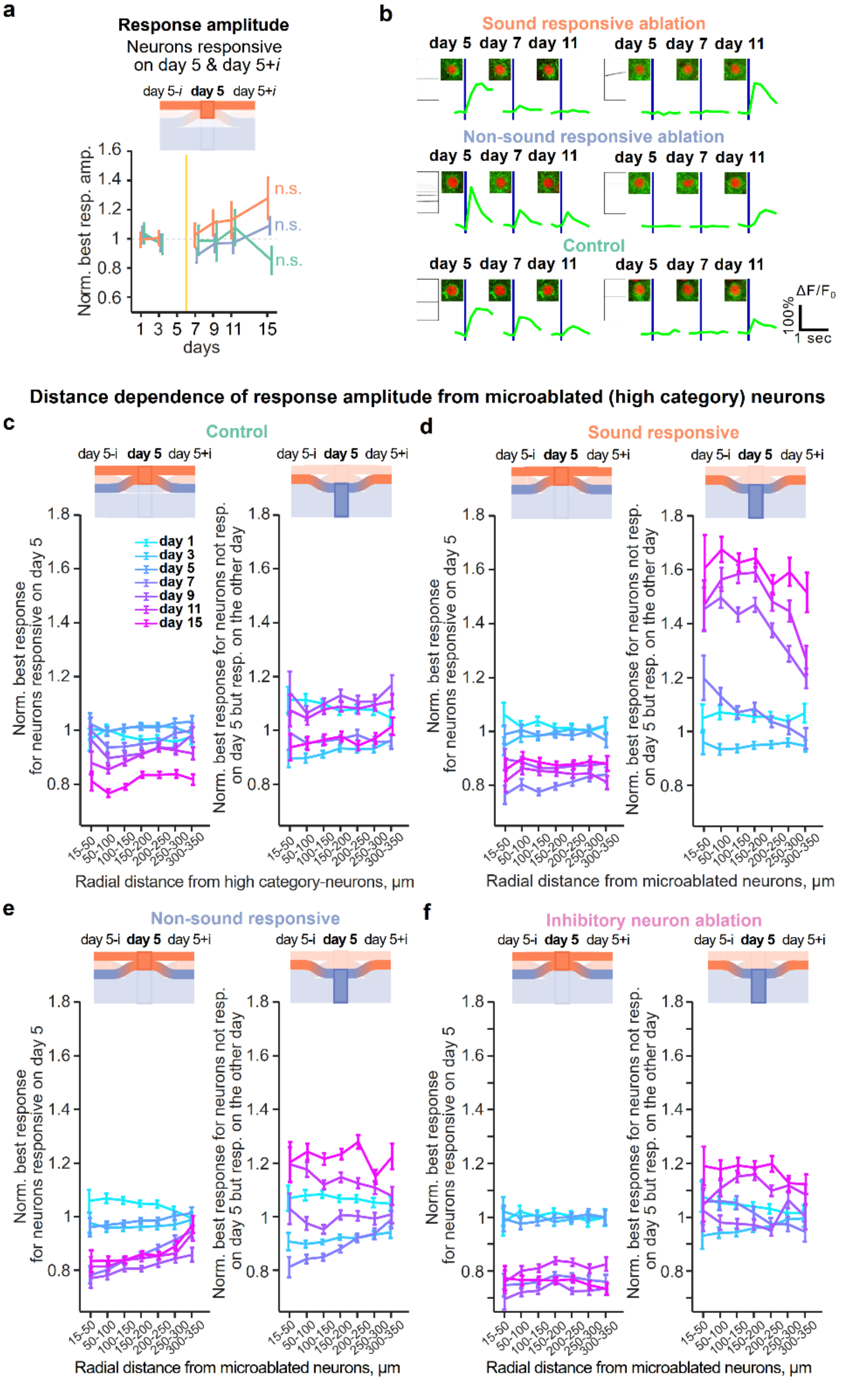


**Extended Data Fig. 6** Additional analyses of the microablation-induced effects on response amplitudes.

**a.** Best response amplitude of neurons responsive on both day 5 and day 5±i normalized by the average best response during early baseline days (1 and 3). One-way ANOVA test, sound responsive ablation: $F(5, 54) = 1.29$, $p = 0.28$; Non-responsive ablation: $F(5, 54) = 1.35$, $p = 0.26$; Control: $F(5, 48) = 0.58$, $p = 0.72$; **b.** Exemplary sound-evoked $Ca^{2+}$ transients of neurons which responded to sounds on day 5 and reduced the amplitudes in these responses in following days after ablation in sound responsive ablation (top left), non-sound responsive ablation (middle left), and control (bottom left), respectively. Right: Exemplary sound-evoked $Ca^{2+}$ transients of neurons unresponsive on day 5, but becoming responsive in the following days after ablation. **c**. Assessing the spatial extent of the microablation-induced effect on normalized best response amplitudes in the spared neurons. In the control group, the high category neurons with strong sound-evoked responses that were excluded from the further analysis were considered as reference (see Methods). Spared neurons were split into neurons responsive on day 5 (Left) and newly responsive neurons (Right). In a given mouse, the three-dimensional distance of a spared neuron to each of the 30-40 microablated neurons (or high category neurons in the control group) was calculated, i.e. contributed multiple distance measurements associated with the same effect on the best response amplitude. All distance combinations were categorized into distance bins ranging between 15 and 350 µm. For each distance bin, the corresponding best response amplitude in each neuron was normalized to the average baseline amplitude for each mouse, the normalized best response was averaged across neurons (error bars are shown as s.e.m.). In the control group, normalized best responses of neurons responsive on day 5 gradually decreased over days with generally constant amplitude across distance, but exhibited modest reduction of normalized amplitude on later (day 11 and 15) days (Two-sample t-test between short and long distance, day 1 to day 15: p = 0.129, p = 0.823, p = 0.138, p = 0.0486, p = 0.0612, p = 0.0014, p = 0.0012). Normalized best responses of newly responsive neurons remained rather flat across distance and did not show any systematic change during and after baseline days (Two-sample t-test between short and long distance, day 1, p = 0.0656; day 3, p = 0.014; day 7, p = 0.957; day 9, p = 0.127; day 11, p = 0.0797; day 15, p = 0.194). **d**-**f**. Same as **c**, but from microablated neurons for sound responsive ablation, for non-responsive ablation, and for inhibitory neuron ablation, respectively. In sound responsive ablation, normalized best responses of neurons responsive on day 5 decreased over days, but with constant amplitude across distance (Two-way ANOVA, $F(6, 244429) = 0.451$, $p = 0.845$ across distance bins; $F(6, 244429) = 151.58$, $p < 1.0\times10^{-10}$ across days). On the other hand, normalized best responses of newly responsive neurons strongly increased the amplitudes later days (Two-way ANOVA, $F(6, 121590) = 10.9$, $p = 3.56\times10^{-12}$ across distance bins; $F(5, 121590) = 543.19$, $p < 1.0\times10^{-12}$ across days). Three days after microablation, the normalized amplitude with nearby distance exhibited larger amplitude than the amplitude distant from ablated neurons (Two-sample t-test between short (< 100 µm) and long distance (250-350 µm), day 1, p = 0.488; day 3, p = 0.419; day 7, p = 0.0001; day 9, $p < 1.0\times10^{-4}$; day 11, p = 0.0002; day 15, p = 0.124). For non-sound responsive ablation, normalized best responses of neurons responsive on day 5 decreased over days and the reduction was more prominent at shorter distance from ablated neurons (Two-way ANOVA, $F(6, 284429) = 9.47$, $p = 1.99\times10^{-10}$ across distance bins; $F(6, 284429) = 230.36$, $p < 1.0\times10^{-10}$ across days; Two-sample t-test between short and long distance, day 1, p = 0.0116; day 3, p = 0.495; day 5, p = 0.0313; day 7, $p < 1.0\times10^{-4}$; day 9, p = 0.0005; day 11, p = 0.0003; day 15, p = 0.0004). On the other hand, normalized best responses of newly responsive neurons did not change over distance, but showed slight increase in amplitude later days after ablation (Two-way ANOVA, $F(6, 146974) = 1.85$, $p = 0.0853$ across distance bins; $F(5, 146974) = 274.42$, $p = 1.0\times10^{-10}$ across days). The normalized amplitude on day 7 was suppressed at shorter distance, but conversely increased later days (Two-sample t-test between short and long distance: day 1, p = 0.26; day 3, p = 0.0359; day 7, $p < 1.0\times10^{-5}$; day 9, p = 0.63; day 11, p = 0.0122; day 15, p = 0.0846).

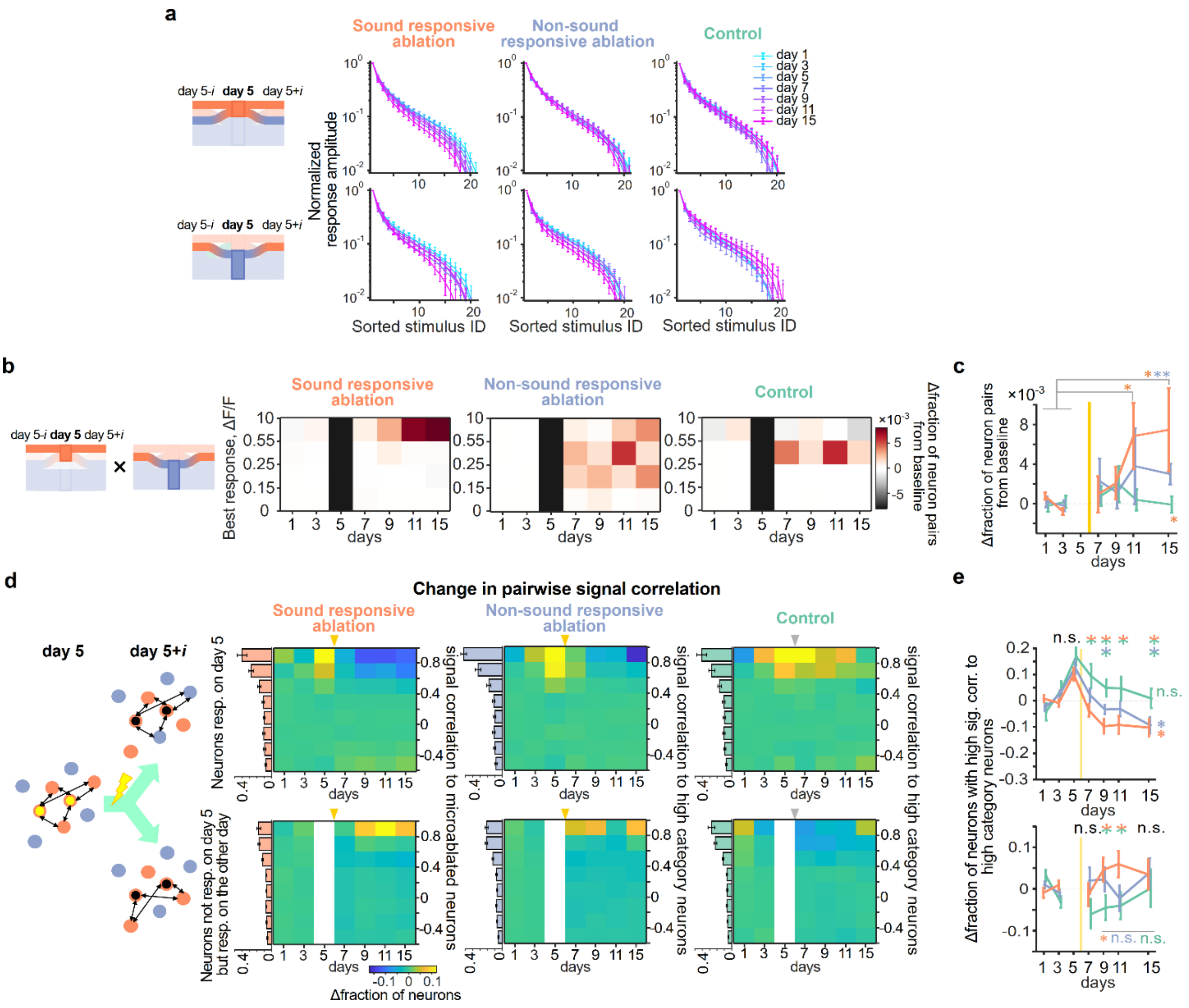


**Extended Data Fig. 7** Additional analyses of single-neuron response dynamics contributing to the change of the representational maps.

**a**. Same as Fig. 3**e** bottom, but normalized tuning curves of day 5 responsive neurons (top) and of neurons unresponsive on day 5 but responsive on the other day (bottom), for sound responsive cohort (left), non-sound responsive cohort (middle) and control cohort (right), respectively. **b**. Same as Fig. 4**c**, but the colormaps displaying the change in the fraction of high-signal-correlation neuron pairs, one of which was from neurons responsive on both day 5 and any other day, and the other of which was from neurons unresponsive on day 5 but responsive on any other day. **c**. Change in the fraction of neuron pairs with high signal correlation from baseline at the largest response amplitude bin in the corresponding colormaps in **b**. Two-sample t-test between baseline days and days after ablation with FDR correction: $p = 0.243$, $p = 0.0667$, $p = 0.0146$, $p = 0.0175$ for day 7, 9, 11, 15 in sound responsive cohort; $p = 0.120$, $p = 0.188$, $p = 0.120$, $p = 0.0055$ for day 7, 9, 11, 15 in non-sound responsive cohort; $p > 0.398$ for all post-sham ablation days in control cohort. Permutation test for group comparison: $p > 0.05$ from day 7 - 11 for all the three cohorts, but $p < 0.05$ on day 15 for sound responsive cohort. **d**. Left: Scheme of signal correlation between microablated neurons and spared neurons, pairs of which are defined based on the responsiveness on day 5 and day 5±i. Right: Colormaps show baseline-subtracted fraction of neurons with signal correlation between microablated neurons (or high category neurons except for the sound responsive cohort) and spared neurons, which are responsive on day 5 (top) and not responsive on day 5 but responsive on the other day (bottom). Bar plots next to the colormap are fractions of neurons for signal correlation bins averaged across baseline days (day 1 and day 3) in each split group. **e**. Baseline normalized change in fraction of spared neurons with high signal correlation toward high category neurons. One-way ANOVA, sound responsive ablation: $F(5,114) = 3.58$, $p = 0.0048$; non-responsive ablation: $F(5,114) = 3.7$, $p = 0.0039$), the fraction did not reduce at all for control group (One-way ANOVA, $F(5,101) = 1.44$, $p = 0.21$, comparison across groups by two-sample t-test with FDR correction between responsive ablation vs. control on day 7, $p = 0.042$; day 9, $p = 0.0051$; day 11, $p = 0.049$; day 15, $p = 0.061$; responsive ablation vs. non responsive ablation on day 7, $p = 0.15$; day 9, $p = 0.057$; day 11, $p = 0.18$; day 15, $p = 0.85$; non-responsive ablation vs. control on day 7, $p = 0.16$; day 9, $p = 0.047$; day 11, $p=0.18$; day 15, $p = 0.061$; For newly responsive neurons, comparison across groups by one-sample t-test with FDR correction between responsive ablation vs. control on day 7, $p = 0.21$; day 9, $p = 0.055$; day 11, $p = 0.049$; day 15, $p = 0.43$; responsive ablation vs. non-responsive ablation on day 7, $p = 0.21$; day 9, $p = 0.29$; day 11, $p = 0.054$; day 15, $p = 0.47$; non-responsive ablation vs. control on day 7, $p = 0.15$; day 9, $p = 0.086$; day 11, $p = 0.35$; day 15, $p = 0.43$.

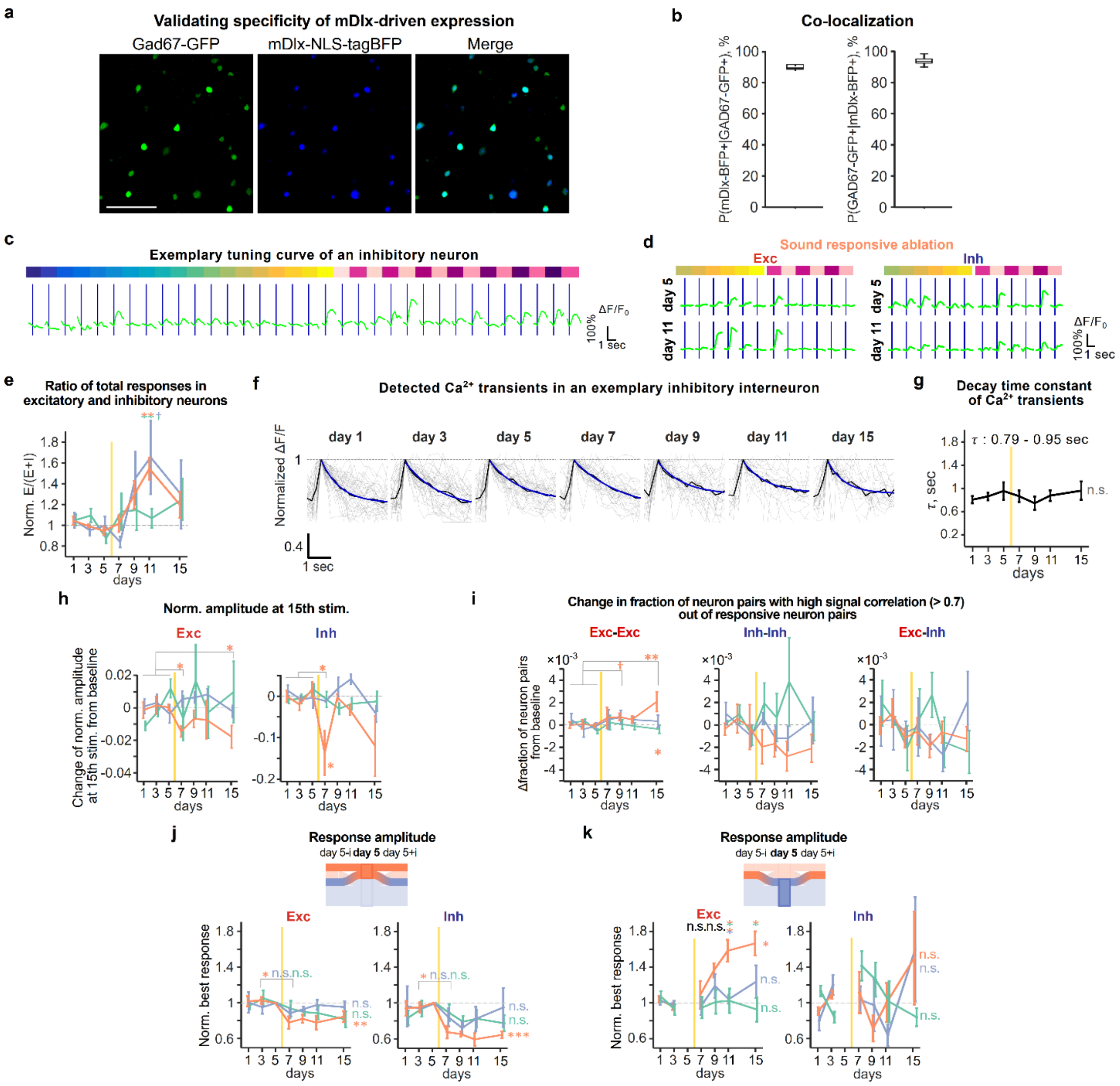


**Extended Data Fig. 8** Validation of the viral construct labeling inhibitory neurons and additional analyses of the microablation-induced effects on excitatory and inhibitory neurons.

**a**. Representative example of colocalization between mDlx-driven tagBFP expression and Gad67. Scale: 100 μm. **b**. Quantification of colocalization. The fraction of interneuron labeled with the blue fluorescent protein marker out of the entire quality qualified neurons was around 10% for each group (10.5 ± 0.5% (mean ± s.e.m. across mice) in sound responsive ablation, 9.8 ± 0.6% in non-sound responsive ablation, and 7.8 ± 0.4% in control, respectively), in line with previous works with auditory cortex in mature adult mice[82]. When we revisited the profile of cell types in microablated neurons, nearly all the ablated neurons were excitatory neurons (92.3 ± 2.0% for sound-responsive ablated neurons, 98.4 ± 0.9% for non-sound responsive ablated neurons), the fraction of which is higher than the average fraction of excitatory neurons in the population. **c**. Exemplary tuning curve of inhibitory neuron. **d**. Representative excitatory (left) and inhibitory (right) neurons, which change response amplitude to sound stimuli from day 5 to day 11, in sound responsive ablation. **e**. Baseline-normalized ratio of the total sound-evoked activity in excitatory and inhibitory neurons (see Methods). Data presented as mean ± s.e.m. across mice. Two-sided t-test with FDR adjusted p-values, sound responsive cohort vs. control, non-sound responsive cohort vs. control and sound responsive cohort vs. non-sound responsive cohort on day 7: p > 0.5 for all combinations; day 9: p = 0.039, p = 0.039, p = 0.16; day 11: p = 0.045, p = 0.032, p = 0.27; day 15: p > 0.5 for all combinations, respectively; the normalized ratio of best amplitudes on day 15, 1.09 ± 0.088 for sound responsive ablation, 1.10 ± 0.11 for non-sound responsive ablation. **f**. Traces of detected calcium transients of an inhibitory neuron over days. We verified that the changes of response amplitude in inhibitory neurons after microablation (Fig. 5**e**) were not due to another feature like $Ca^{2+}$ kinetics in somas of inhibitory neurons characterized by the decay time constant[83]. **g**. Decay time constant of calcium transients of all identified interneurons over days across mice (mean ± s.e.m.). One-way ANOVA test, F(6, 28) = 0.48; p = 0.82. **h**. Same as Fig. 3f, but change of tuning width by using normalized response amplitudes at 15th largest stimulus index in the tuning curve over days for excitatory (left) and inhibitory (right) neurons. Two-sample t-test of Δnormalized amplitude for excitatory neurons between baseline days vs. days after ablation with FDR correction. Sound responsive cohort: p = 0.0106, p = 0.253, p = 0.346, p = 0.0106, for day 7, 9, 11, 15, respectively; non-sound responsive cohort: p > 0.474 for all post-ablation days; Control: p > 0.67 for all post-ablation days. For inhibitory neurons, sound responsive cohort: p = 0.213, p = 0.88, p = 0.330, p = 0.0717 for day 7, 9, 11, 15, respectively; non-sound responsive cohort: p > 0.20 for all post-ablation days; Control: p > 0.24 for all post-ablation days. **i**. Change in fraction of neuron pairs with high signal correlation and with large response amplitude from baseline among excitatory-excitatory neurons (left), inhibitory-inhibitory neurons (middle), and excitatory-inhibitory neurons (right). Two-sample t-test of Δfraction of excitatory-excitatory neuron pairs with FDR correction in sound responsive cohort: p = 0.15, p = 0.051, p = 0.11, p = 0.0034 for day 7, 9, 11, 15; non-sound responsive cohort: p > 0.29 for all post-ablation days; control: p > 0.43 for all post-ablation days. Permutation test for group comparison: p > 0.05 for day 7, 9, 11, p < 0.025 on day 15 in sound responsive cohort; p > 0.05 for all post-ablation days in both non-sound responsive and control cohorts. For inhibitory-inhibitory neuron pairs, sound responsive cohort: p > 0.15 for all post-ablation days; non-sound responsive cohort: p > 0.28 for all post-ablation days; control: p > 0.16 for all post-ablation days. For excitatory-inhibitory neuron pairs, sound responsive cohort: p > 0.25 for all post-ablation days; non-sound responsive cohort: p > 0.30 for all post-ablation days; control: p > 0.26 for all post-ablation days. **j**. Baseline normalized best response amplitude of neurons responsive on day 5 in excitatory (left) and inhibitory (right) neurons for the three experimental groups. When splitting excitatory and inhibitory neurons based on the responsiveness on day 5, as observed in Fig. 4**b**, the normalized best response amplitude of neurons responsive on day 5 significantly decreased on day 7 for both the excitatory and inhibitory neurons in the sound responsive ablation cohort, but not in the other cohorts (One-way ANOVA across days, Excitatory neurons in sound responsive ablation, F(6, 28) = 4.21, p = 0.0038; non-responsive ablation, F(6, 28) = 1.01, p = 0.43; control, F(6, 42) = 1.36, p = 0.25; Inhibitory neurons in sound responsive ablation, F(6, 21) = 7.44, p = $2.3\times10^{-4}$; non-responsive ablation, F(6, 28) = 0.51, p = 0.80; control, F(6, 42) = 1.02, p = 0.42; Two-sample paired t-test of the normalized best amplitude of excitatory neurons between day 3 vs. day 7: sound responsive ablation, p = 0.023; non-responsive ablation, p = 0.34; control, p = 0.38; inhibitory neurons between day 3 vs. day 7: sound responsive ablation, p = 0.049; non-responsive ablation, p = 0.38; control, p = 0.59). **k**. Same as **j**, but for newly responsive neurons. For the neurons unresponsive on day 5 but being responsive on the other day, the normalized best response amplitude of excitatory neurons exhibited a delayed but substantial increase after microablation in the sound responsive ablation (One-way ANOVA across days, F(5, 24) = 7.97, p = $1.5\times10^{-4}$) like Fig. 4b right, but inhibitory neurons did not (F(5, 24) = 1.01, p = 0.43). For the non-responsive ablation and the control cohorts, both the excitatory and inhibitory neurons did not change the normalized best response amplitude over days (Excitatory neurons in non-responsive ablation: F(5, 24) = 1.12, p = 0.37; control: F(5, 36) = 0.28, p = 0.92; Inhibitory neurons in non-responsive ablation: F(5, 23) = 1.18, p = 0.35; control: F(5, 25) = 1.44, p = 0.24). Group comparison revealed that the normalized amplitude of the newly responsive excitatory neurons in sound responsive ablation was significantly larger than those in non-responsive ablation and in control late after microablation (Two-sample t-test with FDR correction, day 7 and day 9: p > 0.2 for either combination of the experimental groups; day 11: sound responsive ablation vs. non-responsive ablation, p = 0.011; sound responsive ablation vs. control, p = 0.024; non-responsive ablation vs. control, p = 0.93; day 15: sound responsive ablation vs. non-responsive ablation, p = 0.14; sound responsive ablation vs. control, p = 0.011; non-responsive ablation vs. control, p = 0.19).

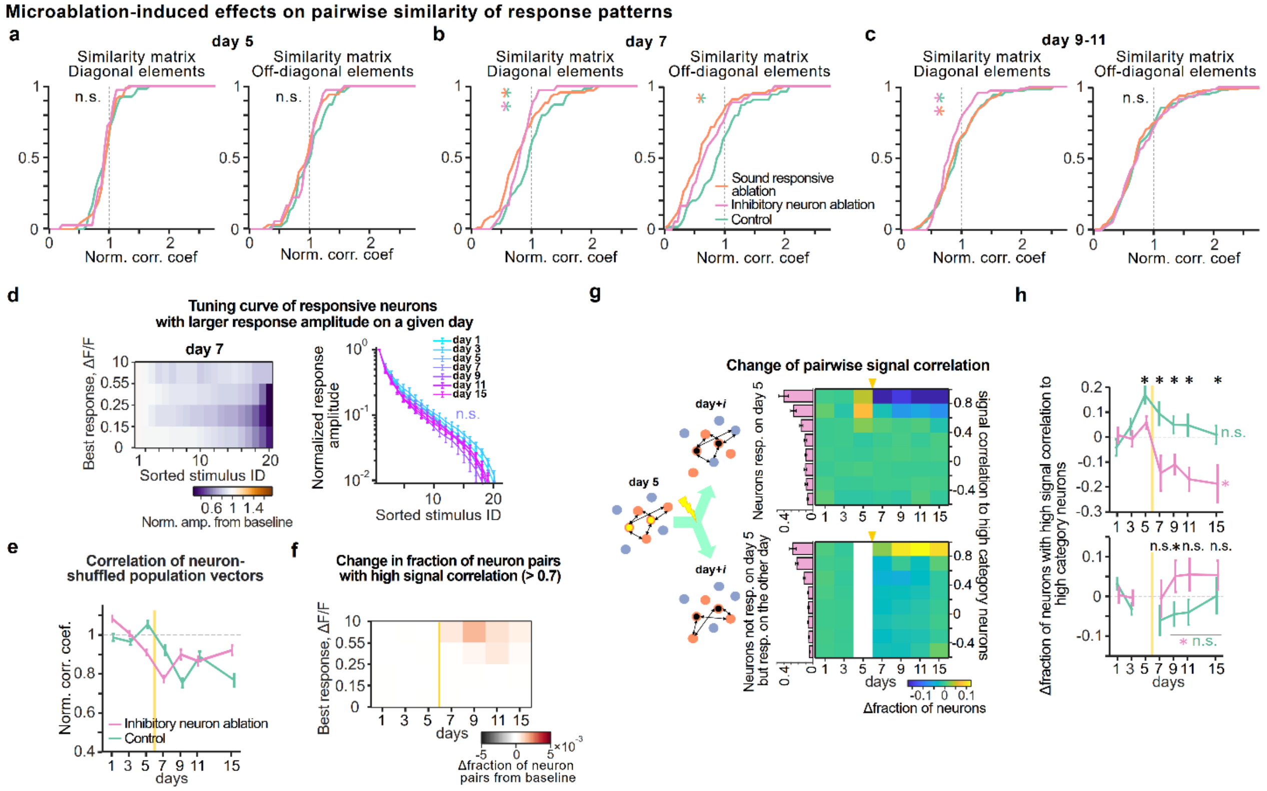


**Extended Data Fig. 9** Additional analyses of the effects induced by selective microablation of inhibitory neurons on the representational map and response characteristics of individual neurons.

**a**. Cumulative distribution of baseline normalized correlation coefficient on day 5 in the representational map for each experimental group. In inhibitory neuron ablation experiment, 91.7 ± 1.5% was successfully ablated out of the targeted neurons, based on the criteria we already described in Methods. Same as Extended Data Fig. 5a, to compare the effect of ablation between inhibitory neuron ablation and sound responsive ablation groups on the day after microabtaion, difference in these cumulative distributions was tested by Dunn's test as well as Mann-Whitney *U* test with FDR correction. These cumulative distributions on day 5 were not statistically different by definition between the groups (Dunn's test for all group combinations in both diagonal and off-diagonal elements, Q < Qcritical; Mann-Whitney *U* test, adjusted p values > 0.5). **b**. Same as **a**, but the cumulative distributions of normalized correlation on day 7. Especially for diagonal elements, the distributions in both sound responsive ablation and inhibitory neuron ablation cohort shifted toward lower correlation than control (Mann-Whitney *U* test, adjusted p-values, sound responsive ablation vs. control, p = 0.016; inhibitory vs. control, p = 0.020; sound responsive ablation vs. inhibitory, p = 0.51; Dunn's test, sound responsive ablation vs. control, Q = 3.00 (> Qcritical); inhibitory neuron ablation vs. control, Q < Qcritical). For off-diagonal elements, the distributions in sound responsive ablation cohort was lower than that in control (Mann-whitney *U* test, with adjusted p values, sound responsive ablation vs. control, p = 0.0018; inhibitory neuron ablation vs. control, p = 0.06; p = 0.29; sound responsive ablation vs. inhibitory neuron ablation; Dunn's test, sound responsive ablation vs. control, Q = 3.48 (> Qcritical); Q < Qcritical for the other group comparison). **c**. Same as **a**, but during day 9-11. On the other hand, late after microablation (day 9-11), while the distribution in sound responsive ablation went back to baseline level, the distribution in inhibitory neuron ablation, especially the diagonal elements maintained lower correlation than the other groups (Dunn's test, inhibitory neuron ablation vs control, Q = 2.44 (> Qcritical); sound responsive ablation vs. inhibitory neuron ablation, Q = 2.43 (> Qcritical); Mann-Whitney *U* test, adjusted p-values, sound responsive ablation vs. control, p = 0.87; inhibitory neuron ablation vs.

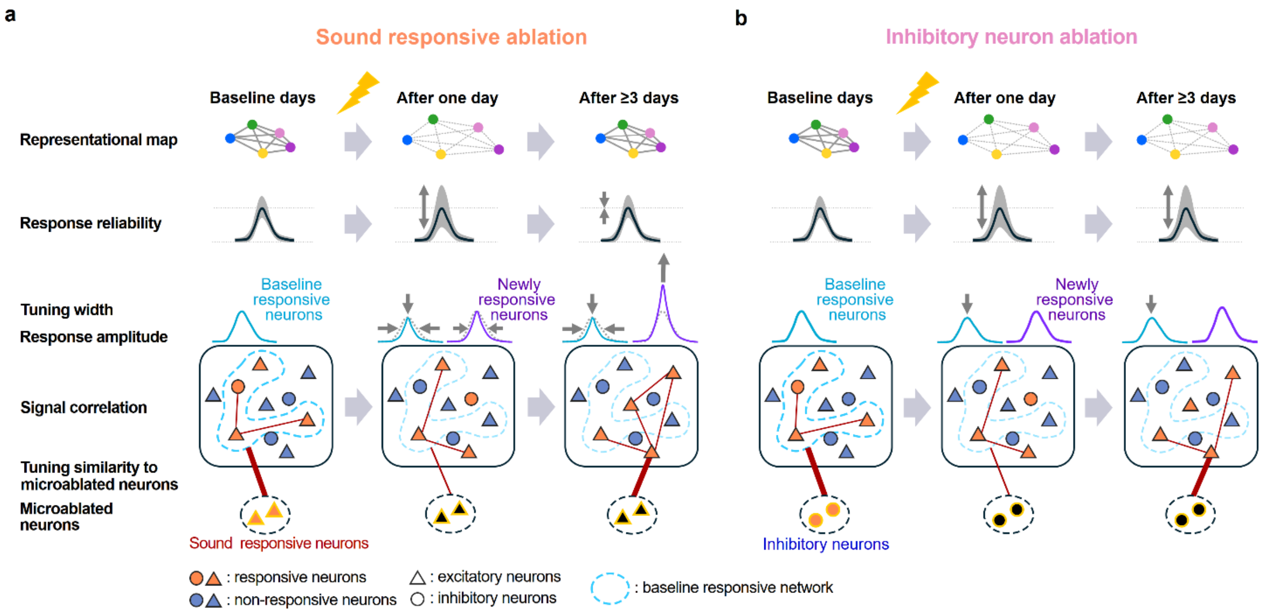


**Extended Data Fig. 10** Schematic summary of homeostasis in the representational map and the underlying single-neuron mechanisms.

**a.** Schematics in ablation of sound responsive neurons. After microablation of sound responsive neurons, the representational map undergoes temporal disturbance and recovery in 3-5 subsequent days. Here the representational map is depicted as the colored dots corresponding to different sound identities in representational space from population activities (top row). Longer distance between dots in this space indicates lower correlation i.e., dissimilar relationship of population activities between the sounds. Response reliability across trials in individual neurons is reduced on the day after ablation shown as the example tuning curve with a shaded gray band (the 2$^{nd}$ row). The broader band indicates a larger variability in responses across trials. The width of tuning curve in the individual neurons is reduced on the day after ablation and remains narrowed over several days (the 3$^{rd}$ row). The narrowing of the width is more pronounced in baseline responsive neurons (the 3$^{rd}$ row, light blue curve). The average best response amplitude for the baseline responsive neurons keeps reduced after microablation (the 3$^{rd}$ row, light blue curve), while the best amplitude for the newly responsive neurons, which are unresponsive on the day before ablation but become responsive on any other day, increased a few days after microablation (the 3$^{rd}$ row, purple curve). The rate of response turnover, where some responsive neurons lose their responsiveness and other unresponsive neurons gain their responsiveness, is accelerated after microablation. The more neurons in the baseline responsive network (rounded squares in the 4$^{th}$ row, light blue dotted circle) lose their responsiveness, the more neurons outside the baseline network gain responsiveness. The fraction of neuron pairs with high signal correlation, i.e., highly similar tunings increase later days after microablation (the number of red edges in the population inside the rounded squares in the 4$^{th}$ row). The increase of fraction is specifically driven by the newly responsive neurons, which are located outside of the baseline responsive network. Similarity in tuning between the microablated neurons and the spared neurons increased later days after microablation, especially between the microablated neurons and the newly responsive neurons (bottom, shown as the thickness of red edges between these neurons). **b**. Same as **a**, but in ablation of inhibitory neurons. Different from microablation of sound responsive neurons, microablation of inhibitory neurons induces long-lasting disturbance of the representational map, which is depicted as the more widely distributed color dots in representational space (top row). The single-neuron response reliability across trials maintains a lower level than the baseline level (the 2$^{nd}$ row) with the broader shaded gray band. Since the reduction of similarity matrix is mostly due to the destabilization of response across trials, the tuning width (the 3rd row), best response amplitude (the 3rd row) and tuning similarity between neurons (rounded squares in the 4th row) do not change after microablation. Interestingly, the tuning similarity between the microablated neurons (bottom, dotted circle) and the newly responsive neurons increases also for inhibitory neuron ablation.
